# A chromosome-level assembly of an aquatic passerine bird, the northern white-throated dipper, *Cinclus cinclus cinclus* (Linnaeus, 1758)

**DOI:** 10.64898/2026.08.20.746034

**Authors:** Marius A. Strand, Ole K. Tørresen, Morten Skage, Giada Ferrari, Ave Tooming-Klunderud, Arild Johnsen, Kjetill S. Jakobsen

**Affiliations:** Department of Biosciences, Centre for Ecological and Evolutionary Synthesis (CEES), University of Oslo, Oslo, Norway; Department of Research and Collections, Natural History Museum, University of Oslo, P.O. Box 1172 Blindern, 0318 Oslo, Norway

## Abstract

We present a chromosome-level genome assembly of a female Norwegian white-throated dipper (*Cinclus cinclus cinclus*) generated using Oxford Nanopore Technologies (ONT) long reads and Hi-C scaffolding. The assembly comprises two pseudo-haplotypes, hap1 (1186 Mb) and hap2 (1115 Mb), with 96.7% and 94.4% of sequences assigned to chromosome-scale scaffolds, respectively. Both pseudo-haplotypes contain 40 autosomes, with the Z and W sex chromosomes assigned to hap1. Compared with the PacBio HiFi-based *C. c. gularis* reference assembly bCinCin1.1.pri, which contains 38 autosomes, sequence represented as a single dot-chromosome (chr 36) is resolved into three distinct dot-chromosomes (chr 36, 39, and 40), a configuration supported by Hi-C contact patterns. BUSCO completeness was high for hap1 (99.2%) and hap2 (95.0%), with 19,003 and 17,746 predicted protein-coding genes, respectively. Compared with the HiFi-based *C. c. gularis* reference and HiFi-based assemblies generated from the same individual, the ONT-derived assemblies were substantially less fragmented and recovered more sequence from the smallest chromosomes. Synteny was otherwise largely conserved between subspecies. HiFi depletion increased strongly from macrochromosomes to micro- and dot-chromosomes, and HiFi-depleted regions were enriched for repeats and predicted non-B-DNA-associated features, particularly G-quadruplexes and direct repeats, whereas ONT coverage remained comparatively stable. These results show that conventional genome-wide assembly metrics can obscure substantial differences in the recovery of repeat-rich avian dot-chromosomes and highlight the value of chromosome-aware evaluation and ONT sequencing for recovering these regions.

## Introduction

The white-throated dipper (C*inclus cinclus*), Norway’s national bird (Holgersen, 1963), is unique among European Passeriformes because of its aquatic lifestyle characterized by the ability to dive, even walk underwater and swim using its wings to forage for insects and aquatic invertebrates (D’Amico & Hémery, 2007). It has a dark brown body with a white throat and upper breast and is about 18 cm long. The semi-aquatic lifestyle of dippers involves a suite of morphological and physiological adaptations, including dense, waterproof plumage supported by an unusually large preen gland, and specialized ocular anatomy featuring a well-developed sphincter muscle in the iris thought to facilitate vision in both air and water (Tyler & Ormerod, 1994). The species exhibits high hemoglobin rates and heart rate to accommodate the demands of active diving (Murrish, 1970; Ormerod & Tyler, 2005; Tyler & Ormerod, 1994).

The typical habitat of the white-throated dipper is fast-flowing, well-oxygenated rivers and streams. Due to its dependence on pollution-sensitive insects (mayflies, stoneflies and caddisflies) and other invertebrates as prey, its presence is correlated with water quality, and it is therefore a critical bioindicator for health of aqueous ecosystems. Several studies have shown that dipper breeding performance and population density decline in acidified or polluted waters (Tyler & Ormerod, 1994, 1992). It has also been shown that eggs and tissues accumulate different pollutants (see Morrissey et al., 2013) including plastics (see D’Souza et al., 2020). The white-throated dipper has been demonstrated as a model species in time-series studies of how wildlife adapted to cold and northern climates are affected by global warming (Nilsson et al., 2011). These human impact and climate-related impacts of the species call for further ecological, physiological and ecotoxicological studies including studies using genetic and molecular methods. The white-throated dipper has the most widespread distribution of the extant dipper species, ranging across Europe, North Africa, and Central Asia (Ormerod et al., 2020). There are many subspecies with defined geographical ranges including the northern white-throated dipper (*C.c. cinclus)* in Northern/Western Europe; the Irish dipper (*C.c. hibernicus*) in Ireland and Western Scotland and the British dipper (*C.c. gularis*) in west, central and northern England, Wales and Scotland. These abovementioned subspecies as well as the other subspecies differ in geographical distribution, migratory habits as well as small but distinct differences in body coloration (BirdLife International, 2018).

Due to its large distribution range and population size the white-throated dipper is classified as a species of “Least Concern” by the IUCN globally (BirdLife International, 2018). However, in the UK, it is currently amber-listed in the “Birds of Conservation Concern” because it has shown strong localized declines in the last 30 years (Eaton et al., 2015). Previous genetic studies have revealed a complex phylogeographic structuring across the Western Palearctic – possibly the result of several southern glacial refugia (Hourlay et al., 2008). Interestingly, a Norwegian population of *C. c. cinclus* exhibits notably low genetic diversity, with low microsatellite polymorphism and very few single-nucleotide polymorphisms across sequenced nuclear introns compared to other passerines (Øigarden et al., 2010).

High-quality reference genomes for *C. cinclus* can support studies of aquatic adaptation, climate responses, subspecies differentiation, and conservation genomics. Although bird genomes are relatively small, microchromosomes, particularly repeat-rich dot-chromosomes, remain challenging to assemble completely (Formenti et al., 2026; Hron et al., 2026). A chromosome-scale reference is available from a British individual, likely representing *C. c. gularis* based on its sampling locality (bCinCin1.1; Sharp et al., 2024). Here, we present a chromosome-level assembly from a Norwegian individual sampled within the range of *C. c. cinclus*, generated using Oxford Nanopore long-read simplex sequencing and Hi-C scaffolding. Comparison with bCinCin1.1.pri and assemblies generated from our own PacBio HiFi data allows us to assess chromosome-scale synteny and how sequencing technology and sequence composition affect recovery of the smallest chromosomes.

The public availability of this genome resource will support future research on population genomics and genetics of the white-throated dipper. This genome assembly was generated as part of the Earth BioGenome Project Norway.

## Material and Methods

### Sample acquisition and DNA extraction

Blood sampled from the brachial vein of an adult female *Cinclus cinclus cinclus* was collected at Hærland, Indre Østfold, Viken (59.549, 11.411) on 3rd May 2022, under permission from the Norwegian Food Safety Authority (FOTS ID 29575) and the Norwegian Environment Agency (ringing licence 159). Accession number in the DNA bank of the Natural History Museum, University of Oslo: NHMO-BI-108605/2-B.

DNA isolation for Oxford Nanopore Technologies (ONT) and PacBio long read sequencing started from 25µl frozen blood which were split over two Circulomics Nanobind CBB BIG DNA kit reactions (disks), following the manufacturer’s recommendations (Circulomics Inc, now a PacBio company). Quality check of the amount, purity and integrity of the isolated DNA was performed using a combination of Qubit dsDNA BR quantification assay kit (Thermo Fisher), Nanodrop (Thermo Fisher), and Fragment Analyser (DNA HS 50kb large fragment kit, Agilent Tech.).

### Library preparation and sequencing for *de novo* assembly

Before PacBio HiFi library preparation, DNA was purified an additional time using AMPure PB beads (1:1 ratio). Purified HMW DNA was sheared into an average fragment size of 15-20 kbp large fragments using the Megaruptor3 (Diagenode). A HiFi library was prepared following the PacBio protocol for HiFi library preparation using the SMRTbell® Prep Kit 3.0. The final HiFi library was size-selected with a 10 kbp cut-off using a BluePippin (Sage Sciences) and sequencing was performed by the Norwegian Sequencing Centre on a PacBio Sequel IIe instrument (Pacific Biosciences Inc). The library was sequenced on two 8M SMRT cells using a Sequel II Binding kit 3.2 and Sequencing chemistry v2.0.

ONT library preparation involved fragmenting 7.5µg high molecular DNA in 200µl volume of low TE buffer with speedcode 30+31 on the Megaruptor instrument (Diagenode) to achieve an average fragment length of 30-35kb, of which 3µg was used as input for library preparation using the ONT Ligation Sequencing Kit V14 (SQK-LSK114) by following the protocol outlined in the human variation sequencing from 30kb extracted cell line samples (using SQK-LSK114 protocol). The final library was sequenced on a single R10.4.1 PromethION flow cell (FLO-PRO114M) using a PromethION 2 Integrated (P2i) device. Reads were basecalled using Dorado v1.1.1 with the super-accurate model dna_r10.4.1_e8.2_400bps_sup@v5.2.0.

A Hi-C library was prepared from 10-15µl frozen nucleated blood using the Arima High Coverage HiC kit (Arima Genomics Inc.) by following the manufacturer’s recommendations (document part number A160162v01). Final library quality was assayed as above in addition to qPCR using the Kapa Library quantification kit for Illumina (Roche Inc.). The library was sequenced with other libraries on the Illumina NovaSeq SP flowcell with 2*150 bp paired end mode at the Norwegian Sequencing Centre (https://www.sequencing.uio.no).

### Genome assembly and curation, annotation and evaluation

A full list of relevant software tools and versions is presented in Table 1. We assembled the species using a pre-release of the EBP-Nor genome assembly pipeline (https://github.com/ebp-nor/GenomeAssembly), with some additional analyses as needed. KMC (Kokot et al., 2017) was used separately to count k-mers of size 32 in the ONT and PacBio HiFi reads, excluding k-mers occurring more than 10,000 times in both datasets. GenomeScope (Ranallo-Benavidez et al., 2020) was run on the k-mer histogram output from KMC, and used to estimate genome size, heterozygosity and repetitiveness. Ploidy level was investigated using Smudgeplot. HiFiAdapterFilt (Sim et al., 2022) was applied to the HiFi reads to remove possible remnant PacBio adapter sequences. Long-read assemblies were generated using hifiasm (Cheng et al., 2021) with Hi-C integration. In total, six assemblies were generated, comprising two pseudo-haplotype assemblies for each of the ONT, HiFi, and HiFi+ONT assembly strategies. For brevity, the two pseudo-haplotypes within each assembly strategy are hereafter referred to as hap1 and hap2.

**Table 1.**
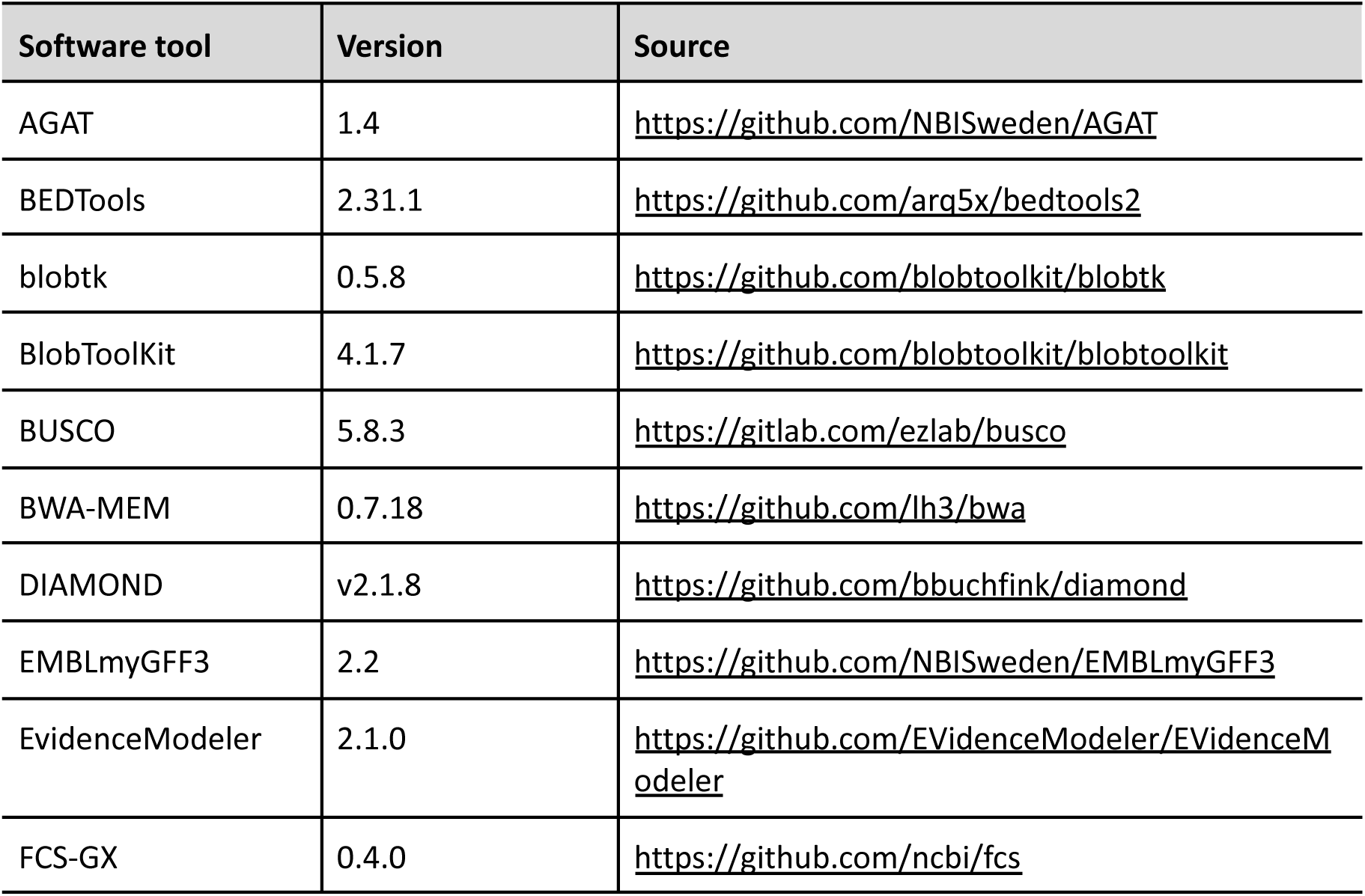
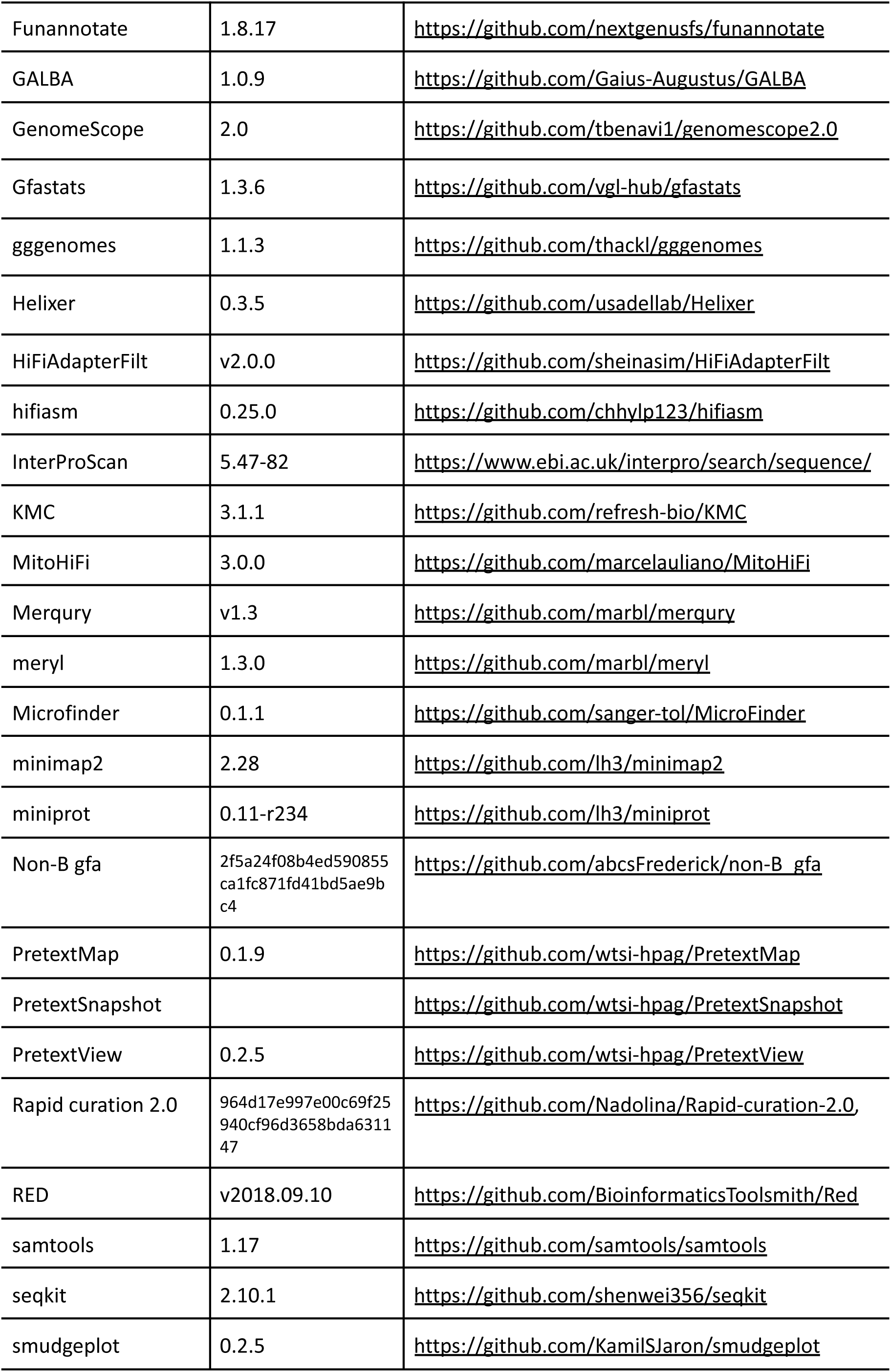
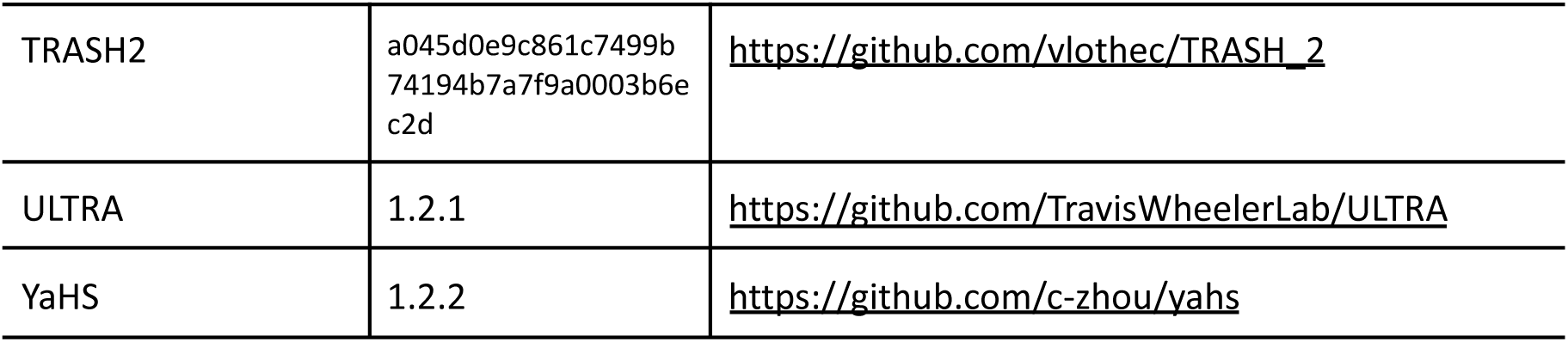
Software tools: versions and sources.

For each assembly strategy, meryl (Rhie et al., 2020) was used to identify k-mers unique to each pseudo-haplotype. These k-mers were used to generate two filtered Hi-C read sets: one excluding reads containing k-mers unique to hap1, and one excluding reads containing k-mers unique to hap2. The k-mer-filtered Hi-C reads were aligned to each assembly using BWA-MEM (H. Li, 2013) with the -5SPM options. Alignments were name-sorted with samtools (H. Li et al., 2009), after which samtools fixmate was used to remove unmapped reads and secondary alignments and to add mate scores. Duplicates were then removed using samtools markdup. The resulting BAM files were used for scaffolding with YaHS (Zhou et al., 2023) using default options. Assemblies were screened for contamination using FCS-GX (Astashyn et al., 2024), and contaminated sequences were removed. Merqury was used to assess assembly completeness and quality by comparing each assembly to the k-mer content of the Hi-C, HiFi and ONT reads. Assemblies were manually curated in PretextView using Rapid Curation 2.0. Hi-C contact maps were generated by mapping Hi-C reads to the assemblies with BWA-MEM (H. Li, 2013), using the same parameters as for scaffolding. Contact maps were produced with PretextMap and visualized with PretextSnapshot. Chromosomes, including sex chromosomes, were identified by inspecting the Hi-C contact maps. A second curation pass focused on microchromosomes was performed using scaffolds ≤20 Mb. MicroFinder (Mathers et al., 2026) was used to map conserved dot-chromosome genes to scaffolds, and candidate dot-chromosomes were identified by generating a gene-density BED track for visualization in PretextView. Curated scaffolds were reintegrated into the assembly, followed by a final curation pass in PretextView. Gfastats (Formenti et al., 2022) was used to calculate assembly statistics. These statistics were visualized using BlobToolKit and BlobTools2 (Laetsch & Blaxter, 2017), together with blobtk. BUSCO (Manni et al., 2021) was used to assess genome completeness against the expected gene content of the aves lineage. MitoHiFi (Uliano-Silva et al., 2023) was used to identify mitochondrial DNA in the final assembly and read datasets.

We annotated the genome assemblies using a pre-release version of the EBP-Nor genome annotation pipeline (https://github.com/ebp-nor/GenomeAnnotation). First, AGAT (https://zenodo.org/record/7255559) agat_sp_keep_longest_isoform.pl and agat_sp_extract_sequences.pl were used on the *Gallus gallus* (bGalGal1.mat.broiler.GRCg7b) genome assembly and annotation to generate one protein (the longest isoform) per gene. Miniprot (H. Li, 2023) was used to align the proteins to the curated assemblies. UniProtKB/Swiss-Prot (UniProt Consortium, 2023) release 2025_03 in addition to the Aves part of OrthoDB v12 (Tegenfeldt et al., 2025) were also aligned separately to the assemblies. RED (Girgis, 2015) was run via redmask (https://github.com/nextgenusfs/redmask) on the assemblies to mask repetitive areas. In addition to gene models from protein alignments, GALBA (Brůna et al., 2023; Buchfink et al., 2015; Hoff & Stanke, 2019; H. Li, 2023; Stanke et al., 2006) was run with the chicken proteins using the miniprot mode on the masked assemblies to generate *ab initio* predicted gene models. Helixer (Holst et al., 2025) was run using the vertebrate-specific model (vertebrate_v0.3_m_0080).

The funannotate-runEVM.py script from Funannotate was used to run EvidenceModeler (Haas et al., 2008) on the alignments of chicken proteins, UniProtKB/Swiss-Prot proteins, Aves proteins and the predicted genes from GALBA and Helixer. The resulting predicted proteins were compared to the protein repeats that Funannotate distributes using DIAMOND blastp and the predicted genes were filtered based on this comparison using AGAT. The filtered proteins were compared to the UniProtKB/Swiss-Prot release 2025_03 using DIAMOND (Buchfink et al., 2015) blastp to find gene names and InterProScan (Jones et al., 2014) was used to discover functional domains. AGATs agat_sp_manage_functional_annotation.pl was used to attach the gene names and functional annotations to the predicted genes. EMBLmyGFF3 (Norling et al., 2018) was used to combine the fasta files and GFF3 files into a EMBL format for submission to ENA.

All the evaluation tools have also been implemented in a pipeline, similar to assembly and annotation (https://github.com/ebp-nor/GenomeEvaluation). Merqury (Rhie et al., 2020) was used to assess the completeness and quality of the genome assemblies by comparing them to the k-mer content of the Hi-C, PacBio HiFi, and ONT reads. BUSCO (Manni et al., 2021) was used to assess the completeness of the genome assemblies by comparing against the expected gene content in the Aves lineage. Gfastats (Formenti et al., 2022) was used to output different assembly statistics of the assemblies.

BlobToolKit and BlobTools2 (Laetsch & Blaxter, 2017), in addition to blobtk were used to visualize assembly statistics. To generate the Hi-C contact map image, the Hi-C reads were mapped to the assemblies using BWA-MEM (H. Li, 2013) using the same approach as above. Finally, PretextMap was used to create a contact map which was visualized using PretextSnapshot.

To characterize the differences between hap1 and hap2, we ran a genome alignment using minimap2 (H. Li, 2018) on the homologous chromosomes. The resulting alignment was processed with the minimap2-included paftools.js producing a report listing the numbers of substitutions, insertions, and deletions between them.

Whole-genome synteny was assessed among the bCinCin1.1 primary assembly and the two bCinCin5.1 hap1 and hap2 assemblies. Pairwise alignments were generated with minimap2 (H. Li, 2018) using the asm10 preset, first between bCinCin1.1.pri and bCinCin5.1 hap1, and then between bCinCin5.1 hap1 and hap2. PAF alignments with a mapping quality of at least 60 and an aligned length of at least 100 kb were visualized as chained synteny plots using gggenomes in R, with bCinCin5.1 hap1 connecting the two other assemblies. A zoomed plot was generated for chromosomes at or below the length of bCinCin5.1 hap1 chromosome 26, with retained MicroFinder (Mathers et al., 2026) hits overlaid per sequence. Here, MicroFinder was used to summarize conserved dot-chromosome gene matches across the retained synteny sequences, rather than to define curation candidates.

### Comparison of HiFi reads and assembly support against the ONT assembly

To examine differences in sequence recovery between ONT and PacBio HiFi data, and whether these differences were associated with chromosome class and local sequence composition, we compared HiFi read coverage and assembly support against the pseudo-haplotype-resolved ONT assembly.

PacBio HiFi and ONT reads were mapped separately to both pseudo-haplotypes of the bCinCin5.1 ONT assembly with minimap2 (Li, 2018), using the ‘map-hif’ and ‘map-ont’ presets, respectively. Coverage was calculated from all mapped alignments, including secondary and supplementary alignments. HiFi-only zero-coverage regions were defined as continuous intervals ≥1 kb without HiFi read coverage that did not overlap an ONT zero-coverage interval. Autosomes were assigned to chromosome classes using haplotype-specific chromosome length and MicroFinder hit counts. Chromosomes ≥20 Mb were classified as macrochromosomes. Among chromosomes <20 Mb, chromosomes with ≥40 retained MicroFinder hits were classified as dot-chromosomes, reflecting the clear discontinuity in hit counts, and the remaining chromosomes were classified as (non-dot) microchromosomes.

Non-B-DNA-forming motifs were predicted on the combined ONT assembly using GFA from the Non-B DB tool suite with default motif-calling parameters (Cer et al., 2013). The retained motif classes were A-phased repeats (A), inverted repeats (IR), Z-DNA motifs (Z), direct repeats (DR), G-quadruplex motifs (G4), mirror repeats (MR), and short tandem repeats (STR). GFA predictions were subsequently filtered to restrict direct-, inverted-, and mirror-repeat spacers to ≤10 bp, inverted-repeat arms to 6–30 bp, and mirror repeats to those consisting entirely of purines or entirely of pyrimidines; these filtering criteria were informed by Smeds et al. (2025). Default GFA predictions were retained for G4 and Z-DNA motifs. Tandem repeats (in general) and repeat arrays were identified with ULTRA (Olson & Wheeler, 2024) and TRASH2 (Wlodzimierz et al., 2023), respectively, using default parameters. Both annotations were retained to capture repeat organization at different hierarchical scales: TRASH2 array calls represent approximate envelopes of broad satellite-repeat regions, whereas ULTRA identifies individual tandem-repeat intervals with periods up to 100 bp. GC content was calculated with bedtools nuc (Quinlan & Hall, 2010). Gene positions were obtained from the gene annotations described above, and terminal telomeric regions were identified from ULTRA annotations.

The combined pseudo-haplotype-resolved HiFi assembly was aligned to the combined ONT assembly with minimap2 using the asm10 preset (Li, 2018), and the two ONT pseudo-haplotypes were aligned reciprocally using the same settings. The HiFi assembly was aligned to the ONT assembly, and CIGAR strings were used to identify ONT regions supported by HiFi alignments. Alignments ≥50 kb with MAPQ ≥30 were considered high confidence. Candidate transition points were boundaries of ≥20-kb gaps in merged HiFi-alignment outer spans. Candidate boundaries additionally required a high-confidence HiFi alignment in the adjacent supported block, limited residual alignment within the HiFi-poor interval, complete 20-kb flanks without assembly Ns, and ≥95% high-confidence reciprocal-ONT support across both flanks.

At the chromosome level, HiFi-depleted coverage was calculated as the proportion of each haplotype-specific autosome contained within HiFi-only depleted regions, and non-B DNA coverage as the proportion covered by merged non-B DNA intervals. Values from the two ONT pseudo-haplotypes were averaged for each chromosome. For sequence-composition analyses, assembly Ns and ONT zero-coverage regions were excluded. HiFi-depleted sequence comprised the retained HiFi-only depleted regions, whereas HiFi-covered sequence comprised bases with HiFi coverage greater than zero; other HiFi zero-coverage intervals were excluded from both groups. Sequence was pooled across both pseudo-haplotypes within macro-, micro- and dot-chromosome classes, and enrichment was calculated as the fraction of HiFi-depleted sequence occupied by each feature divided by the corresponding fraction in HiFi-covered sequence. GC enrichment was calculated from GC fractions, with GC expressed relative to ACGT bases. To quantify sequence supported by the reciprocal ONT haplotype but lacking HiFi-assembly support, HiFi-supported regions were subtracted from reciprocal-ONT-supported regions. The remaining sequence was expressed as a percentage of chromosome length and averaged between pseudo-haplotypes for each chromosome.

Retained transition boundaries were oriented with the HiFi-supported sequence to the left and the HiFi-poor sequence to the right. A 40-kb window centred on each transition was divided into eight non-overlapping 5-kb bins. Mean ONT and HiFi read coverage, GC content, and the fraction of each bin covered by each non-B DNA class, ULTRA tandem repeats and TRASH2 repeat arrays were calculated. When both boundaries of the same HiFi-poor interval were retained and separated by less than 40 kb, their corresponding oriented bin values were averaged to avoid counting overlapping windows as independent profiles.

Within each chromosome class, mean values and t-based 95% confidence intervals were calculated across profiles for each 5-kb bin. Relative change across the transition was calculated separately for each profile by comparing the mean of the two bins immediately before and after the transition as 100 × (post − pre) / pre; these per-profile changes were then averaged within chromosome class.

Chromosome-scale visualizations were generated using the coverage, sequence-annotation, assembly-support, HiFi-only zero-region and transition-point tracks described above. H1 chromosome 37 was used as the representative chromosome-scale example in Figure 4A. Equivalent plots for all autosomes were compiled as Supplementary Figure 4, with H1 and H2 displayed side by side in their respective coordinate systems. The W and Z chromosomes were displayed together at the end of Supplementary Figure 4.

Methods and Results sections are based on a standardized template used across species published by the EBP-Nor project.

## Results

### *De novo* genome assembly and annotation

The genome of the female *Cinclus cinclus cinclus* (Figure 1) had an estimated genome size of 1.14 Gb, with 0.284% heterozygosity and a bimodal distribution based on the k-mer spectrum (Figure 1C). A total of 38-fold coverage in Oxford Nanopore R10 reads and 101-fold coverage in Arima Hi-C reads resulted in two pseudo-haplotype-separated assemblies. The final assemblies have total lengths of 1186 Mb and 1115 Mb (Table 2 and Figure 2), respectively. Hap1 and hap2 have scaffold N50 size of 74.2 Mb and 64.6 Mb, respectively, and contig N50 of 35.8 Mb and 32.0 Mb, respectively (Table 2, Figure 2). GC-coverage profiles for both assemblies are shown in Supplementary Figure 2. 40 autosomes were identified in both pseudo-haplotypes, with sex chromosomes W and Z added to hap1. Chromosomes were named primarily according to the bCinCin1.1.pri (GCF_963662255.1) assembly, derived from a British individual putatively assigned to the subspecies *Cinclus cinclus gularis*, and secondarily by length in hap1.

**Figure 1.**
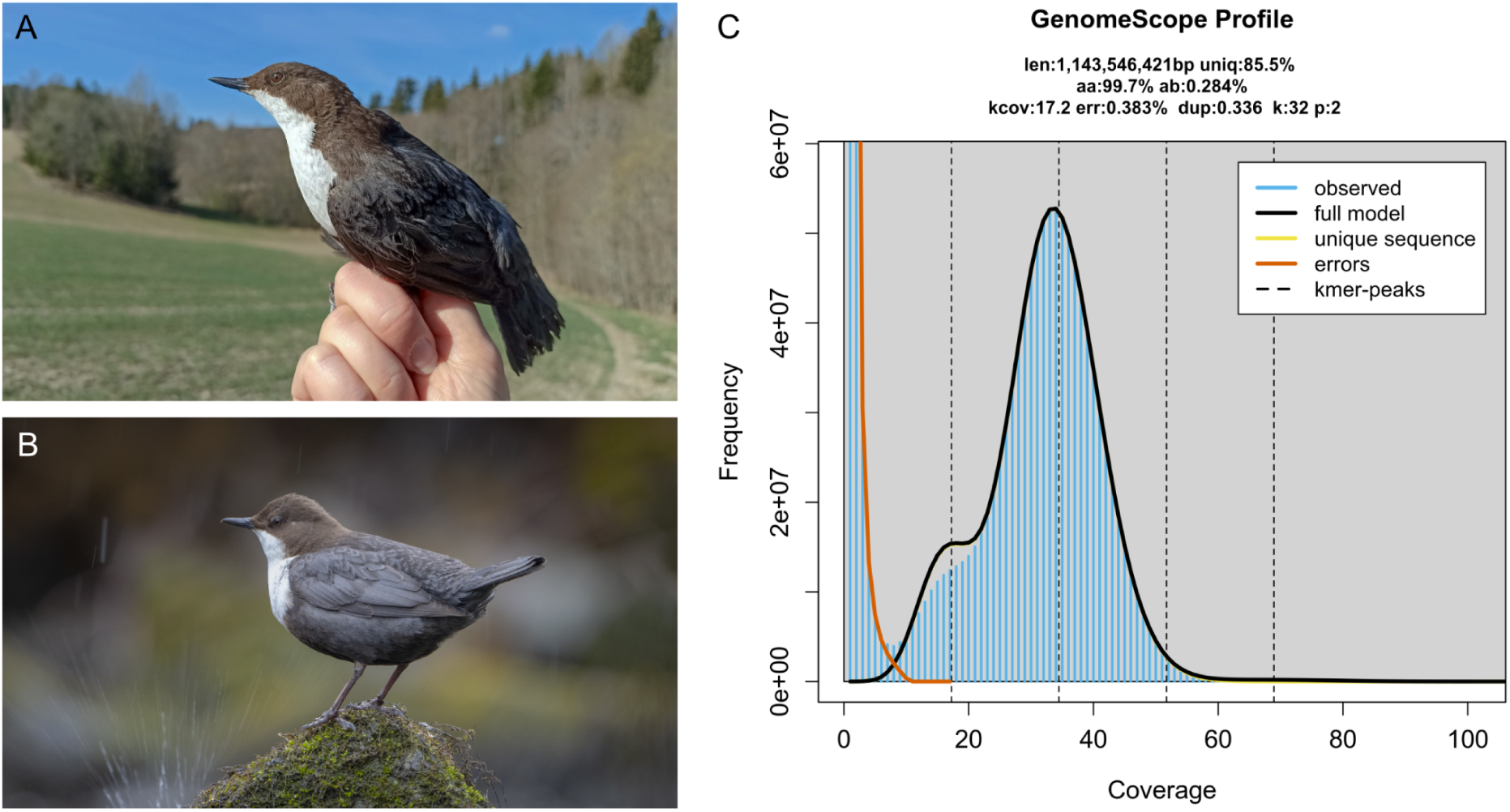
Sequenced specimen and genome profile. **A)** The female bird used for sequencing; photo by Laima Bagdonaitė, reproduced with permission. **B)** *C. c. cinclus* in its natural habitat; photo by Bjørn Aksel Bjerke, reproduced with permission. **C)** GenomeScope profile of the ONT reads from the sequenced individual. This analysis estimates a 1144 Mb genome, with 0.284% heterozygosity. The left-hand shoulder peak of k-mers corresponds to k-mers from heterozygous regions of the genome, while the right-hand peak is from homozygous regions.

**Figure 2:**
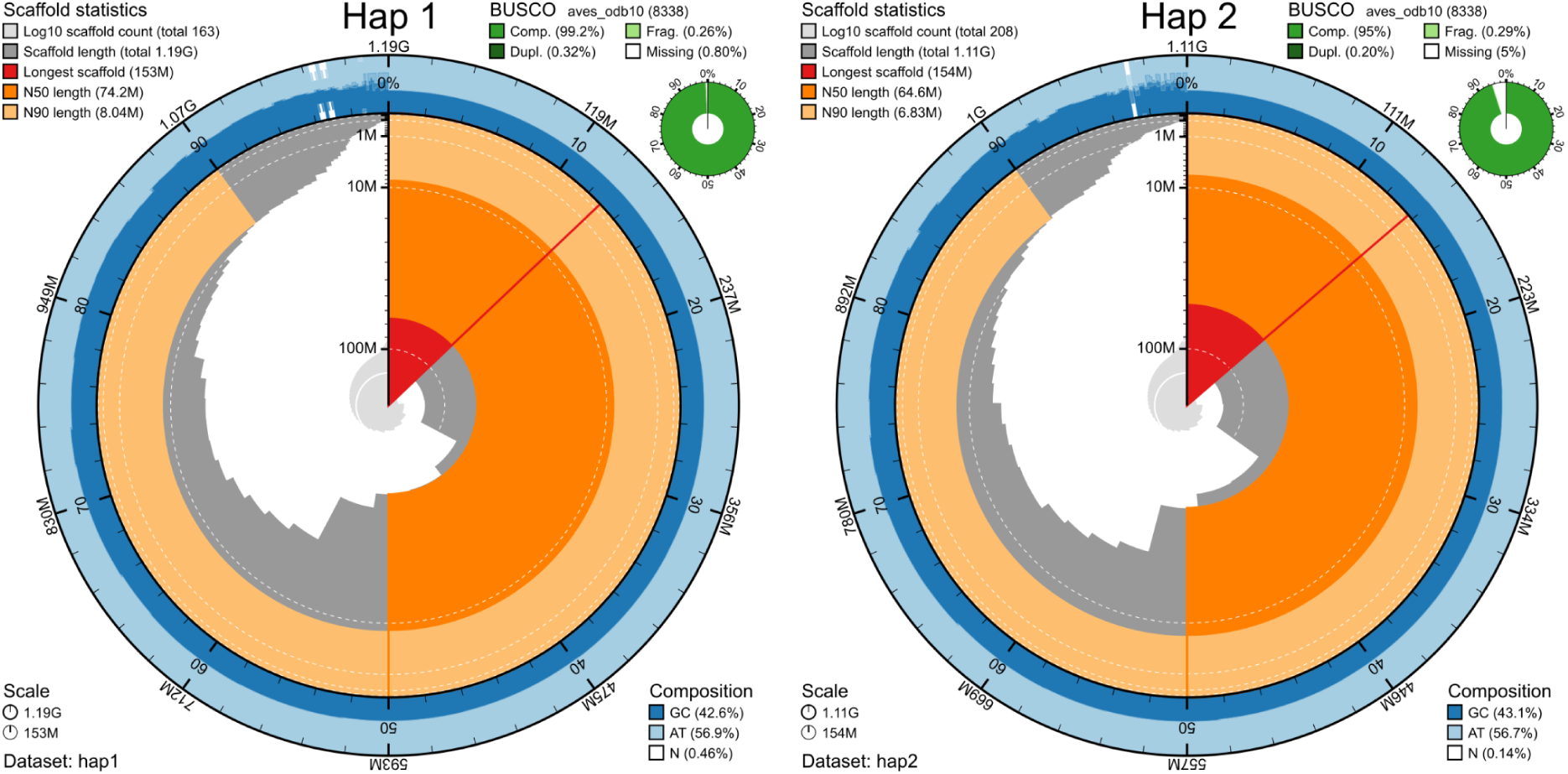
Metrics of the genome assemblies of *C. c. cinclus* hap1 and hap2. The BlobToolKit Snailplots show N50 metrics and BUSCO gene completeness. The two outermost bands of the circle signify GC versus AT composition at 0.1% intervals, with mean, maximum, and minimum. The third outermost shows the N90 scaffold length, while the fourth is N50 scaffold length. The line from middle to second outermost band shows the size of the largest scaffold. All the scaffolds are arranged in a clockwise manner from largest to smallest, and shown in darker gray with white lines at different orders of magnitude, shown as a scale on the radius. The light gray shows the cumulative scaffold count. The scale inset in the lower left corner shows the total amount of sequence in the whole circle, and the fraction of the circle encompassed in the largest scaffold.

**Table 2:** Genome data for *C. c. cinclus*.

| Project accession data |  |  |
| --- | --- | --- |
| Species | *Cinclus cinclus cinclus* |  |
| Specimen | bCinCin5 |  |
| NCBI Taxonomy ID | 127875 |  |
| BioProject | PRJEB114807 |  |
| BioSample ID | SAMEA122794430 |  |
| Isolate information | Blood sampling from brachial vein |  |
| Raw data accessions |  |  |
| PacBio HiFi reads | ERR17389582 and ERR17389584 | 2 PACBIO_SMRT (Sequel II) runs: 4.9 M reads, 77.66 Gb |
| Hi-C Illumina reads | ERR17389581 and ERR17389583 | 2 ILLUMINA (Illumina NovaSeq SP) run: 397 M pairs of reads, 119.87 Gb |
| ONT reads | ERR17389699 | 1 ONT (PromethION P2i) runs: 3.2 M reads, 44.67 Gb |
| Genome assembly metrics |  |  |
| ONT read coverage | 38 |  |
| Hi-C read coverage | 101 |  |
| Assembly accession | GCA_985200095 | GCA_985200165 |
| Assembly identifier | bCinCin5.1.hap1 | bCinCin5.1.hap2 |
| Span (Mb) | 1,186 | 1,115 |
| Number of contigs | 215 | 256 |
| Contig N50 length (Mb) | 35.8 | 32.0 |
| Longest contig (Mb) | 110.7 | 152.3 |
| Number of gaps | 52 | 48 |
| Number of scaffolds | 163 | 208 |
| Scaffold N50 length (Mb) | 74.2 | 64.6 |
| Longest scaffold (Mb) | 153.1 | 153.6 |
| GC content (%) | 42.84 | 43.18 |
| Percentage of assembly mapped to chromosomes | 96.7 | 94.4 |
| Sex chromosomes | W, Z |  |
| Organelles | MT |  |
| Consensus quality (QV) compared to |  |  |
| Hi-C | 41.21 | 41.77 |
| Hi-C both | 41.47 |  |
| HiFi | 35.39 | 35.25 |
| HiFi both | 35.32 |  |
| ONT | 64.36 | 63.94 |
| ONT both | 64.15 |  |
| K-mer completeness (%) compared to |  |  |
| Hi-C | 96.8 | 96.4 |
| Hi-C both | 97.8 |  |
| HiFi | 97.7 | 90.2 |
| HiFi both | 99.8 |  |
| ONT | 97.0 | 89.7 |
| ONT both | 99.4 |  |
| **Genome annotation metrics** |  |  |
| Number of protein-coding genes | 19,003 | 17,746 |
| Number of protein-coding genes with functional domain* | 18,685 | 17,410 |
| Number of protein-coding genes with gene names | 16,149 | 14,985 |
| BUSCO** | C:99.2%[S:98.9%,D:0.3%],F:0.3%,M:0.5%,n:8338 | C:95.0%[S:94.8%,D:0.2%],F:0.3%,M:4.7%,n:8338 |

\*Number of genes annotated with a functional domain as found by InterProScan. \*\* BUSCO scores based on the aves\_odb10 BUSCO set using v5.8.3. C = complete [S = single copy, D = duplicated], F = fragmented, M = missing, n = number of orthologues in comparison.

When compared with a k-mer database derived from the Hi-C reads, hap1 and hap2 had k-mer completeness values of 96.8% and 96.4%, respectively, increasing to 97.8% when the two pseudo-haplotypes were considered together. The corresponding assembly consensus quality values (QV) were 41.21 for hap1 and 41.77 for hap2, where a QV of 40 corresponds to approximately one error per 10,000 bp, or 99.99% accuracy (Table 2). K-mer completeness based on the ONT reads was 97.0% for hap1 and 89.7% for hap2, increasing to 99.4% for the combined assembly (Table 2; Supplementary Figure 3). When comparing the two pseudo-haplotypes using minimap2, there are 1,143,264 SNP differences (0.11% of the aligned sequence), 144,034 deletions in hap2 compared to hap1 ranging from 1 bp to more than 1,000 bp and 143,280 insertions from 1 bp to more than 1,000 bp in size (Supplementary Table 1). A total of 19,003 and 17,746 protein-coding genes were annotated in hap1 and hap2, respectively (Table 2).

Hi-C contact maps showed clear separation of chromosomes into homologous sets in both *C. c. cinclus* hap1 and hap2 (Supplementary Figure 1). This separation was clearest for macrochromosomes, whereas microchromosomes showed elevated background contact signals and less sharply resolved boundaries, consistent with previous observations in birds and reptiles (Perry et al., 2021).

Chromosome-scale synteny was largely conserved between the bCinCin1.1.pri assembly and both bCinCin5.1 hap1 and hap2, with mostly one-to-one correspondence across macrochromosomes and larger microchromosomes (Figure 3). Chromosome names and orientations in *C. c. cinclus* were assigned to match the published bCinCin1.1.pri assembly where possible, and otherwise by chromosome size. This mainly affected the smallest microchromosomes, where syntenic support was weaker and chromosome size order was less consistent. In particular, regions homologous to bCinCin1.1.pri chromosome 36 were distributed across bCinCin5.1 chromosomes 36, 39, and 40, indicating that these chromosome labels are not directly comparable between assemblies.

**Figure 3.**
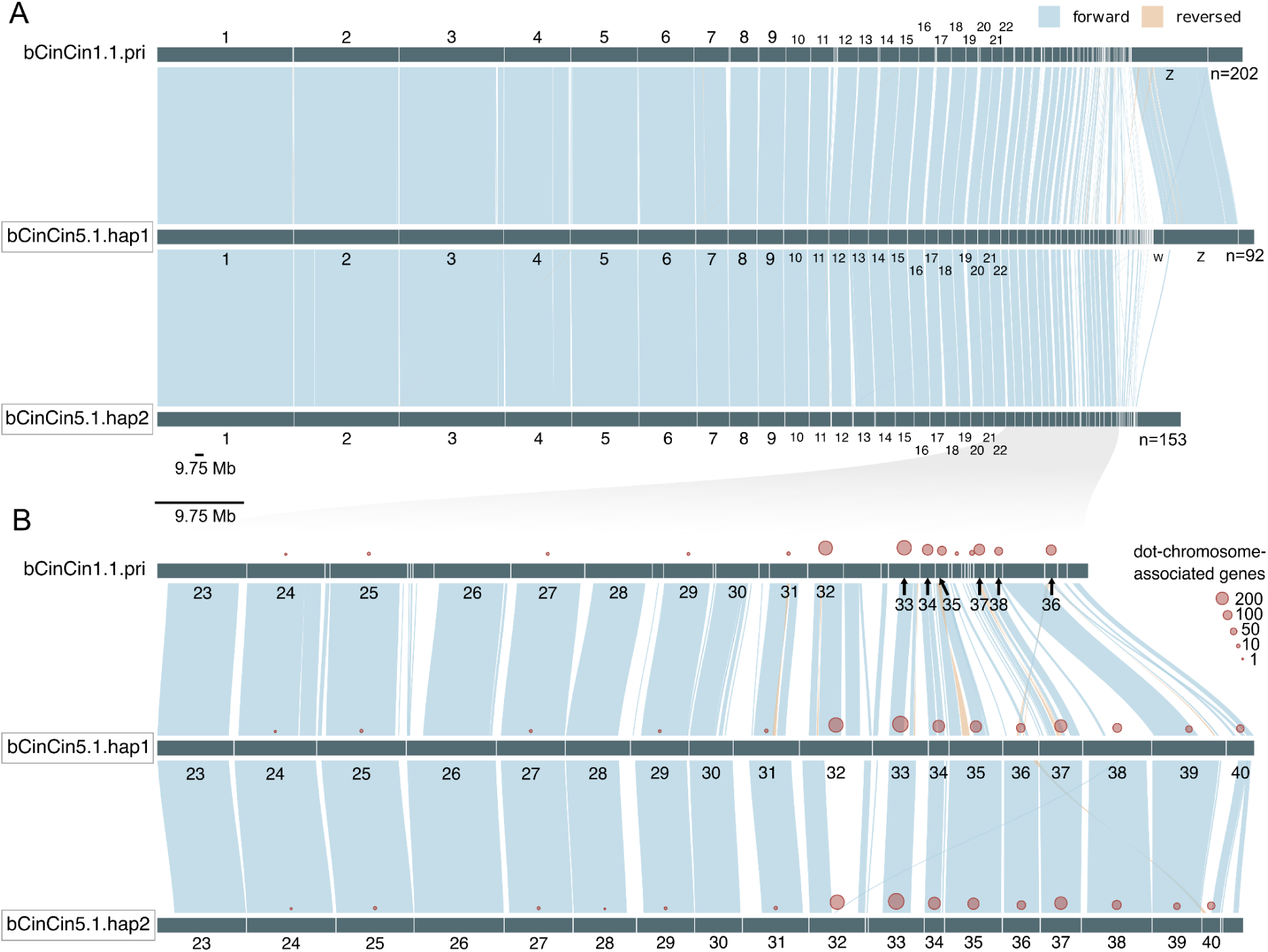
Chromosome-scale synteny between the British *C. c. gularis* assembly (bCinCin1.1.pri, top) and the two Norwegian *C. c. cinclus* pseudo-haplotypes (bCinCin5.1.hap1 and bCinCin5.1.hap2, bottom). Chromosomes and chromosome-associated scaffolds were ordered and oriented relative to bCinCin5.1.hap1. When a source chromosome matched multiple sequences, these were arranged by inferred position to minimize crossing links. Blue and orange links indicate forward and reverse alignments, respectively, and arrows (in panel B) mark sequences reversed from their original orientation. Only alignments ≥100 kb with mapping quality 60 are shown. Only scaffolds marked as chromosomes are named. **A)** Full assembly view, including sequences without retained alignments; unconnected non-chromosomal sequences are aggregated. **B)** Enlarged view of source sequences ≤9.75 Mb and their aligned counterparts. Shading indicates the region enlarged in panel B. Red circles show the number of hits to conserved dot-chromosome-associated genes, with circle area proportional to hit count.

We compared three assembly strategies generated here: ONT-derived, HiFi-derived, and HiFi+ONT—with the pre-existing HiFi-based bCinCin1.1.pri hifiasm primary assembly. All were scaffolded with Hi-C. Scaffold N50 values were similar across assemblies (61–74 Mb), but contig-level continuity and completeness differed substantially (Supplementary Table 2). The ONT-derived assemblies were the least fragmented, spanning 1,186 and 1,115 Mb with contig N50 values of 35.8 and 32.0 Mb. HiFi+ONT assemblies spanned 997 and 1,044 Mb with contig N50 values of 8.9 and 9.0 Mb, while HiFi-derived assemblies spanned 1,132 and 903 Mb with contig N50 values of 6.1 and 7.5 Mb. In all three assembly types generated here, Z and W were assigned to hap1 and were absent from hap2. The HiFi-based bCinCin1.1.pri primary assembly had a similar span to ONT hap1 but a substantially lower contig N50 of 3.6 Mb.

ONT hap1 also had the highest BUSCO completeness of any individual assembly at 99.2%, closely followed by bCinCin1.1 primary at 99.1%, while ONT hap2 was 95.0% complete. Since the hifiasm non-haplotype resolved method assemblies combined long phased blocks without preserving a single biological haplotype throughout, the union of BUSCO calls across ONT hap1 and hap2 was used to approximate bCinCin1.1 primary gene-space completeness. This union recovered all complete BUSCO orthologs found in bCinCin1.1 primary plus eight additional complete orthologs, yielding 99.23% completeness compared with 99.14% for bCinCin1.1 primary.

Chromosome-scale comparisons of the ONT and HiFi datasets showed variation in read coverage and assembly support across the *C. c. cinclus* genome. Hap1 chromosome 37 illustrates this pattern, containing multiple HiFi-only zero runs and gaps in HiFi-assembly support, whereas ONT read coverage and assembly support from the alternate ONT pseudo-haplotype (hap2) extend across nearly the entire chromosome (Figure 4A). Similar chromosome-scale variation was observed across both pseudo-haplotypes for the remaining chromosomes (Supplementary Figure 4). When values were averaged between pseudo-haplotypes for each chromosome, the median proportion of HiFi-depleted sequence was 0.43% among macrochromosomes (range 0.09–1.46%; n = 15), 1.30% among microchromosomes (0.31–13.65%; n = 16), and 9.33% among dot-chromosomes (2.05–21.77%; n = 9). Predicted non-B-DNA coverage had a median of 6.13% among macrochromosomes (range 5.58–7.85%), 9.64% among microchromosomes (6.60–19.06%), and 18.42% among dot-chromosomes (11.76–39.76%) (Figure 4B).

**Figure 4.**
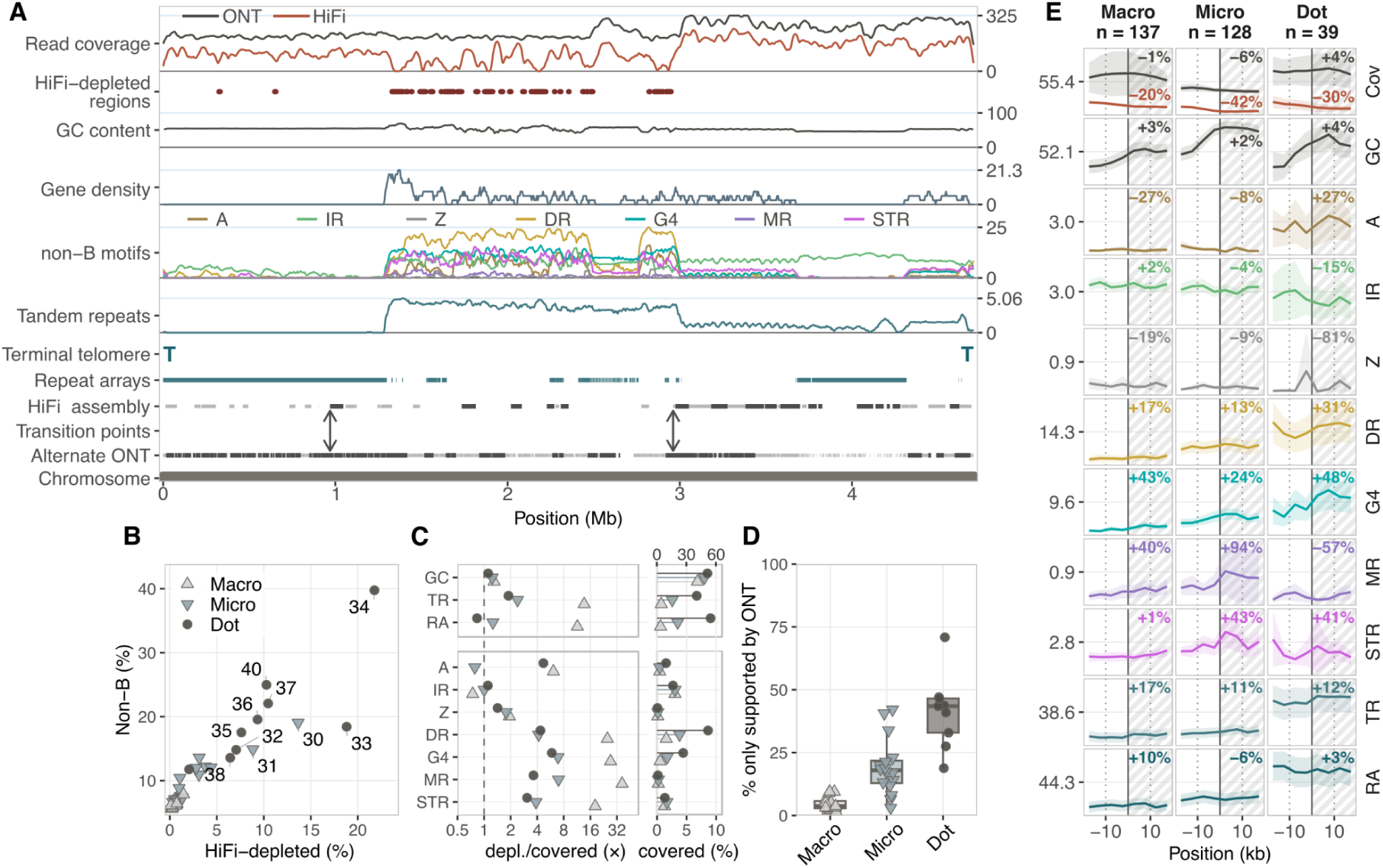
Chromosome-scale and local patterns of HiFi read and assembly support in bCinCin5.1. **A) Chromosome-scale context for H1 chromosome 37.** Tracks show smoothed ONT and HiFi read coverage, HiFi-depleted regions, GC content, gene-start density, predicted non-B-DNA motif density, ULTRA tandem-repeat density, terminal telomeres, TRASH2 repeat arrays, assembly support, retained transition points, and chromosome span. HiFi-depleted regions are continuous intervals ≥1 kb with zero HiFi coverage that do not overlap ONT zero coverage; these and other interval tracks retain their exact coordinates. The “HiFi assembly”-support track pools alignments from both HiFi pseudo-haplotypes, whereas “Alternate ONT” shows the hap2 assembly aligned to hap1. Light grey denotes support from all alignments, and dark grey high-confidence support ≥50 kb with MAPQ ≥30. Double-headed arrows mark retained transition points. Coverage and count tracks were log1p transformed and smoothed for display, with gene starts expressed per 100 kb. Right-hand values show the joint 99th-percentile coverage limit, 100% for GC, and observed maxima for the remaining quantitative tracks. **B) Chromosome-level HiFi depletion and non-B-DNA territory.** Each point represents one named autosome, with hap1 and hap2 percentages averaged by chromosome name. The proportion of chromosome length in HiFi-depleted regions is plotted against the proportion covered by the union of predicted non-B-DNA motifs. Symbols distinguish macrochromosomes, microchromosomes, and dot-chromosomes; chromosomes with >5% HiFi-depleted sequence are labelled. **C) Sequence composition of HiFi-depleted regions.** Hap1 and hap2 sequence was pooled within chromosome class, and points show fold enrichment of each feature in HiFi-depleted relative to HiFi-covered territory. Eligible territory excludes assembly Ns and ONT zero-coverage intervals; HiFi-covered territory comprises the remaining sequence with nonzero HiFi coverage. Adjacent bars show the absolute percentage occupied by each feature in HiFi-covered territory, with endpoint symbols identifying chromosome class as in panel B. GC, TR, and RA use the 0–60% upper axis, whereas the remaining features use the 0–10% lower axis. GC is calculated relative to ACGT bases; other values are the fraction of territory overlapped by each feature. Fold enrichments are plotted on a log2 scale and labelled in fold-change units; the dashed line at 1 indicates equal values in HiFi-depleted and HiFi-covered territory. Feature classes can overlap and are therefore not additive. **D) Alternate-ONT-supported sequence lacking HiFi assembly support.** Support from both HiFi pseudo-haplotypes was pooled and subtracted from support provided by the opposing ONT hap. The remaining sequence, supported by the alternate ONT hap but by neither HiFi hap, was expressed as a percentage of the full ONT target chromosome. Each point is the hap1/hap2 mean for one named chromosome; boxplots summarize chromosome-level values within chromosome classes. **E) Profiles across ONT-conditioned HiFi assembly transitions.** Retained boundaries were oriented with HiFi-supported sequence to the left and HiFi-poor sequence to the right. Lines show chromosome-class means in non-overlapping 5-kb bins across a 40-kb window, with t-based 95% confidence intervals among profile units. The solid line marks the transition at 0 kb, dotted lines mark −10 and +10 kb, and hatching denotes the HiFi-poor side. Percentage labels compare the mean of the two bins immediately before (−10 to 0 kb) and after (0 to +10 kb) the transition. Cov denotes ONT and HiFi read coverage; remaining rows show GC content or the fraction of each bin covered by the indicated feature. **Abbreviations** A: A-phased repeat, IR: inverted repeat, Z: Z-DNA motif, DR: direct repeat; G4: G-quadruplex motif, MR: mirror repeat, STR: short tandem repeat, TR: ULTRA tandem repeat, RA: TRASH2 repeat array.

Compared with HiFi-covered sequence, HiFi-depleted sequence differed in several annotated sequence features (Figure 4C). HiFi-depleted regions were enriched for several repeat- and non-B-DNA-associated features, although the magnitude and, for some features, the direction of these differences varied among chromosome classes. Dot-chromosomes had higher background G4 and DR coverage than microchromosomes (4.42% versus 1.72% and 8.62% versus 3.81%, respectively), but both features remained enriched in HiFi-depleted sequence by 5.9-fold and 4.3-fold. Macrochromosomes showed the strongest relative enrichments for G4 and DR (27.0-fold and 24.9-fold), largely because their background coverage was particularly low (0.52% and 1.56%).

The chromosome-class differences were also apparent at the assembly level. Sequence that aligned between the two ONT pseudo-haplotypes but lacked corresponding HiFi assembly support accounted for a median of 3.73% of macrochromosomes (IQR 2.87–5.78%), compared with 17.95% of microchromosomes (12.86–21.80%) and 43.47% of dot-chromosomes (32.92–46.53%) (Figure 4D). Thus, the proportion of ONT-supported sequence without corresponding HiFi assembly support was highest on the dot-chromosomes.

At transitions from HiFi-supported to HiFi-poor sequence, 304 filtered transition profiles were retained: 137 from macrochromosomes, 128 from microchromosomes, and 39 from dot-chromosomes (Figure 4E). HiFi coverage decreased across these transitions by 20%, 42%, and 30%, respectively, whereas ONT coverage changed little (−1%, −6%, and +4%) and GC content increased only modestly (+3%, +2%, and +4%). G-quadruplex coverage increased in all three chromosome classes (+43%, +24%, and +48%), as did direct repeats (+17%, +13%, and +31%). GFA-predicted STRs changed little on macrochromosomes (+1%) but increased on microchromosomes (+43%) and dot-chromosomes (+41%).

Repeat abundance around these transitions differed more strongly among chromosome classes than across the transition itself. Across the two 10-kb regions compared around each boundary, tandem repeats occupied 16–19% of macrochromosomal, 20–23% of microchromosomal, and 47–52% of dot-chromosome sequence, while repeat arrays occupied approximately 20–22%, 27–28%, and 55–56%, respectively. The high repeat abundance around dot-chromosome transitions therefore primarily reflected their broader sequence background rather than a sharp transition-associated increase.

The transition profiles represent a filtered subset of well-supported boundaries rather than an unbiased sample of all candidate transitions. Candidate retention was lower for dot-chromosomes (16%) than for macrochromosomes (54%) or microchromosomes (43%) because of the reciprocal ONT-support and assembly-gap filters. Retained macrochromosomal transitions were also concentrated near chromosome ends.

## Discussion

Here, we present pseudo-haplotype-resolved ONT-derived assemblies of *C. c. cinclus*, comprising 2 × 40 autosomes, the W and Z sex chromosomes, and the mitochondrial genome. We initially generated HiFi+ONT and HiFi-derived assemblies, but these were substantially more fragmented and showed greater asymmetry between pseudo-haplotypes. We therefore used the ONT-only mode of hifiasm, which constructs assembly graphs directly from corrected R10.4.1 simplex reads (Cheng et al., 2026), to generate a substantially more contiguous pair of pseudo-haplotype-resolved assemblies. Compared with the HiFi-based British *C. c. gularis* reference assembly bCinCin1.1.pri (Sharp et al., 2024), our hap1 assembly includes the W chromosome, has a greater total assembled length, and extends several dot-chromosomes. In addition, the sequence represented as a single dot-chromosome (chr 36) in bCinCin1.1.pri is resolved into three distinct dot-chromosomes (chr 36, 39, and 40), supported by Hi-C contact patterns. The resulting complement of 40 autosomes, together with the Z and W sex chromosomes, implies a diploid chromosome number of 2n = 82, higher than the 2n = 78 reported for the closely related *C. pallasii* (Q.-W. Li et al., 1994). However, cytological estimates of avian chromosome number can be uncertain because the smallest microchromosomes are particularly prone to loss, overlap, or misidentification in metaphase preparations (de Bello Cioffi et al., 2026). The discrepancy may therefore reflect incomplete resolution of the smallest chromosomes in previous karyotypic or genomic resources.

Although scaffold N50 values were similar across assembly strategies, the ONT-derived assemblies were substantially less fragmented at the contig level than either the HiFi+ONT or HiFi-derived assemblies. In line with Cheng et al. (2026), adding ONT reads to a HiFi-derived assembly graph improved some metrics but did not match the contiguity or sequence recovery achieved by assembling directly from ONT reads. The bCinCin1.1 primary assembly and ONT hap1 had nearly identical BUSCO completeness (99.1% and 99.2%, respectively), despite substantial differences in contiguity and recovered sequence: bCinCin1.1.pri had an approximately one tenth the contig N50 (3.6 versus 35.8 Mb) and a ∼15.6 Mb shorter total scaffold span. Although direct comparison should be treated cautiously because the assemblies represent different subspecies, this disparity illustrates the limitations of conventional whole-genome metrics for evaluating avian assemblies. BUSCO markers are sparse on avian dot-chromosomes, making BUSCO completeness relatively insensitive to substantial sequence loss from these chromosomes (Mathers et al., 2026); likewise, scaffold N50 and total assembly span are dominated by macrochromosomes and can mask incomplete recovery of micro- and dot-chromosomes. Chromosome-aware comparisons, including the use of conserved dot-chromosome markers, are therefore particularly important for assessing avian genome completeness. This distinction is consequential for both gene annotation and evolutionary inference, because assembly-derived absence can otherwise be misinterpreted as genuine evolutionary loss (Hron et al., 2026).

HiFi-depleted regions were enriched for repeat- and predicted non-B-DNA-associated features, particularly G-quadruplexes and direct repeats, whereas ONT coverage remained comparatively stable. HiFi depletion increased from macro- to micro- to dot-chromosomes, where repeat and non-B-DNA content was also highest overall. This pattern is consistent with zebra finch analyses showing a stronger association between non-B-DNA content and HiFi than ONT coverage (Smeds et al., 2025), and with experimental evidence that such motifs, particularly G-quadruplexes, can impede SMRT polymerases and reduce sequencing depth and accuracy (Guiblet et al., 2018). ONT is also affected by non-B-DNA structures, which can alter nanopore translocation dynamics (Hosseini et al., 2023), and the zebra finch T2T study of Formenti et al. (2026) reported reduced yield and pore blockage. Nevertheless, both the zebra finch and white-throated dipper data indicate that these sequence features disproportionately impair HiFi sequence recovery, particularly on micro- and dot-chromosomes.

The zebra finch T2T assembly required multiple sequencing platforms, assemblers, and extensive manual graph curation, and no single automated assembly recovered all chromosomes completely (Formenti et al., 2026). By comparison, standard R10.4.1 simplex ONT reads assembled with hifiasm (ONT) and Hi-C recovered avian dot-chromosomes substantially better than the HiFi-based strategies tested here, despite similar conventional genome-wide metrics. A hybrid mode of hifiasm under development may further combine the complementary strengths of ONT and HiFi sequencing when both data types are available (https://github.com/chhylp123/hifiasm/commit/06cf12a; hybrid_v1).

Because *C. cinclus* is the most widely distributed dipper and comprises multiple subspecies (Tyler & Ormerod, 1994), this ONT-based reference assembly provides a resource for comparative and population genomics across its range. Together with other white-throated dipper genomes, including the British assembly (Sharp et al., 2024), it can support analyses of genomic differentiation and subspecies boundaries while providing more complete representation of repeat-rich dot-chromosomes. The strong chromosome-scale collinearity between the British and Norwegian assemblies suggests limited large-scale structural divergence, although broader sampling will be required to assess population- and subspecies-level variation. Ultimately, a graph-based pangenome spanning multiple populations and subspecies would capture this variation more completely.

## Acknowledgements

This project received data management and infrastructure support from ELIXIR Norway, supported by the Research Council of Norway’s grant 350529, the University of Bergen, the University of Oslo, the Arctic University of Norway in Tromsø, the Norwegian University of Science and Technology and the Norwegian University of Life Sciences: NMBU. The authors acknowledge support from the National Infrastructure for High Performance Computing and resources provided by Sigma2 as well as Data Storage in Norway (project NN8013K) for computational work. The Norwegian Sequencing Centre generated the sequencing data used in this project (http://sequencing.uio.no). The authors thank Laima Bagdonaitė and Jan T. Lifjeld for field sampling of the specimen used in this study.

## Data Availability

Data generated for this study are available under ENA BioProject PRJEB114807. Raw PacBio sequencing data for the northern white-throated dipper (ENA BioSample: SAMEA122794430) are deposited in ENA under ERR17389582 and ERR17389584, while Illumina Hi-C sequencing data is deposited in ENA under ERR17389581 and ERR17389583 (collected in PRJEB114662). Base-called ONT reads are found in ERR17389699. Pseudo-haplotype one can be found in ENA at PRJEB114800, while pseudo-haplotype two is PRJEB114801.

Raw ONT data is deposited at NIRD Archive: https://doi.org/10.11582/2026.kl960bb6.

The genome assemblies and gene annotations are available at Zenodo: https://doi.org/10.5281/zenodo.21062977

