## Supplementary for "A chromosome-level assembly of an aquatic passerine bird, the northern white-throated dipper, *Cinclus cinclus cinclus* (Linnaeus, 1758)"

### Supplementary Material

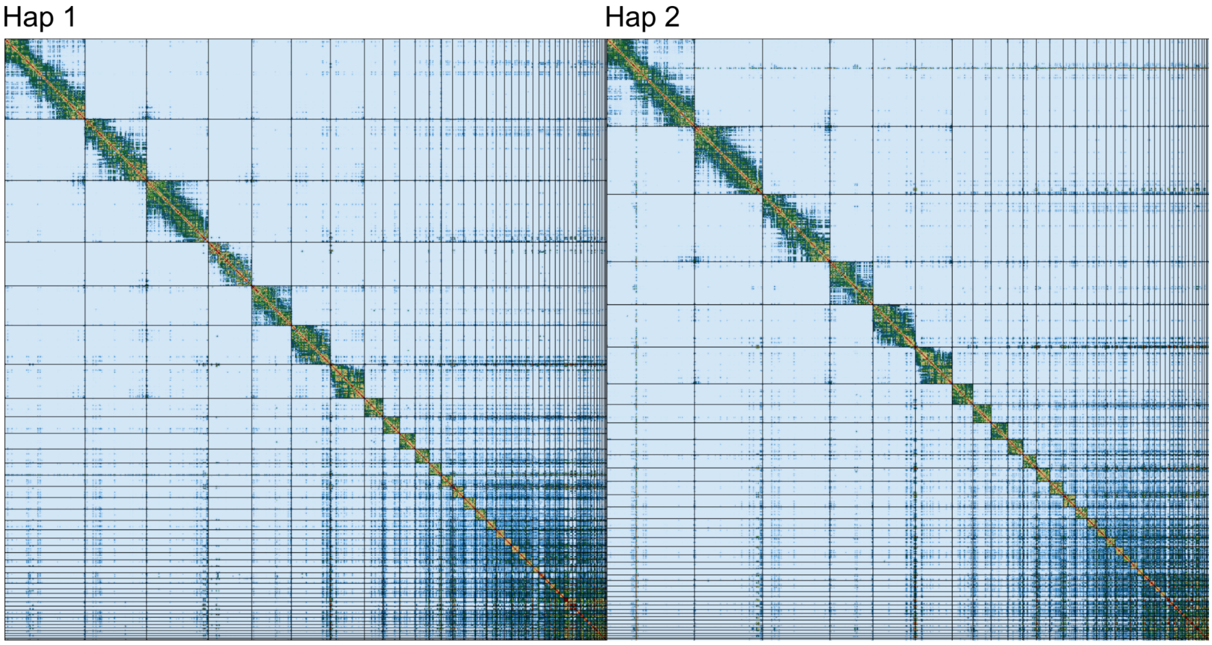


**Supplementary Figure 1:** Hi-C contact maps for *C. c. cinclus* hap1 and hap2. An elevated degree of interchromosomal interactivity (visualized as dark blue) can be seen for the microchromosomes.

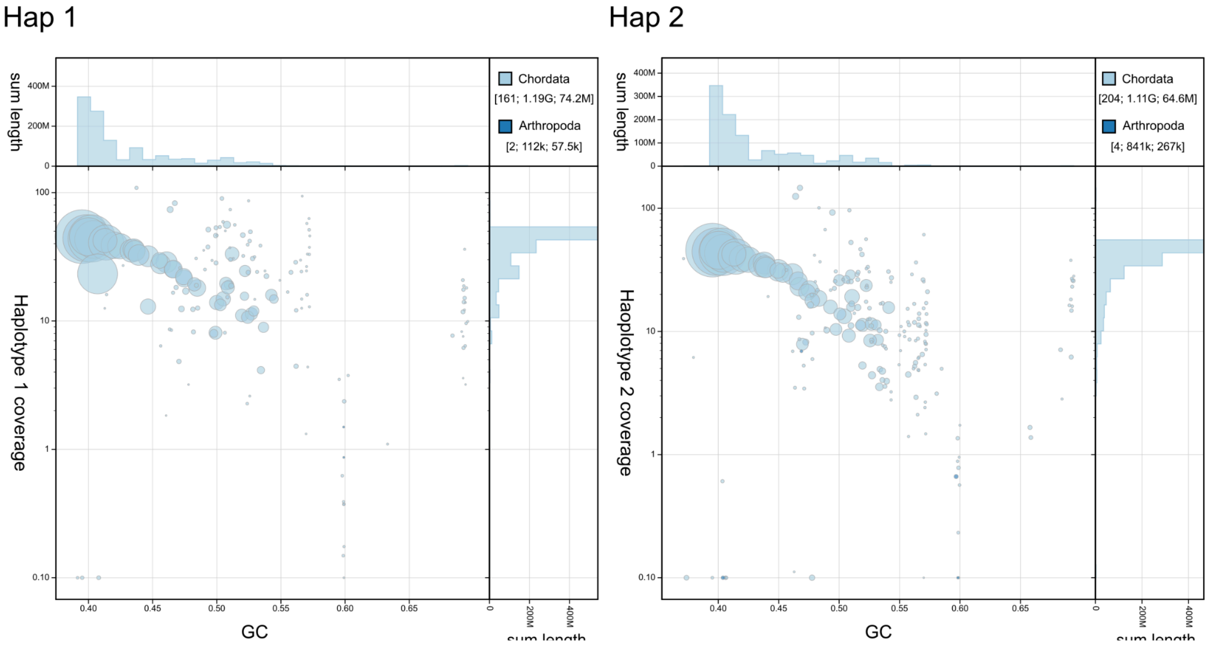


**Supplementary Figure 2. BlobToolKit GC-coverage plots of genome assemblies of *C. c. cinclus* hap1 and hap2.** The scaffolds are coloured by phylum. The size of the circles are in proportion to the length of the scaffolds. Histograms show the distribution of scaffold length sum along each axis.

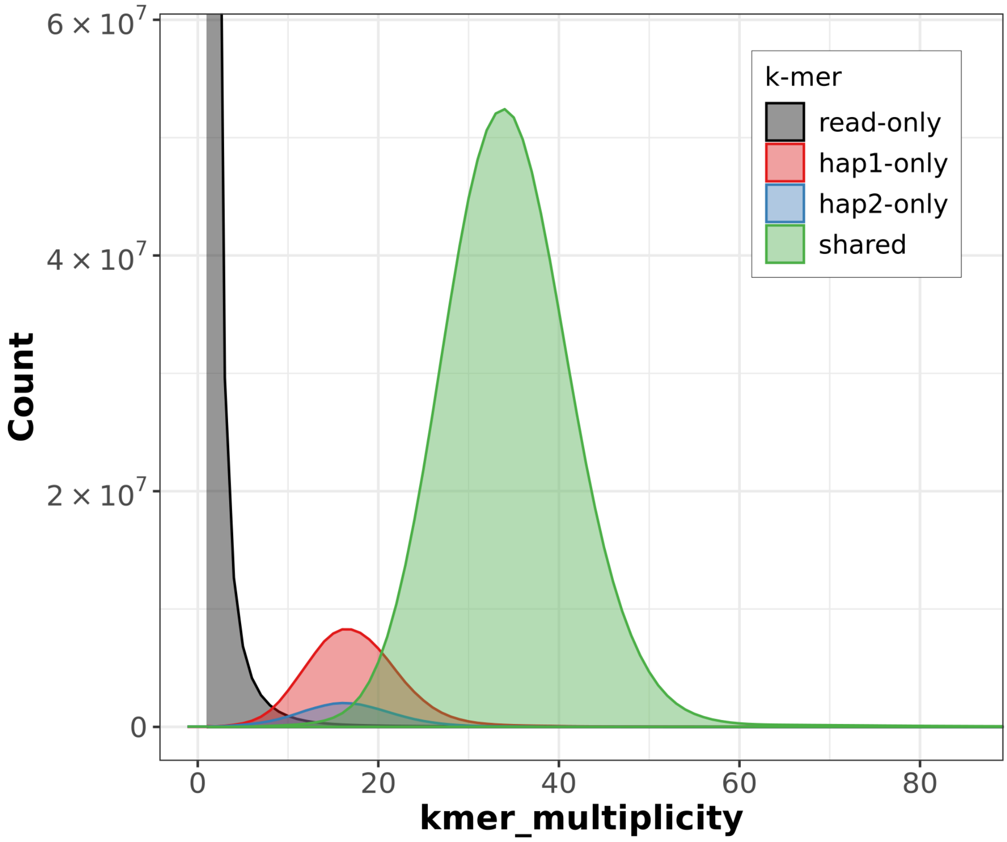


**Supplementary Figure 3. K-mer copy-number spectrum analysis of *C. c. cinclus* compared to k-mers from a database of ONT reads.** Assembly-specific k-mers are shown in red and blue, while k-mers shared by both pseudo-haplotypes are in green. The stack above 0 on the x-axis shows k-mers found in the assemblies, but not in the reads. The figure was generated by Merqury.

**Supplementary Table 1.** Comparisons of *C. c. cinclus* pseudo-haplotypes using minimap2 and paftools. Hap2 aligned to hap1.

| **Comparison** | | |
|---|---|---|
| Bases in alignment: 1,008,931,604 | | |
| Substitutions: 1,143,264 (0.11%) | | |
| **Size** | **Deletions** | **Insertions** |
| 1 bp | 66,941 | 66,349 |
| 2 bp | 18,919 | 18,806 |
| 3-49 bp | 47,703 | 47,480 |
| 50-999 bp | 9,392 | 9,526 |
| ≥1000 bp | 1,079 | 1,119 |

**Supplementary Table 2.** Assembly continuity and BUSCO completeness of the bCinCin5.1 assemblies and bCinCin1.1 primary reference. Assembly statistics were calculated with gfastats. BUSCO completeness was assessed against the 8,338 orthologs in aves_odb10 using BUSCO v5.8.3. Length statistics are reported in Mb.

| Measures | ONT hap1 | ONT hap2 | HiFi hap1 | HiFi hap2 | HiFi+ONT hap1 | HiFi+ONT hap2 | bCinCin1.1 pri |
|---|---|---|---|---|---|---|---|
| # scaffolds | 163 | 208 | 1717 | 628 | 566 | 907 | 277 |
| Total scaffold length (Mb) | 1186.40 | 1114.87 | 1131.69 | 903.42 | 996.68 | 1043.71 | 1170.84 |
| Average scaffold length (Mb) | 7.28 | 5.36 | 0.66 | 1.44 | 1.76 | 1.15 | 4.23 |
| Scaffold N50 (Mb) | 74.25 | 64.61 | 60.74 | 67.59 | 66.61 | 61.65 | 74.36 |
| Scaffold auN (Mb) | 69.23 | 67.78 | 64.13 | 71.33 | 68.79 | 65.38 | 69.97 |
| Scaffold L50 | 6 | 6 | 6 | 5 | 5 | 6 | 6 |
| Largest scaffold (Mb) | 153.12 | 153.56 | 150.29 | 146.95 | 144.37 | 150.35 | 152.36 |
| # contigs | 215 | 256 | 2740 | 1452 | 1135 | 1441 | 935 |
| Total contig length (Mb) | 1180.98 | 1113.27 | 1127.43 | 883.09 | 957.48 | 1038.27 | 1170.71 |
| Average contig length (Mb) | 5.49 | 4.35 | 0.41 | 0.61 | 0.84 | 0.72 | 1.25 |
| Contig N50 (Mb) | 35.76 | 32.03 | 6.07 | 7.54 | 8.89 | 8.95 | 3.56 |
| Contig auN (Mb) | 44.94 | 55.13 | 11.15 | 12.31 | 14.40 | 16.36 | 4.05 |
| Contig L50 | 9 | 8 | 40 | 27 | 25 | 25 | 103 |
| Largest contig (Mb) | 110.66 | 152.33 | 39.40 | 39.07 | 41.95 | 72.79 | 10.38 |
| # gaps in scaffolds | 52 | 48 | 1025 | 827 | 575 | 538 | 658 |
| Total gap length in scaffolds (Mb) | 5.42 | 1.60 | 4.26 | 20.34 | 39.20 | 5.43 | 0.13 |
| GC content % | 42.84 | 43.18 | 42.24 | 41.00 | 40.90 | 42.36 | 42.82 |
| Complete BUSCOs (%) | 99.2 | 95.0 | 96.7 | 79.8 | 80.6 | 93.1 | 99.1 |
| Single-copy BUSCOs (%) | 98.9 | 94.8 | 96.4 | 79.6 | 80.2 | 92.9 | 98.9 |
| Duplicated BUSCOs (%) | 0.3 | 0.2 | 0.3 | 0.2 | 0.5 | 0.1 | 0.3 |
| Fragmented BUSCOs (%) | 0.3 | 0.3 | 0.5 | 1.0 | 0.9 | 0.4 | 0.3 |
| Missing BUSCOs (%) | 0.5 | 4.7 | 2.8 | 19.2 | 18.5 | 6.5 | 0.6 |

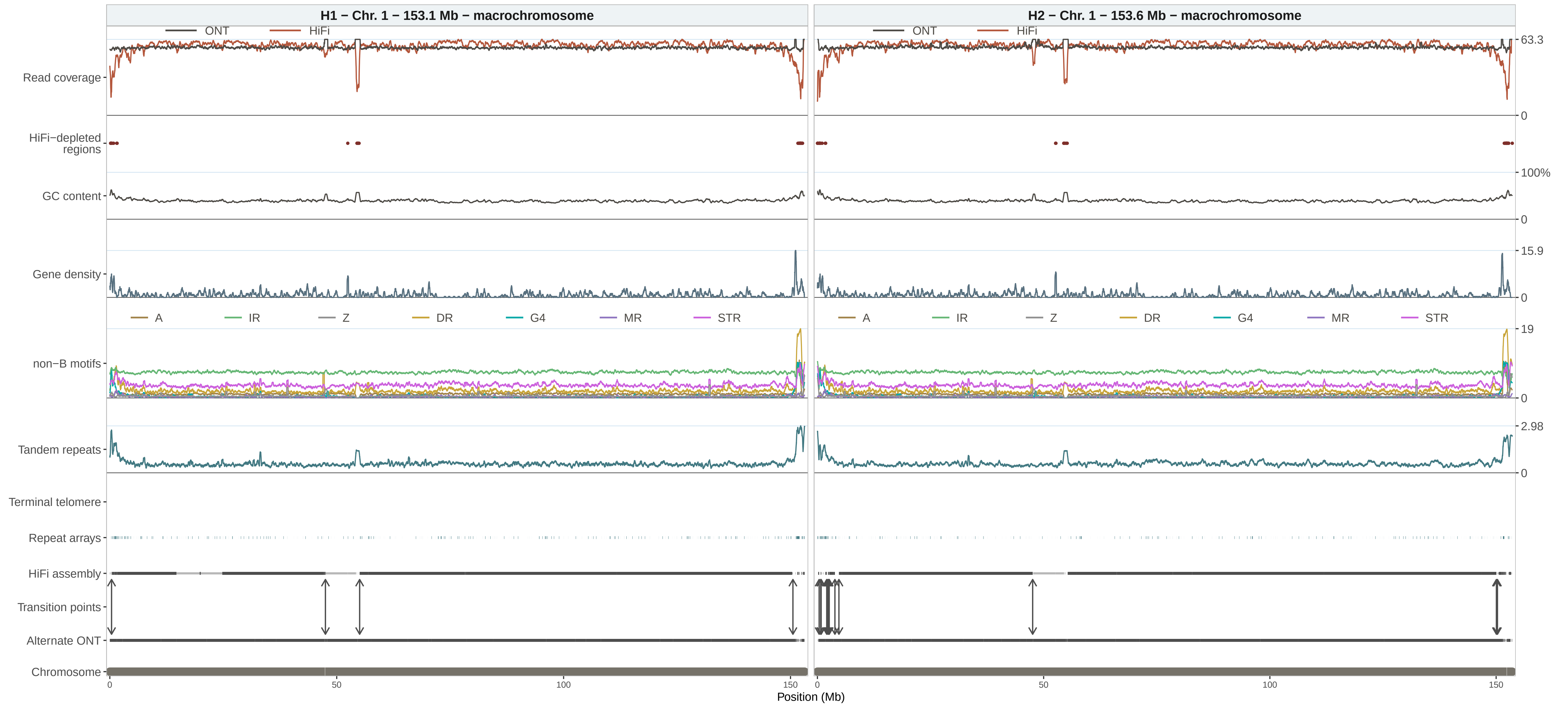

H1 − Chr. 1 − 153.1 Mb − macrochromosome
H2 − Chr. 1 − 153.6 Mb − macrochromosome
ONT
HiFi
Read coverage
63.3
0
HiFi−depleted regions
GC content
100%
0
Gene density
15.9
0
A
IR
Z
DR
G4
MR
STR
non−B motifs
19
0
Tandem repeats
2.98
0
Terminal telomere
Repeat arrays
HiFi assembly
Transition points
Alternate ONT
Chromosome
0
50
100
150
Position (Mb)

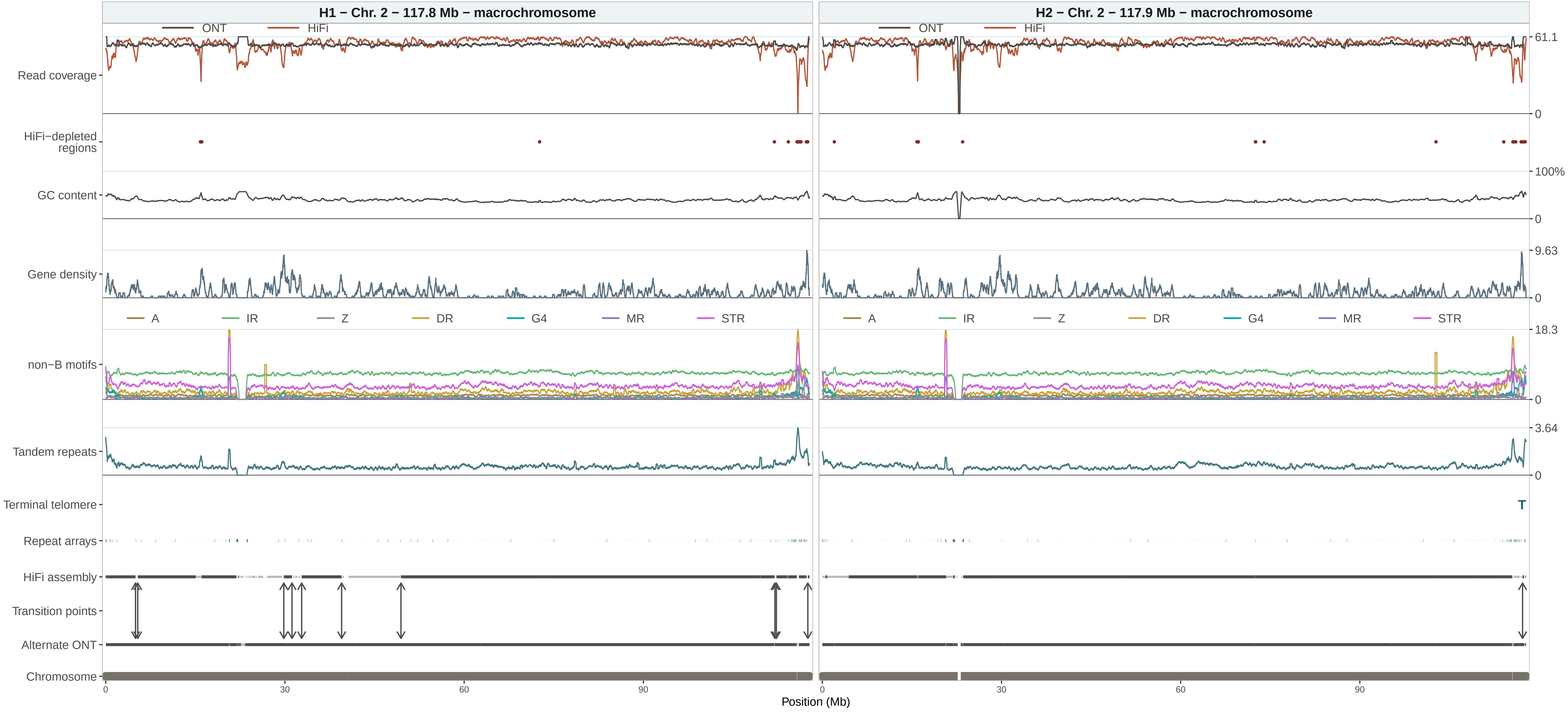

H1 − Chr. 2 − 117.8 Mb − macrochromosome
H2 − Chr. 2 − 117.9 Mb − macrochromosome
ONT
HiFi
Read coverage
HiFi−depleted regions
GC content
Gene density
non−B motifs
A
IR
Z
DR
G4
MR
STR
Tandem repeats
Terminal telomere
T
Repeat arrays
HiFi assembly
Transition points
Alternate ONT
Chromosome
61.1
0
100%
0
9.63
0
18.3
0
3.64
0
0
30
60
90
Position (Mb)

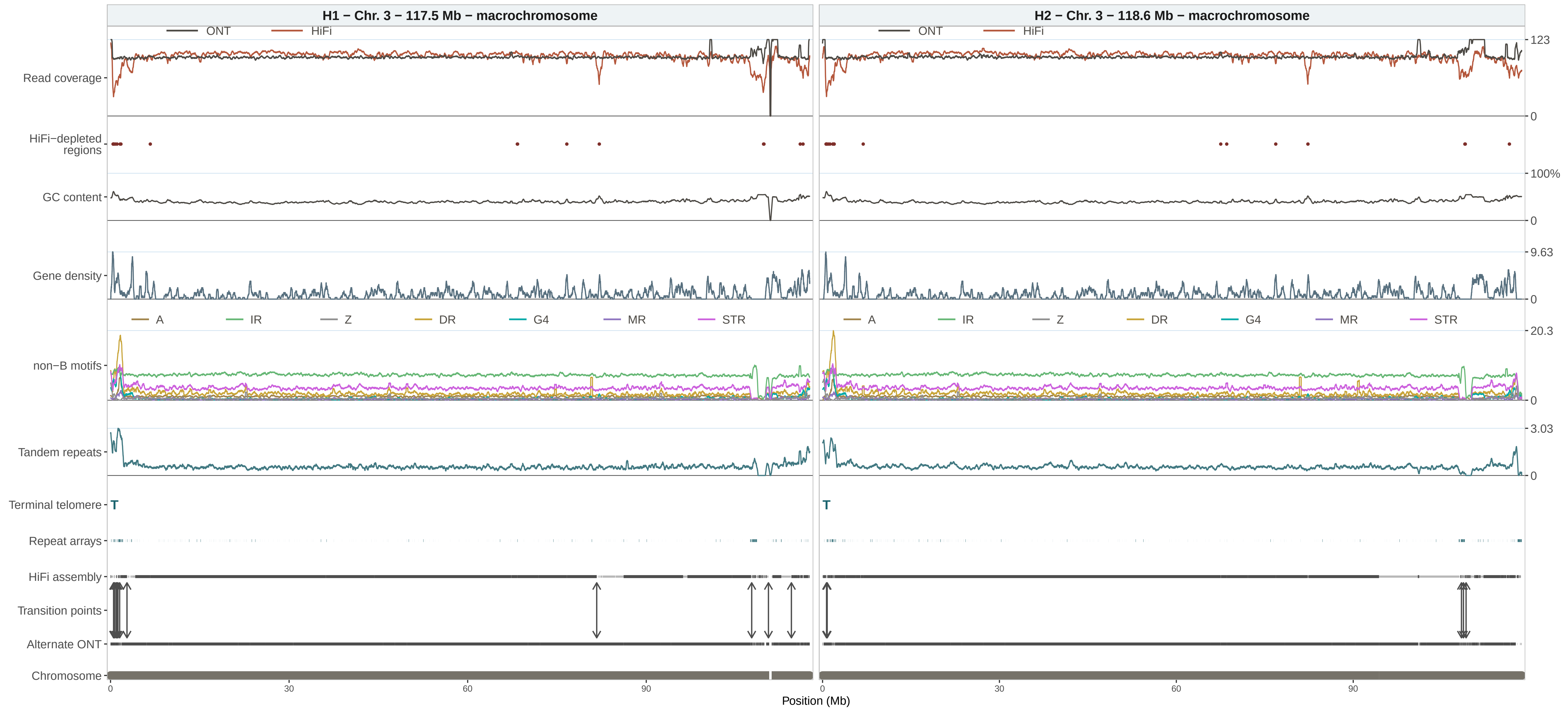

H1 – Chr. 3 – 117.5 Mb – macrochromosome
H2 – Chr. 3 – 118.6 Mb – macrochromosome
ONT
HiFi
Read coverage
HiFi−depleted regions
GC content
Gene density
non−B motifs
A
IR
Z
DR
G4
MR
STR
Tandem repeats
Terminal telomere
T
Repeat arrays
HiFi assembly
Transition points
Alternate ONT
Chromosome
123
0
100%
0
9.63
0
20.3
0
3.03
0
0
30
60
90
Position (Mb)

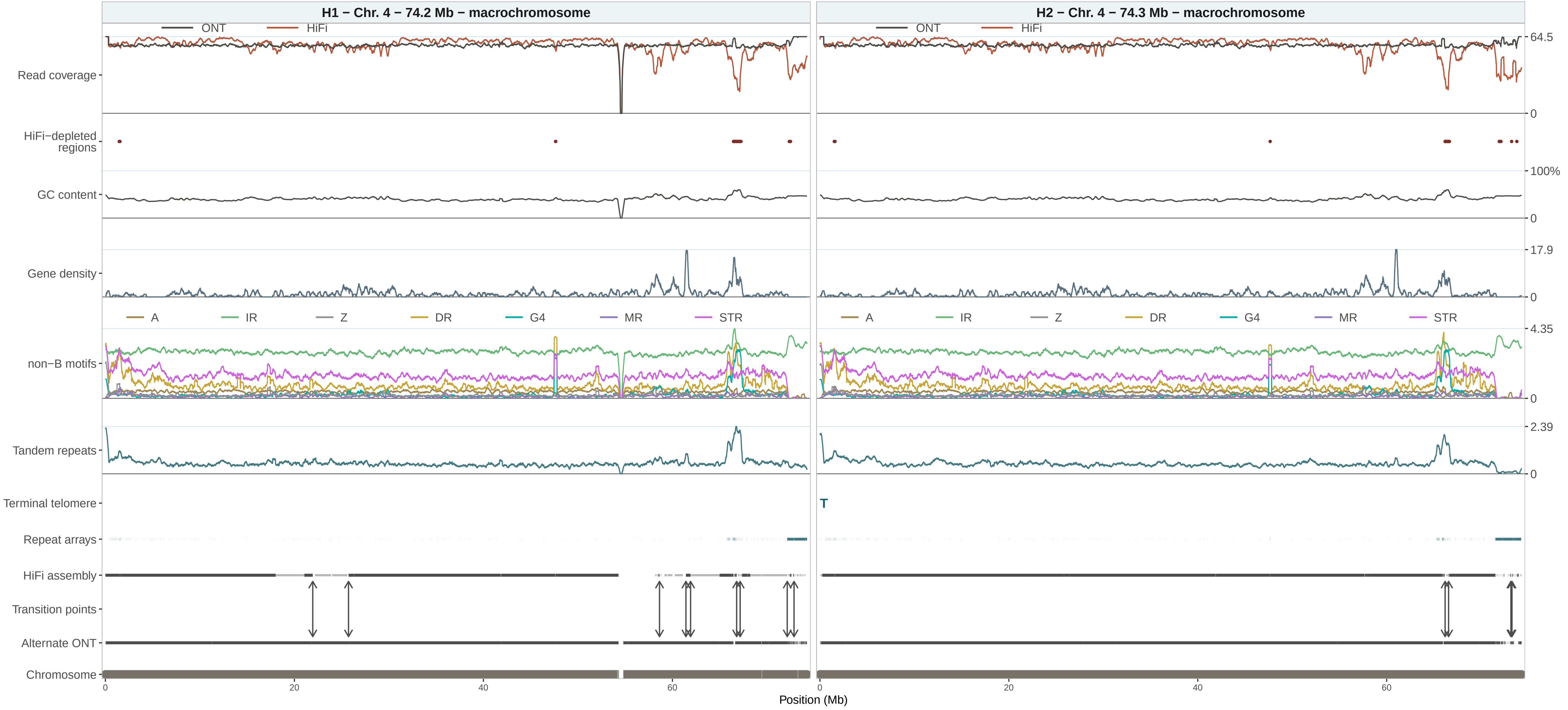

H1 − Chr. 4 − 74.2 Mb − macrochromosome
H2 − Chr. 4 − 74.3 Mb − macrochromosome
ONT
HiFi
Read coverage
64.5
0
HiFi−depleted regions
100%
GC content
0
17.9
Gene density
0
A
IR
Z
DR
G4
MR
STR
4.35
non−B motifs
0
2.39
Tandem repeats
0
Terminal telomere
T
Repeat arrays
HiFi assembly
Transition points
Alternate ONT
Chromosome
0
20
40
60
Position (Mb)

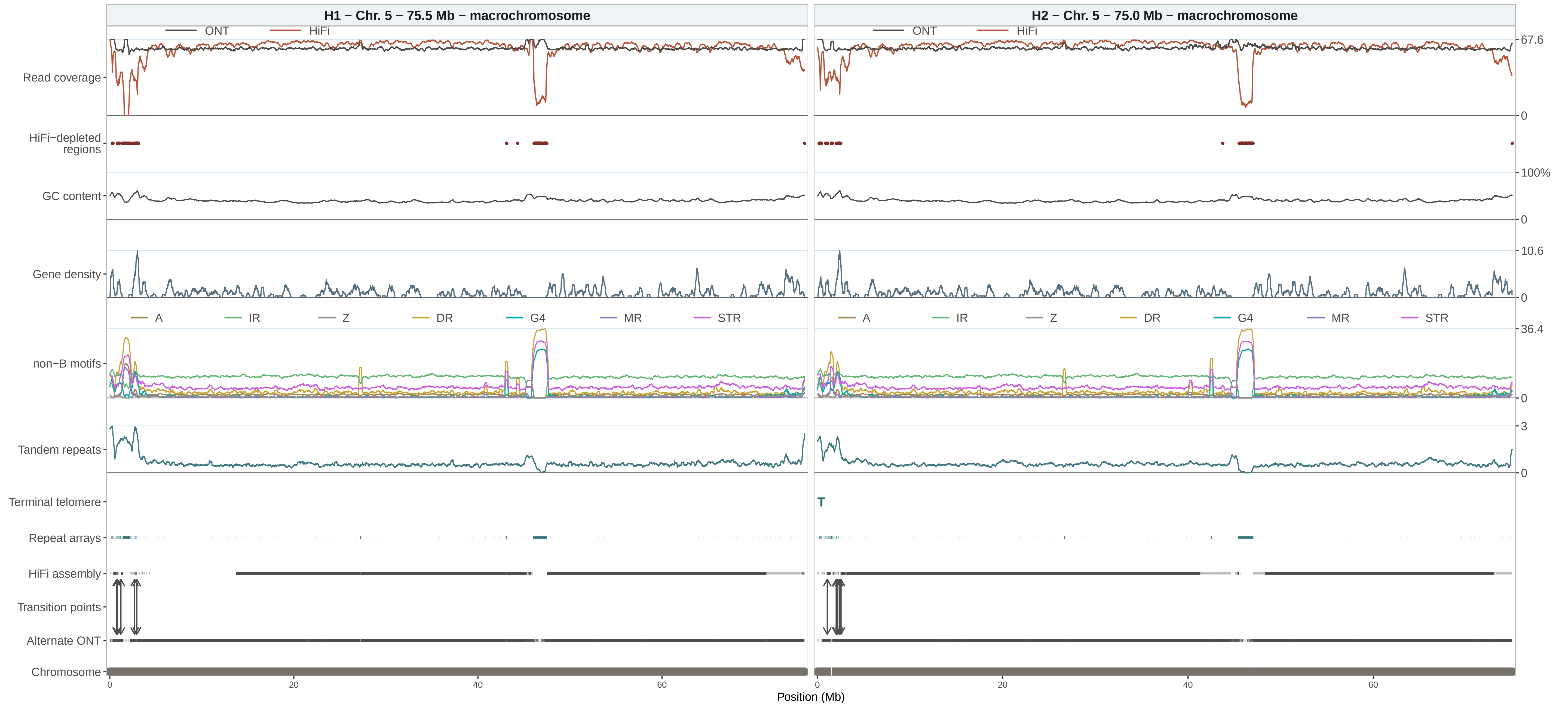

H1 − Chr. 5 − 75.5 Mb − macrochromosome
H2 − Chr. 5 − 75.0 Mb − macrochromosome
ONT
HiFi
Read coverage
HiFi−depleted regions
GC content
Gene density
non−B motifs
A
IR
Z
DR
G4
MR
STR
Tandem repeats
Terminal telomere
T
Repeat arrays
HiFi assembly
Transition points
Alternate ONT
Chromosome
67.6
0
100%
0
10.6
0
36.4
0
3
0
0
20
40
60
Position (Mb)

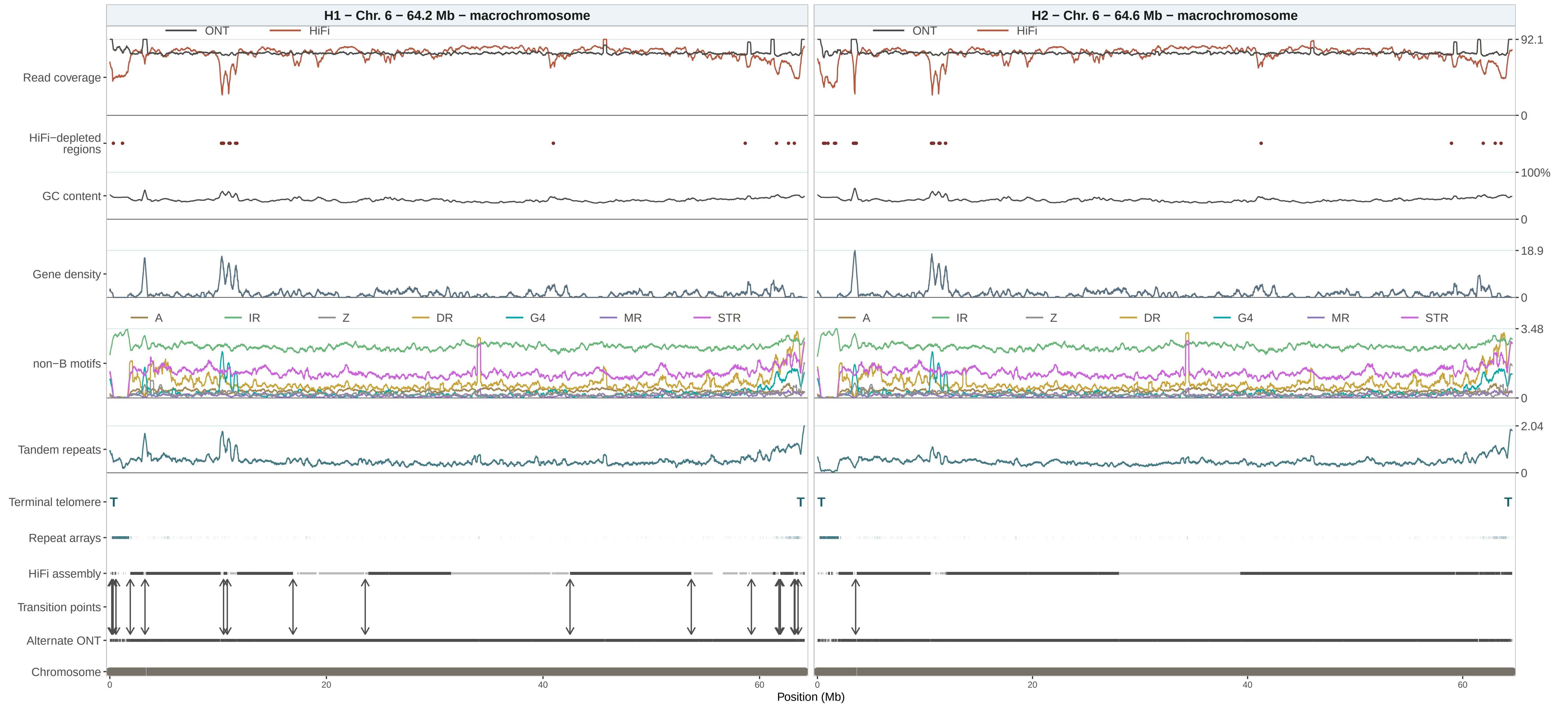
H1 – Chr. 6 – 64.2 Mb – macrochromosome
H2 – Chr. 6 – 64.6 Mb – macrochromosome
ONT
HiFi
Read coverage
HiFi−depleted regions
GC content
Gene density
non−B motifs
A
IR
Z
DR
G4
MR
STR
Tandem repeats
Terminal telomere
T
Repeat arrays
HiFi assembly
Transition points
Alternate ONT
Chromosome
92.1
0
100%
0
18.9
0
3.48
0
2.04
0
0
20
40
60
Position (Mb)

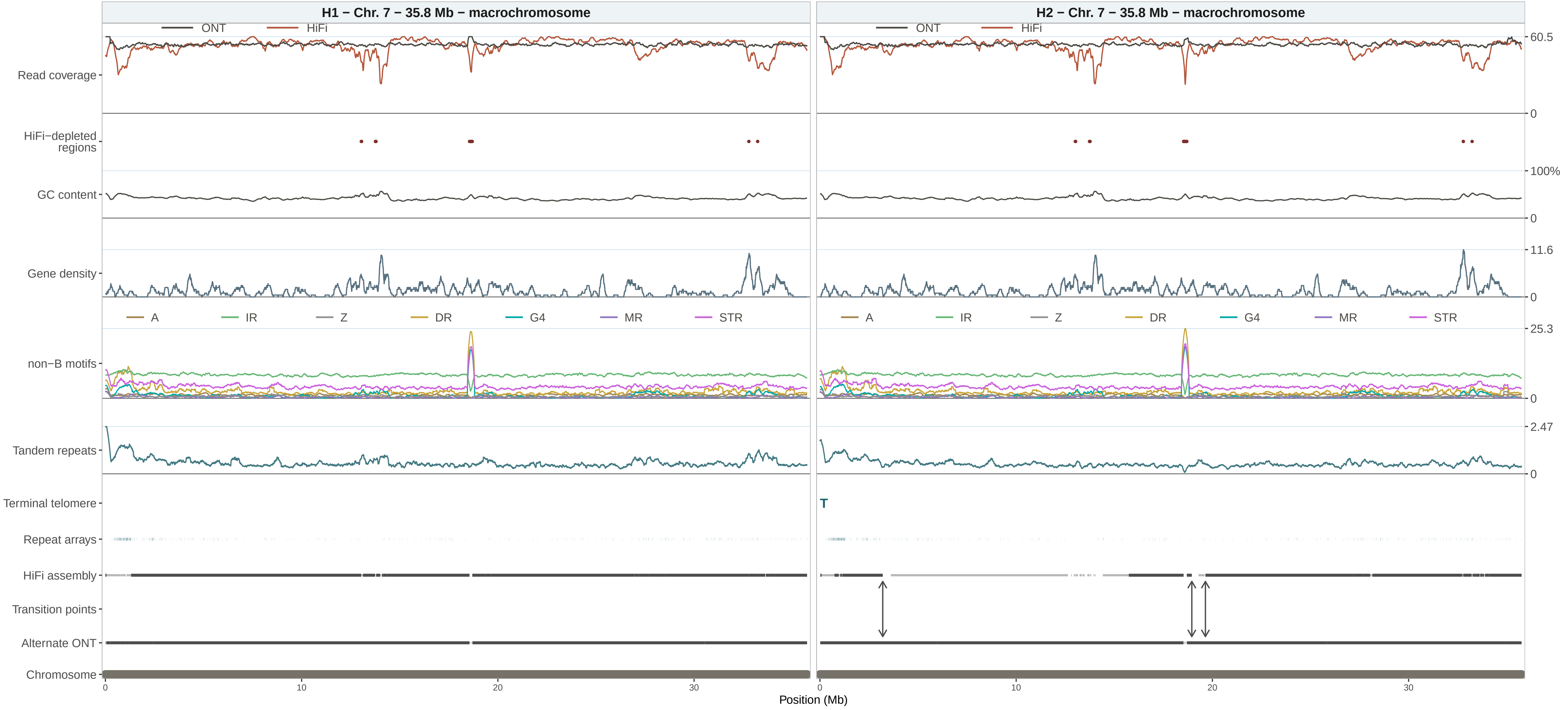

H1 − Chr. 7 − 35.8 Mb − macrochromosome
H2 − Chr. 7 − 35.8 Mb − macrochromosome
ONT
HiFi
Read coverage
HiFi−depleted regions
GC content
Gene density
non−B motifs
A
IR
Z
DR
G4
MR
STR
Tandem repeats
Terminal telomere
T
Repeat arrays
HiFi assembly
Transition points
Alternate ONT
Chromosome
60.5
0
100%
0
11.6
0
25.3
0
2.47
0
0
10
20
30
Position (Mb)

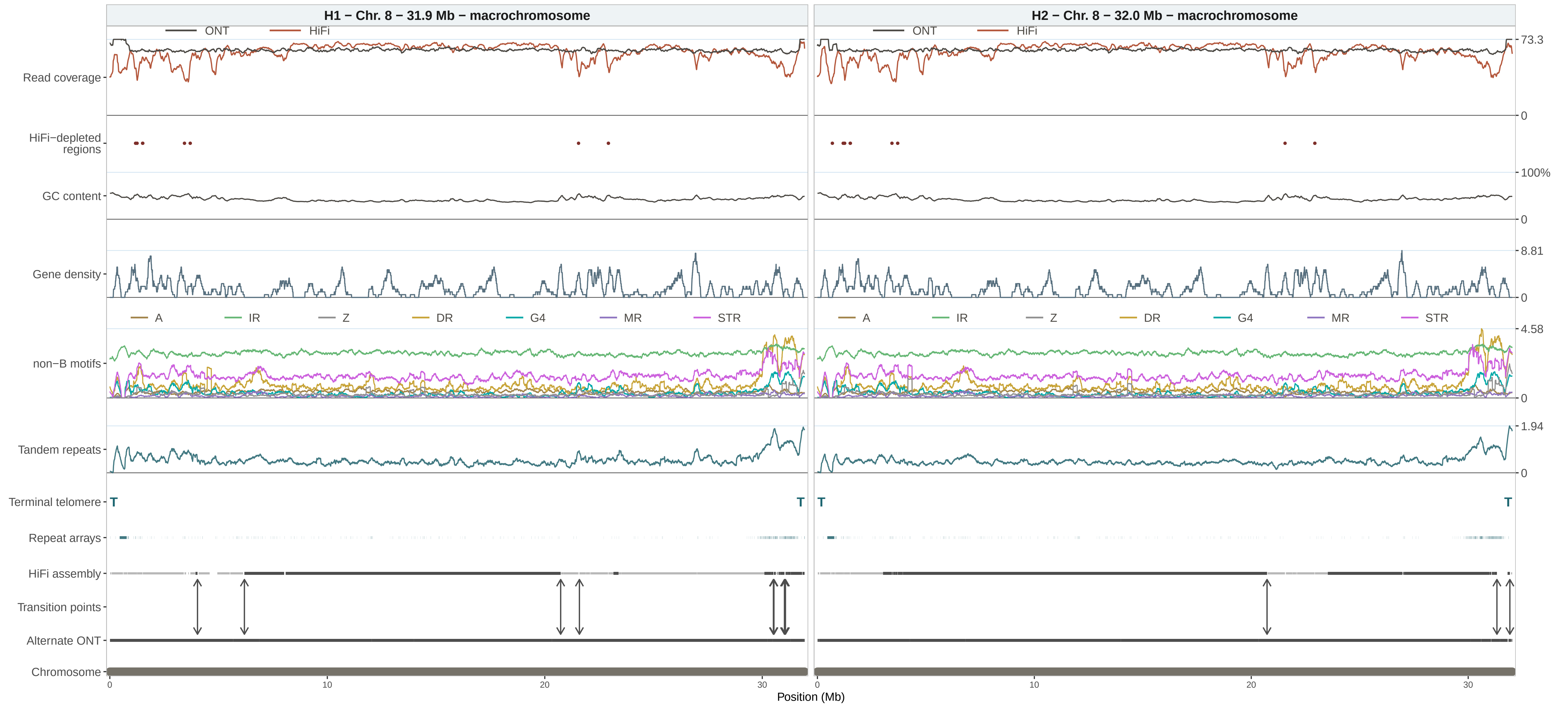

H1 – Chr. 8 – 31.9 Mb – macrochromosome
H2 – Chr. 8 – 32.0 Mb – macrochromosome
ONT
HiFi
Read coverage
HiFi–depleted regions
GC content
Gene density
non–B motifs
A
IR
Z
DR
G4
MR
STR
Tandem repeats
Terminal telomere
Repeat arrays
HiFi assembly
Transition points
Alternate ONT
Chromosome
T
73.3
0
100%
8.81
4.58
1.94
10
20
30
Position (Mb)

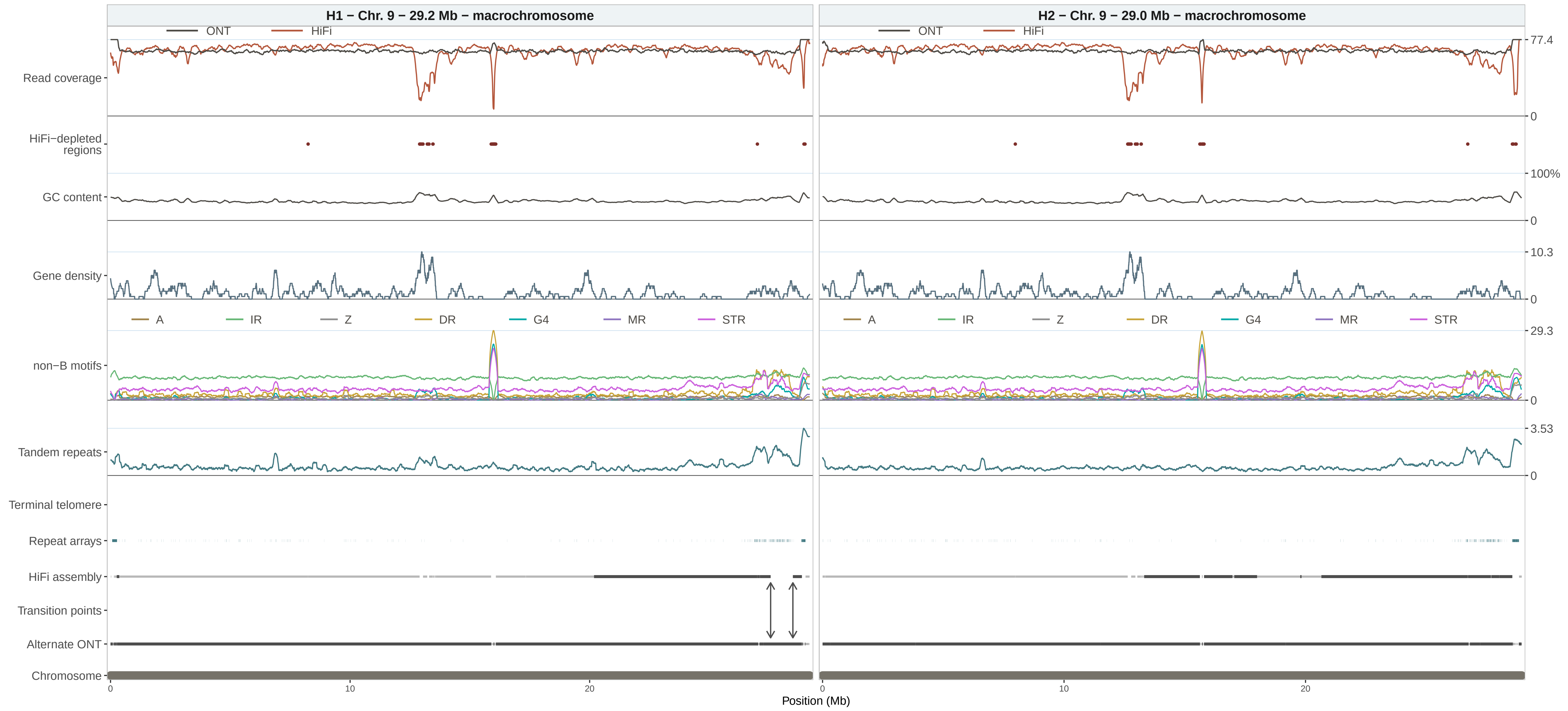

H1 − Chr. 9 − 29.2 Mb − macrochromosome
H2 − Chr. 9 − 29.0 Mb − macrochromosome
ONT
HiFi
Read coverage
HiFi−depleted regions
GC content
Gene density
non−B motifs
A
IR
Z
DR
G4
MR
STR
Tandem repeats
Terminal telomere
Repeat arrays
HiFi assembly
Transition points
Alternate ONT
Chromosome
77.4
0
100%
10.3
29.3
3.53
10
20
Position (Mb)

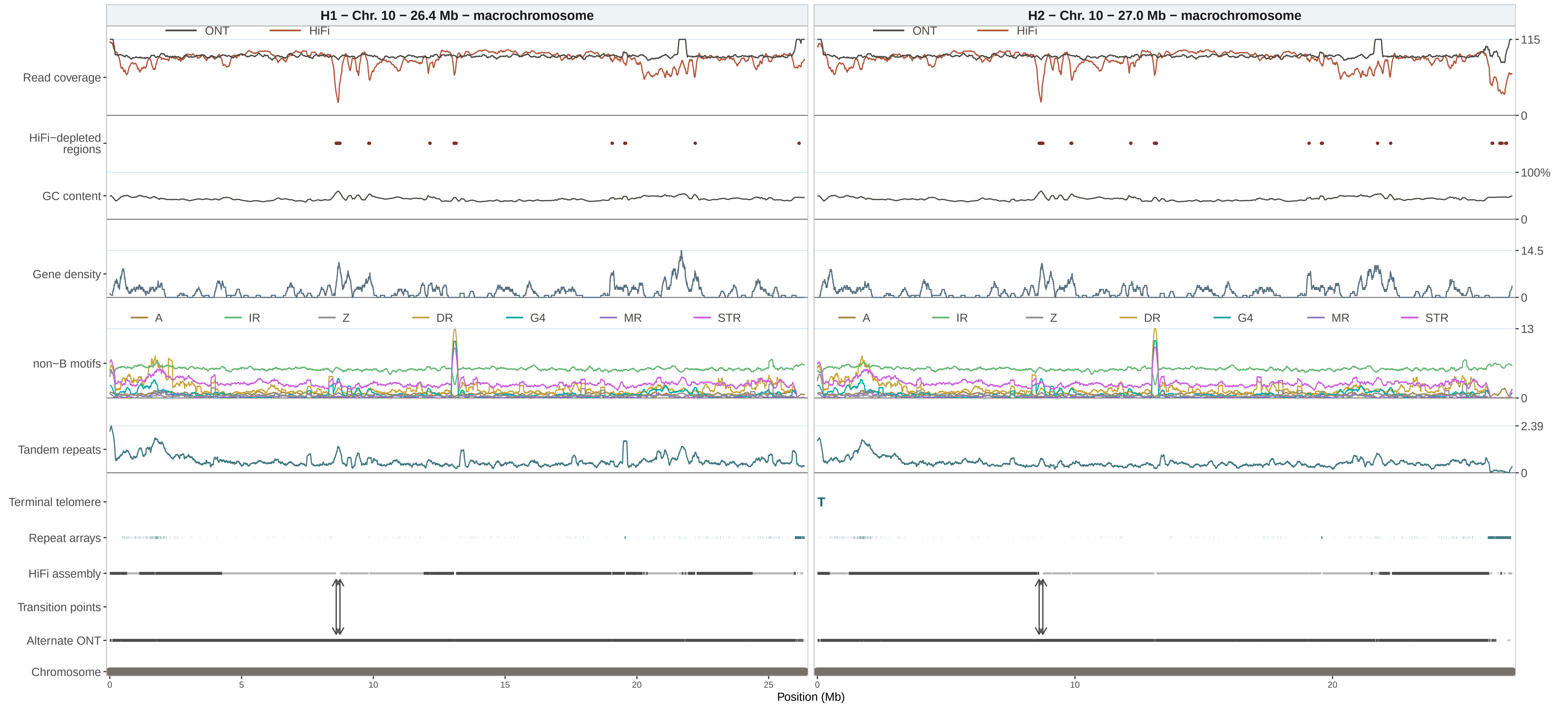
H1 − Chr. 10 − 26.4 Mb − macrochromosome
H2 − Chr. 10 − 27.0 Mb − macrochromosome
ONT
HiFi
Read coverage
HiFi−depleted regions
GC content
Gene density
non−B motifs
A
IR
Z
DR
G4
MR
STR
Tandem repeats
Terminal telomere
T
Repeat arrays
HiFi assembly
Transition points
Alternate ONT
Chromosome
115
0
100%
0
14.5
0
13
0
2.39
0
0
5
10
15
20
25
0
10
20
Position (Mb)

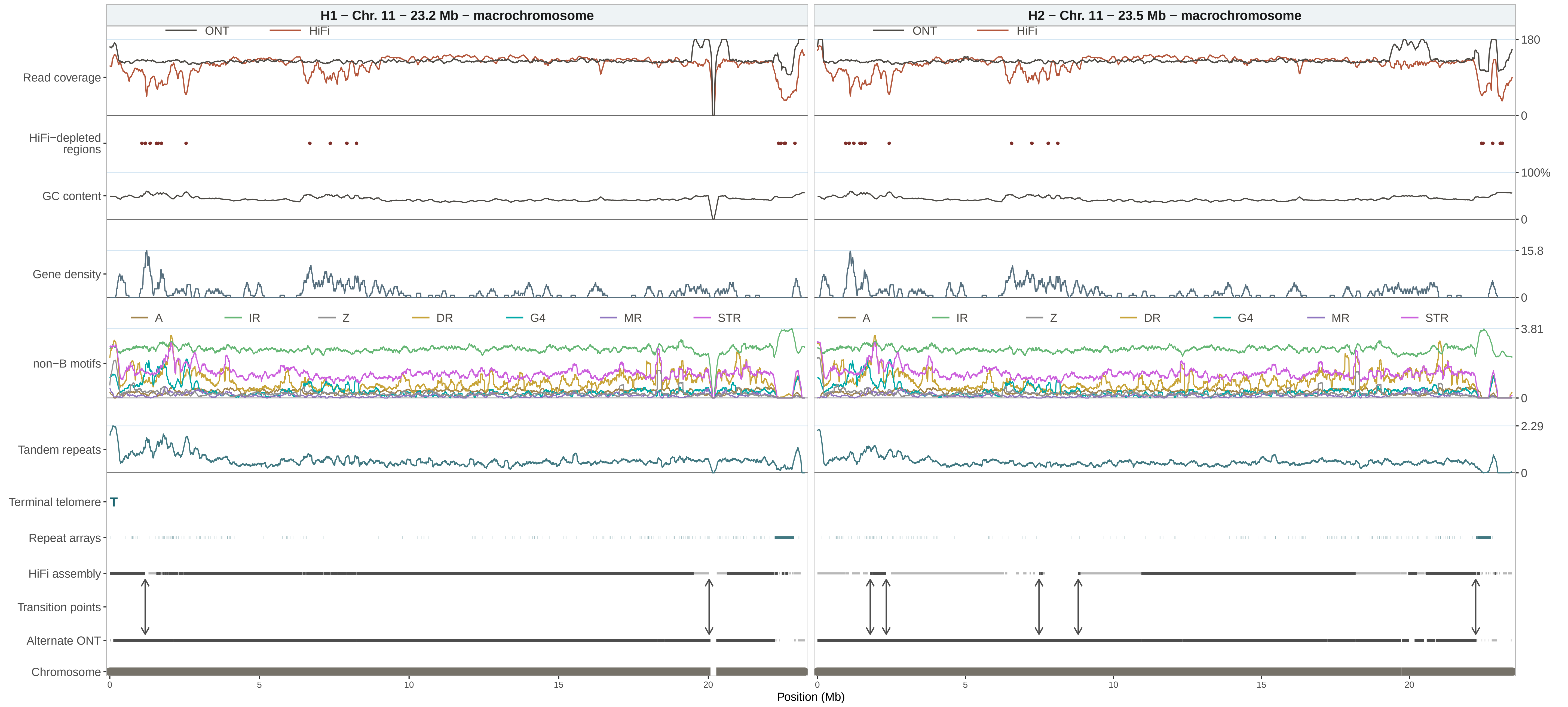

H1 − Chr. 11 − 23.2 Mb − macrochromosome
H2 − Chr. 11 − 23.5 Mb − macrochromosome
ONT
HiFi
Read coverage
HiFi−depleted regions
GC content
Gene density
non−B motifs
A
IR
Z
DR
G4
MR
STR
Tandem repeats
Terminal telomere
T
Repeat arrays
HiFi assembly
Transition points
Alternate ONT
Chromosome
180
0
100%
0
15.8
0
3.81
0
2.29
0
0
5
10
15
20
Position (Mb)

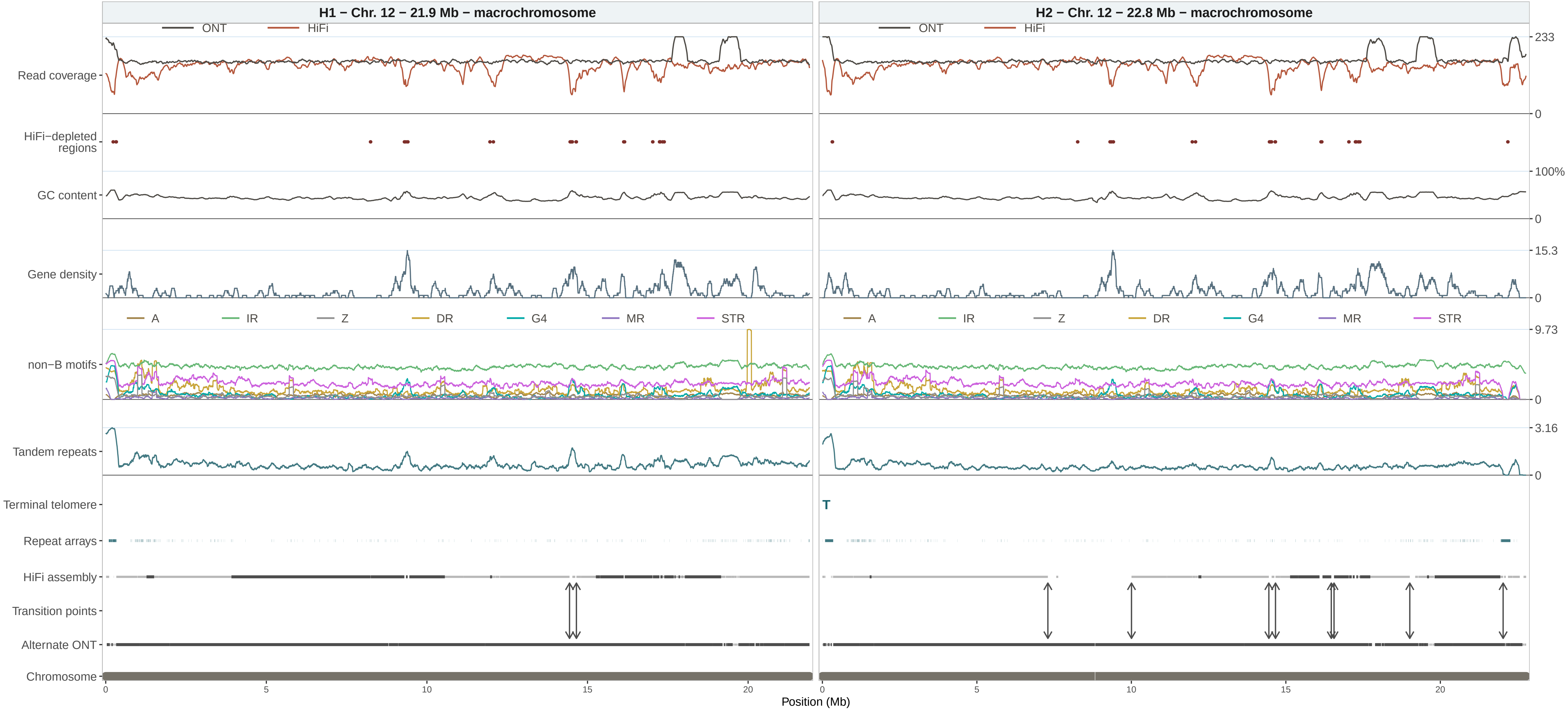

H1 − Chr. 12 − 21.9 Mb − macrochromosome
H2 − Chr. 12 − 22.8 Mb − macrochromosome
ONT
HiFi
Read coverage
HiFi−depleted regions
GC content
Gene density
non−B motifs
A
IR
Z
DR
G4
MR
STR
Tandem repeats
Terminal telomere
T
Repeat arrays
HiFi assembly
Transition points
Alternate ONT
Chromosome
233
0
100%
0
15.3
0
9.73
0
3.16
0
0
5
10
15
20
Position (Mb)

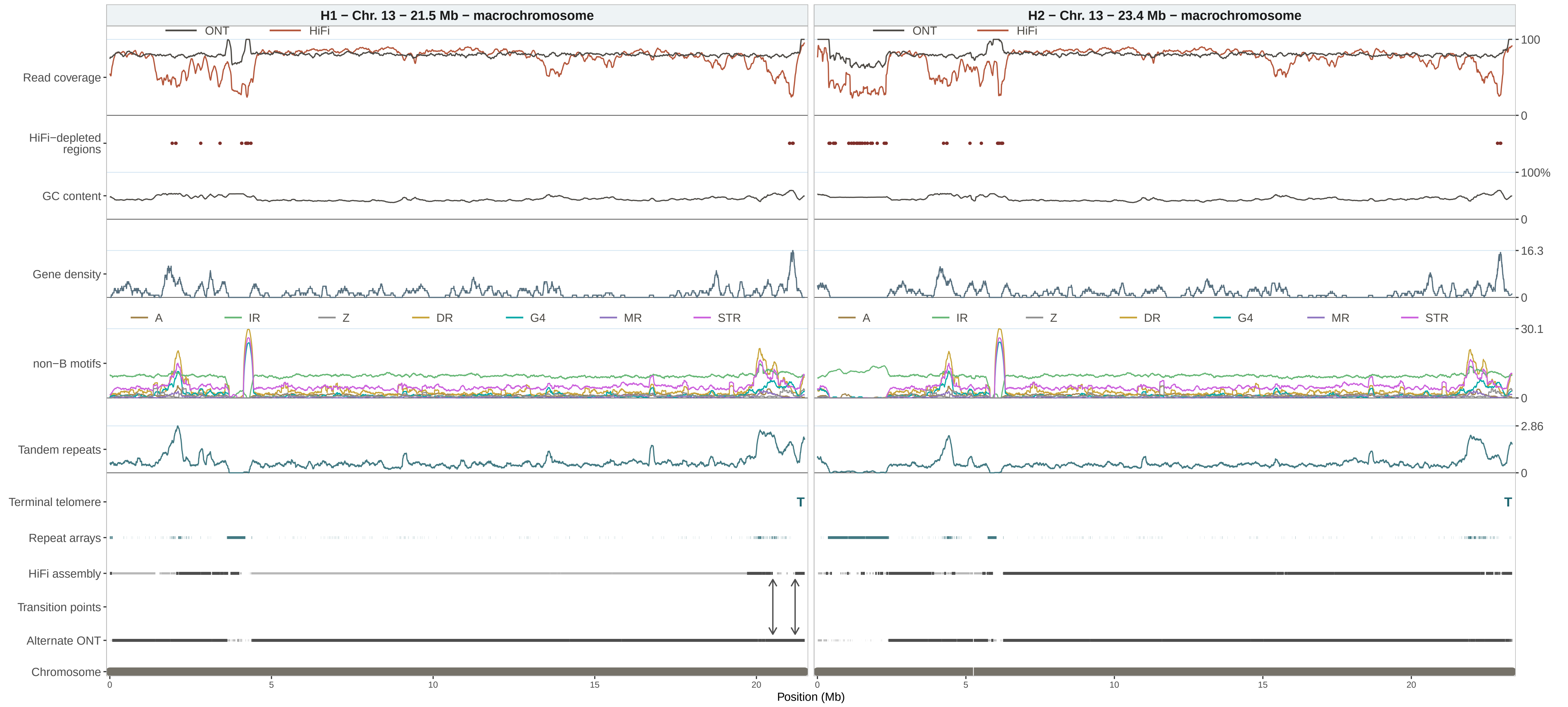

H1 − Chr. 13 − 21.5 Mb − macrochromosome
H2 − Chr. 13 − 23.4 Mb − macrochromosome
ONT
HiFi
Read coverage
HiFi−depleted regions
GC content
Gene density
non−B motifs
A
IR
Z
DR
G4
MR
STR
Tandem repeats
Terminal telomere
T
Repeat arrays
HiFi assembly
Transition points
Alternate ONT
Chromosome
100
0
100%
16.3
30.1
2.86
0
5
10
15
20
Position (Mb)

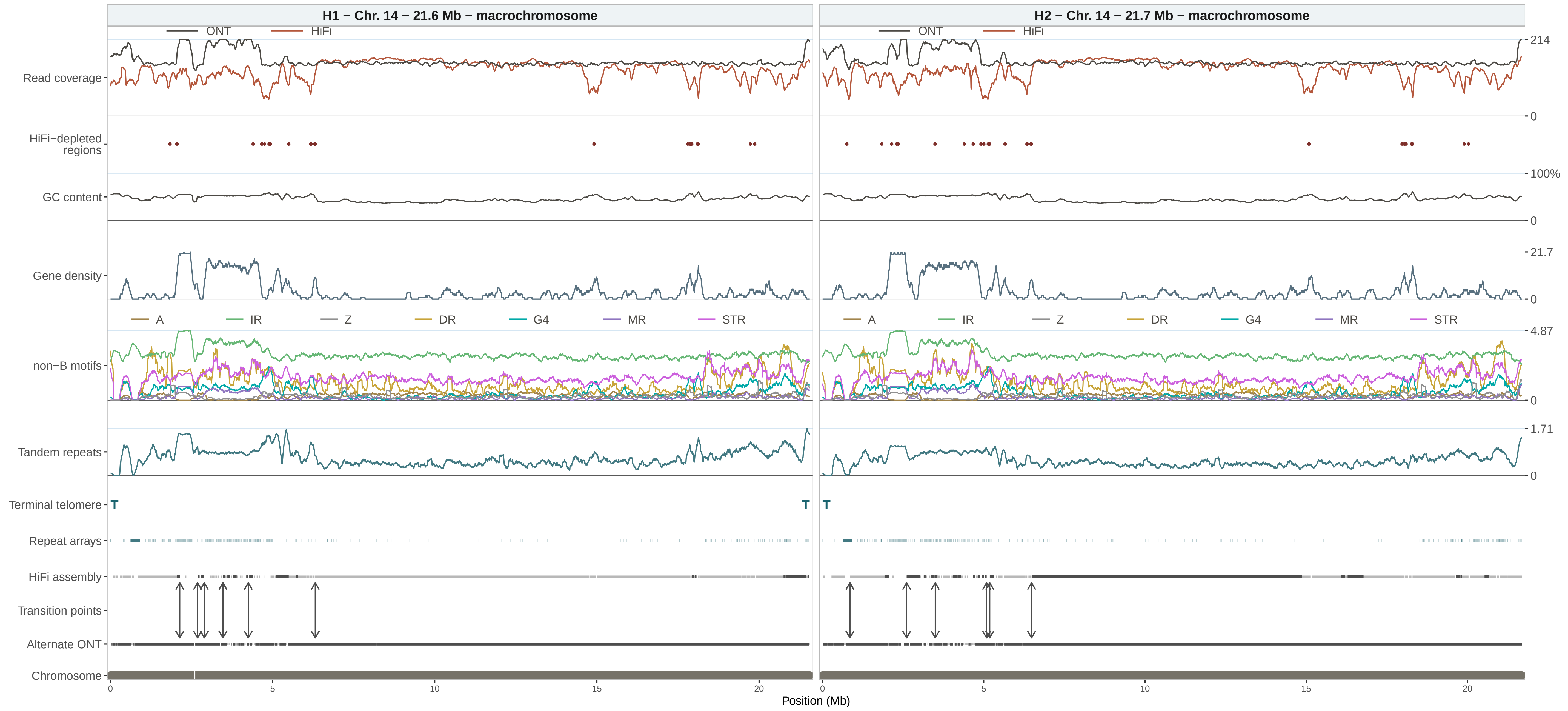

H1 − Chr. 14 − 21.6 Mb − macrochromosome
H2 − Chr. 14 − 21.7 Mb − macrochromosome
ONT
HiFi
Read coverage
HiFi−depleted regions
GC content
Gene density
non−B motifs
Tandem repeats
Terminal telomere
Repeat arrays
HiFi assembly
Transition points
Alternate ONT
Chromosome
A
IR
Z
DR
G4
MR
STR
214
0
100%
21.7
4.87
1.71
T
0
5
10
15
20
Position (Mb)

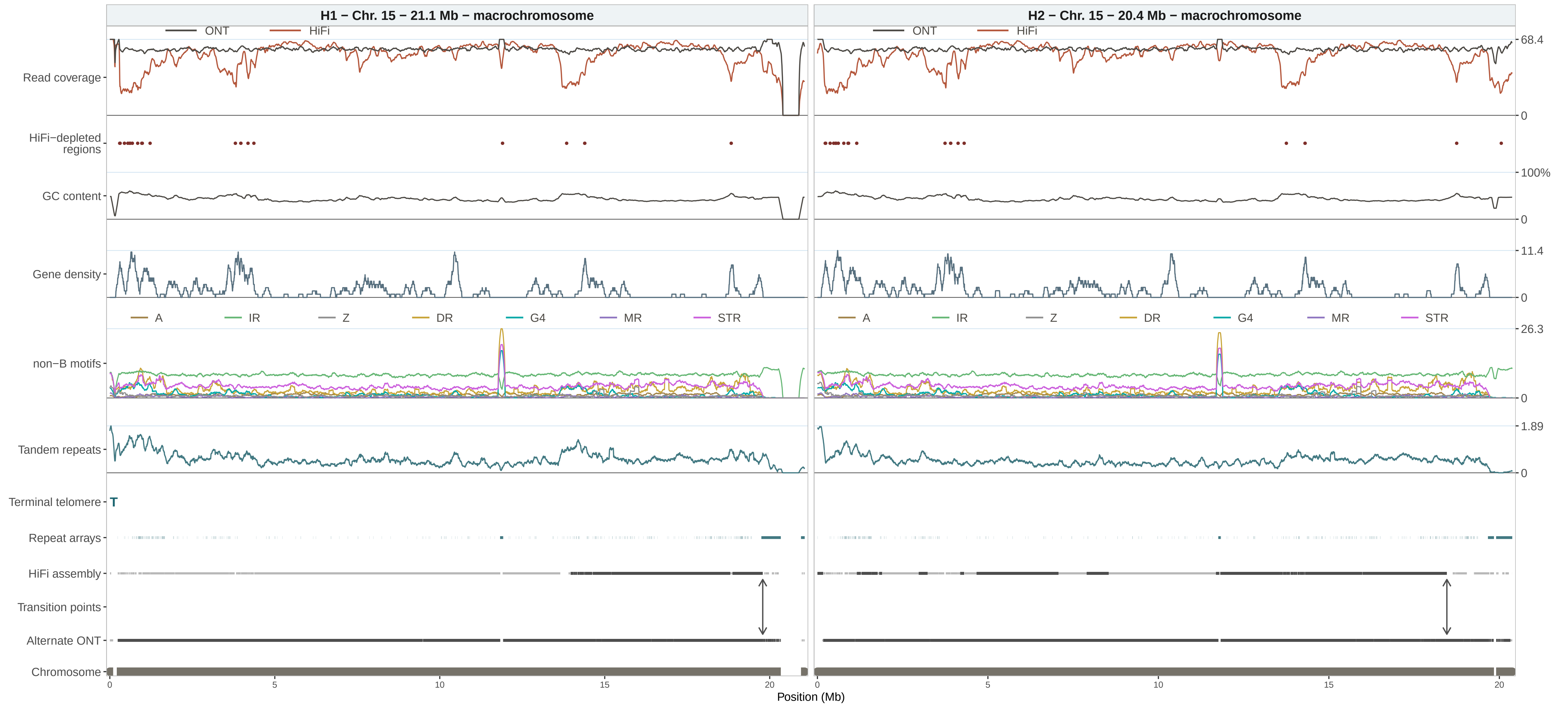

H1 − Chr. 15 − 21.1 Mb − macrochromosome
H2 − Chr. 15 − 20.4 Mb − macrochromosome
ONT
HiFi
Read coverage
HiFi−depleted regions
GC content
Gene density
non−B motifs
Tandem repeats
Terminal telomere
T
Repeat arrays
HiFi assembly
Transition points
Alternate ONT
Chromosome
A
IR
Z
DR
G4
MR
STR
68.4
0
100%
0
11.4
0
26.3
0
1.89
0
0
5
10
15
20
Position (Mb)

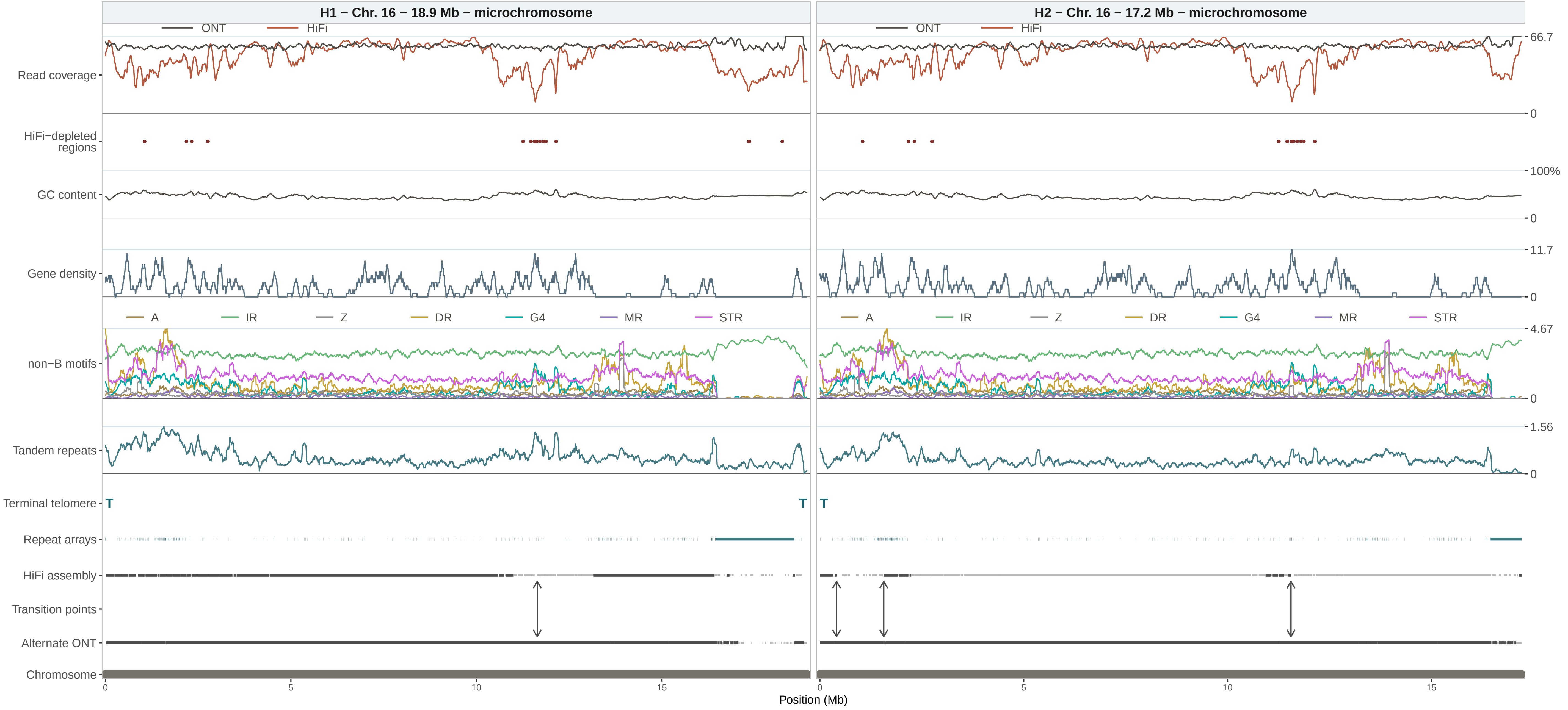
H1 − Chr. 16 − 18.9 Mb − microchromosome
H2 − Chr. 16 − 17.2 Mb − microchromosome
ONT
HiFi
Read coverage
HiFi−depleted regions
GC content
Gene density
non−B motifs
A
IR
Z
DR
G4
MR
STR
Tandem repeats
Terminal telomere
T
Repeat arrays
HiFi assembly
Transition points
Alternate ONT
Chromosome
66.7
0
100%
11.7
4.67
1.56
0
5
10
15
Position (Mb)

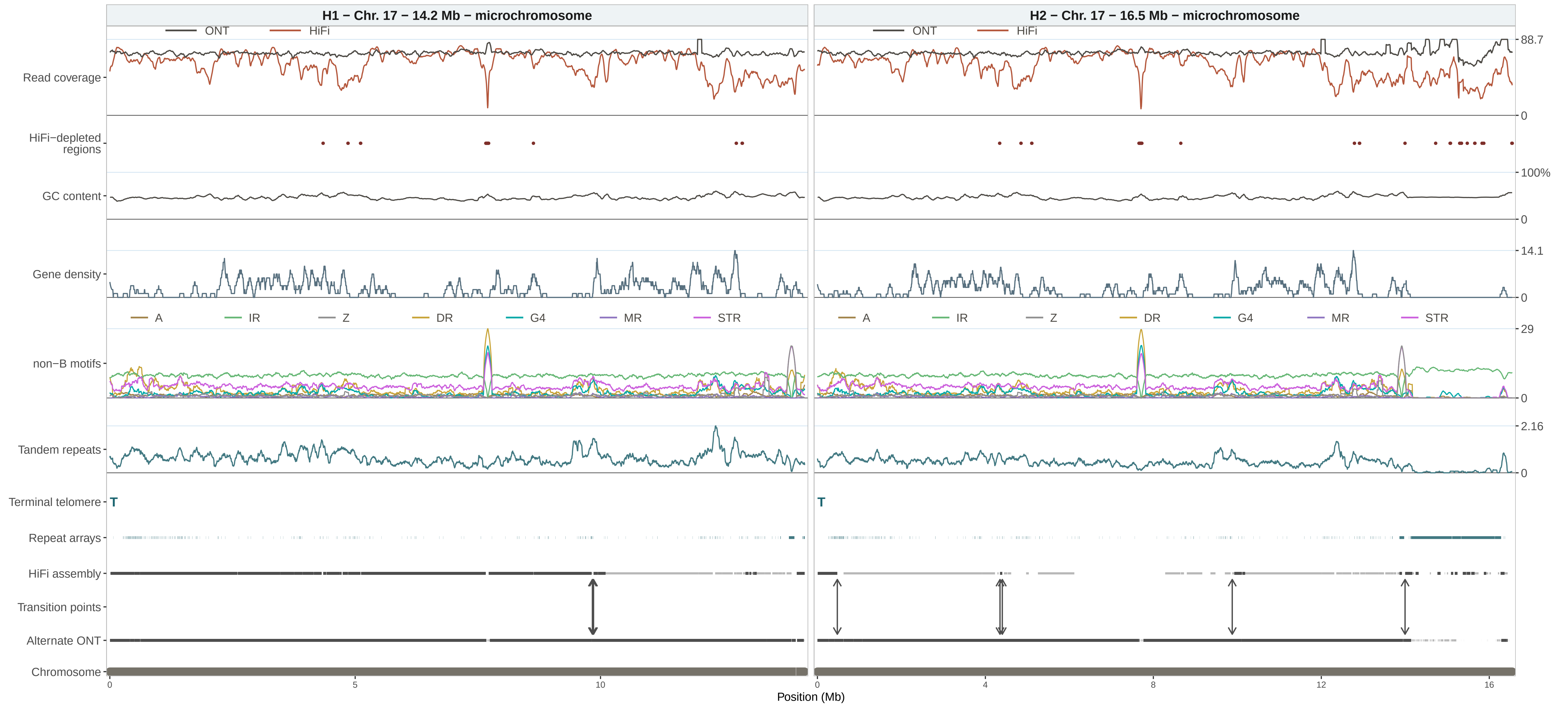
H1 – Chr. 17 – 14.2 Mb – microchromosome
H2 – Chr. 17 – 16.5 Mb – microchromosome
ONT
HiFi
Read coverage
HiFi−depleted regions
GC content
Gene density
non−B motifs
A
IR
Z
DR
G4
MR
STR
Tandem repeats
Terminal telomere
T
Repeat arrays
HiFi assembly
Transition points
Alternate ONT
Chromosome
88.7
0
100%
0
14.1
0
29
0
2.16
0
0
5
10
0
4
8
12
16
Position (Mb)

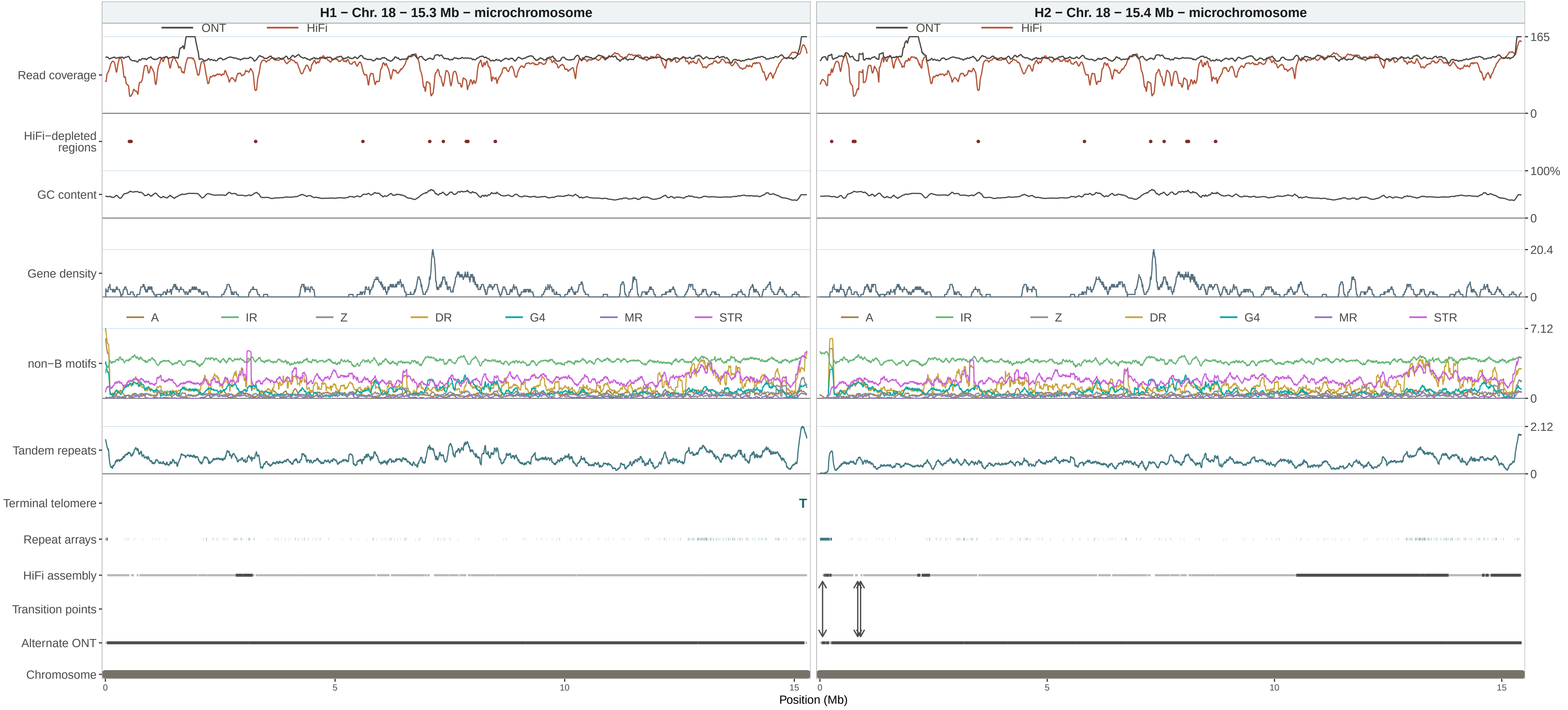

H1 − Chr. 18 − 15.3 Mb − microchromosome
H2 − Chr. 18 − 15.4 Mb − microchromosome
ONT
HiFi
Read coverage
HiFi−depleted regions
GC content
Gene density
non−B motifs
A
IR
Z
DR
G4
MR
STR
Tandem repeats
Terminal telomere
T
Repeat arrays
HiFi assembly
Transition points
Alternate ONT
Chromosome
165
0
100%
0
20.4
0
7.12
0
2.12
0
0
5
10
15
Position (Mb)

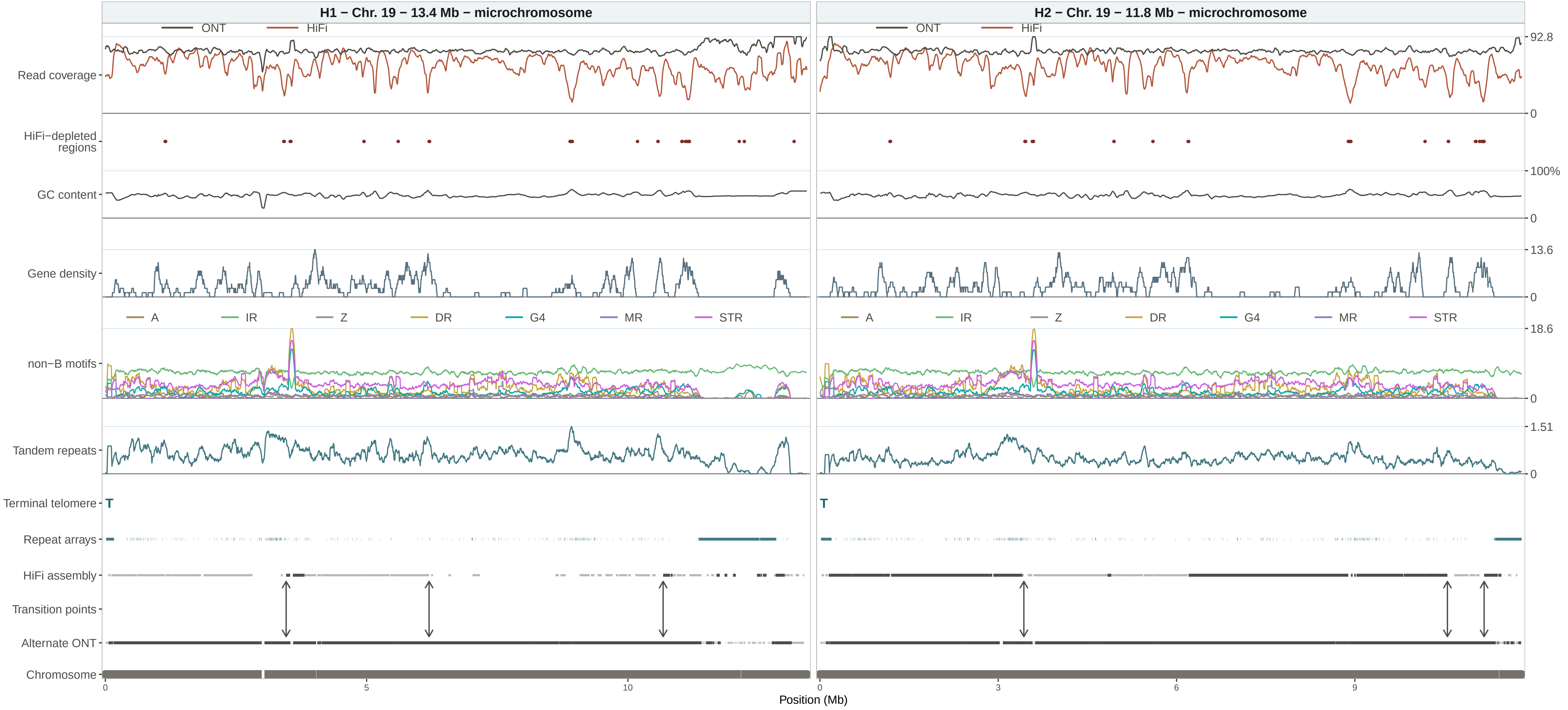

H1 – Chr. 19 – 13.4 Mb – microchromosome
H2 – Chr. 19 – 11.8 Mb – microchromosome
ONT
HiFi
Read coverage
HiFi–depleted regions
GC content
Gene density
A
IR
Z
DR
G4
MR
STR
non–B motifs
Tandem repeats
Terminal telomere
T
Repeat arrays
HiFi assembly
Transition points
Alternate ONT
Chromosome
92.8
0
100%
0
13.6
0
18.6
0
1.51
0
0
5
10
0
3
6
9
Position (Mb)

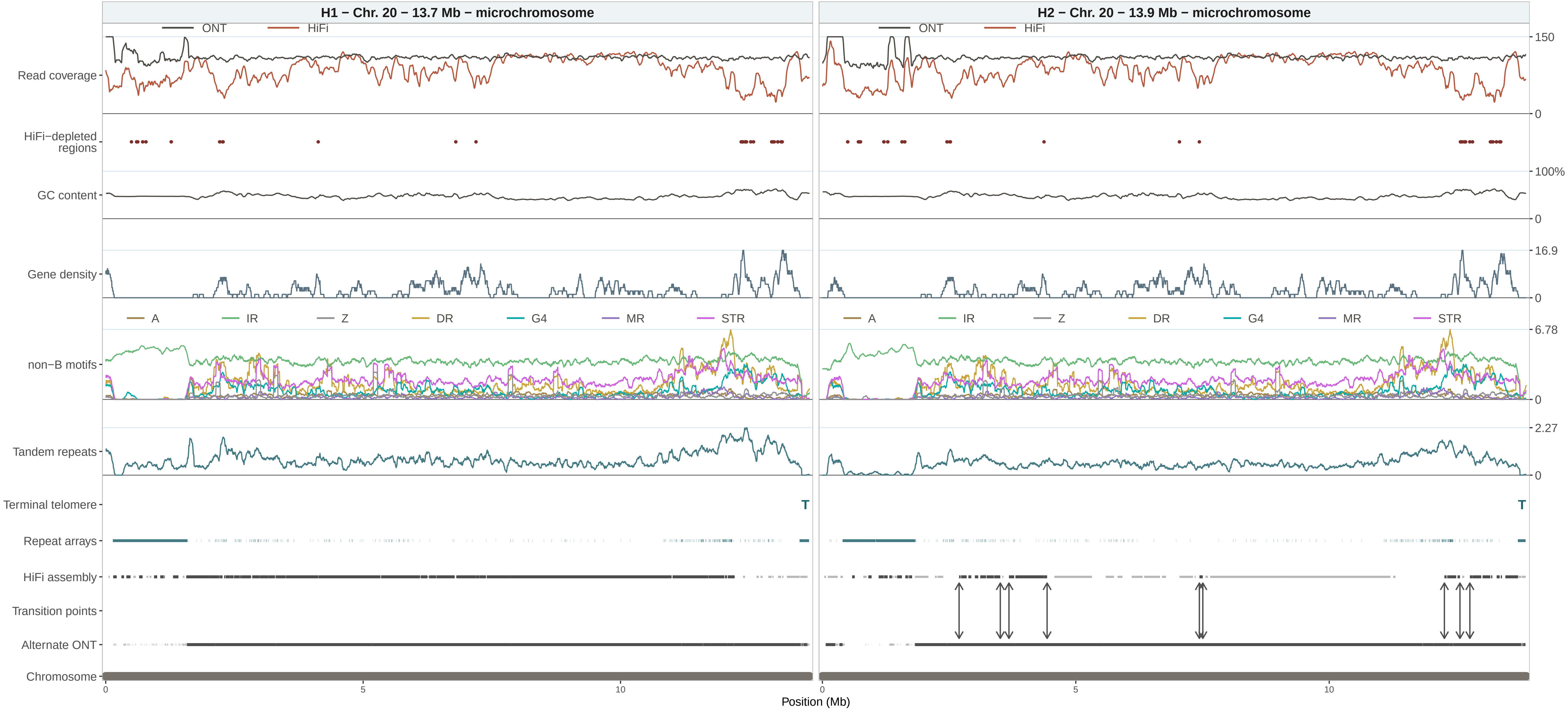

H1 – Chr. 20 – 13.7 Mb – microchromosome
H2 – Chr. 20 – 13.9 Mb – microchromosome
ONT
HiFi
Read coverage
HiFi−depleted regions
GC content
Gene density
non−B motifs
A
IR
Z
DR
G4
MR
STR
Tandem repeats
Terminal telomere
T
Repeat arrays
HiFi assembly
Transition points
Alternate ONT
Chromosome
150
0
100%
0
16.9
0
6.78
0
2.27
0
0
5
10
Position (Mb)

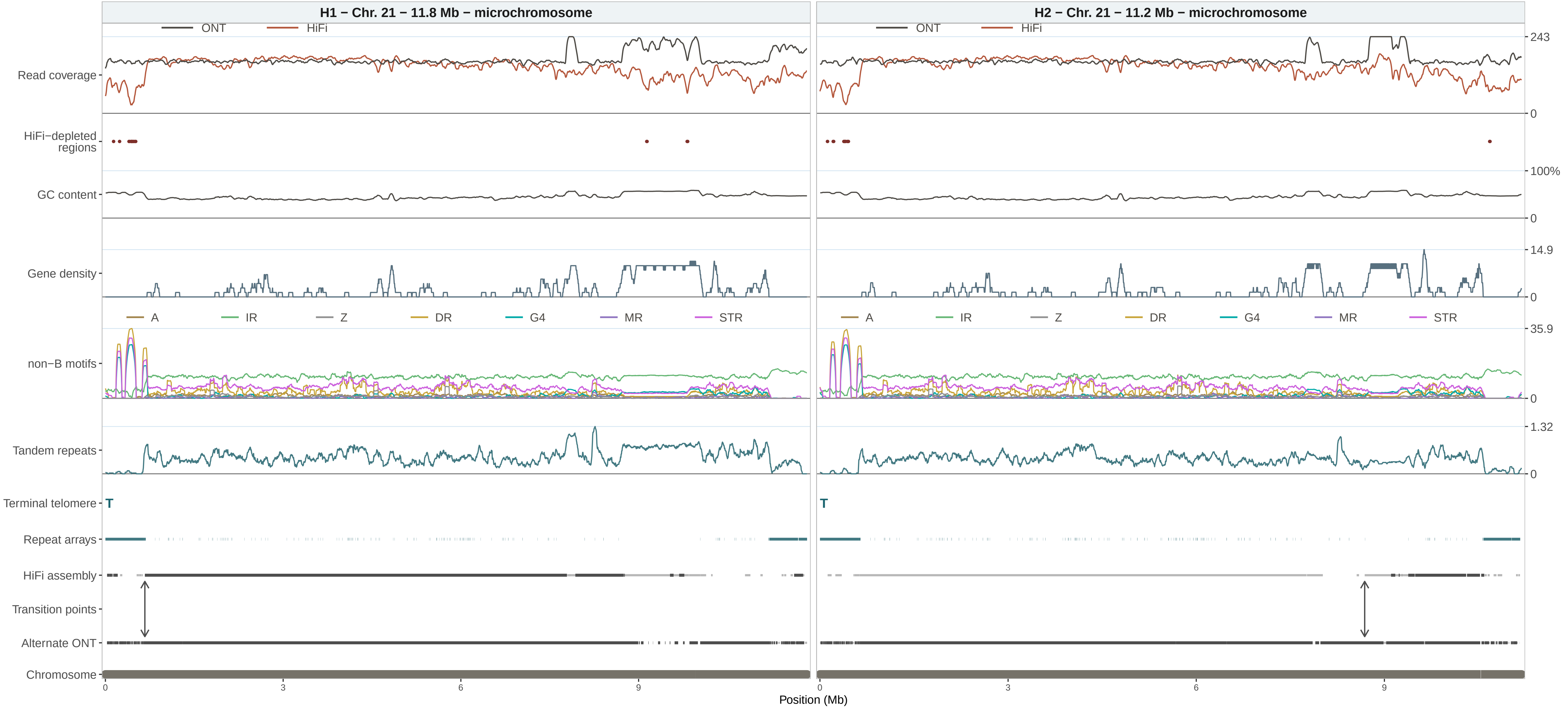

H1 − Chr. 21 − 11.8 Mb − microchromosome
H2 − Chr. 21 − 11.2 Mb − microchromosome
ONT
HiFi
Read coverage
HiFi−depleted regions
GC content
Gene density
non−B motifs
A
IR
Z
DR
G4
MR
STR
Tandem repeats
Terminal telomere
T
Repeat arrays
HiFi assembly
Transition points
Alternate ONT
Chromosome
243
0
100%
14.9
35.9
1.32
0
3
6
9
Position (Mb)

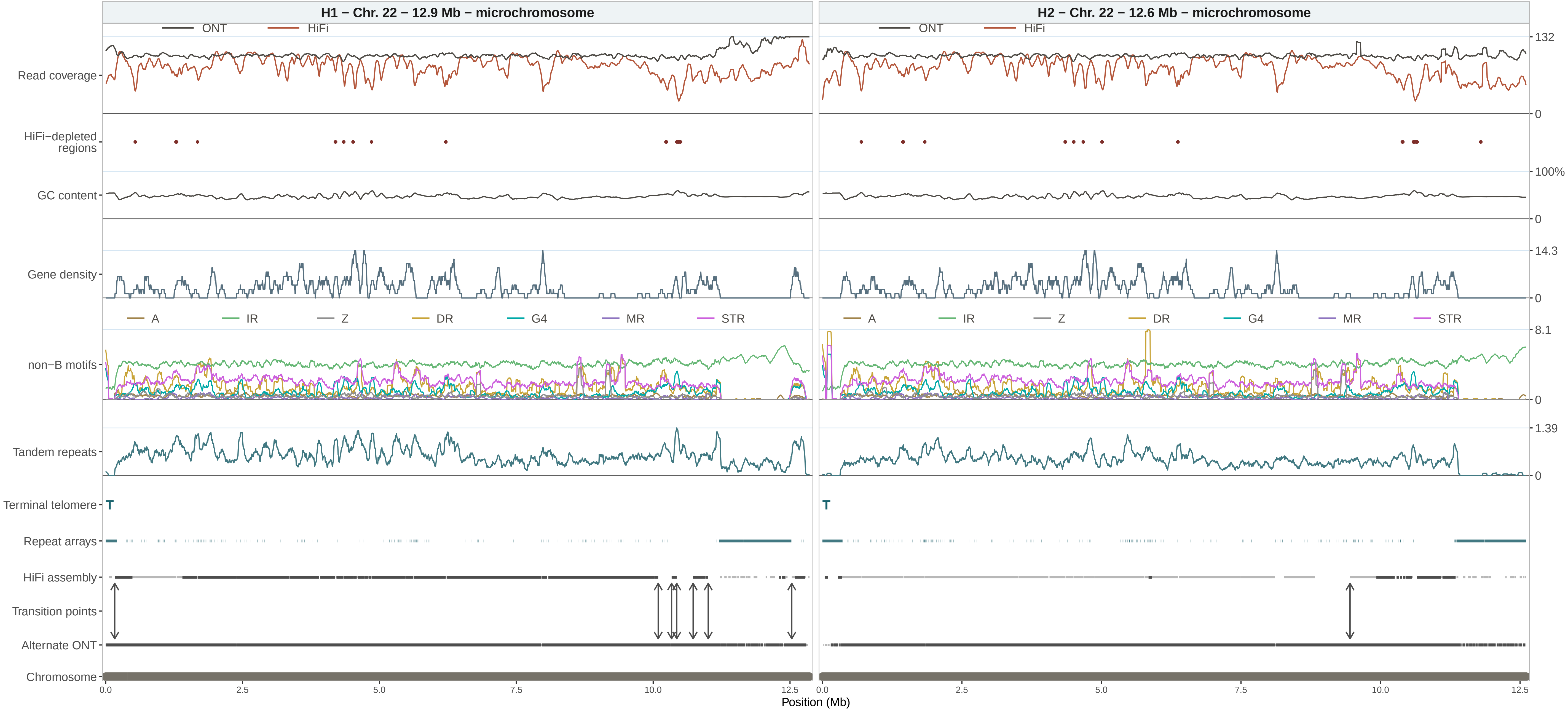

H1 − Chr. 22 − 12.9 Mb − microchromosome
H2 − Chr. 22 − 12.6 Mb − microchromosome
ONT
HiFi
Read coverage
HiFi−depleted regions
GC content
Gene density
non−B motifs
A
IR
Z
DR
G4
MR
STR
Tandem repeats
Terminal telomere
T
Repeat arrays
HiFi assembly
Transition points
Alternate ONT
Chromosome
132
0
100%
0
14.3
0
8.1
0
1.39
0
0.0
2.5
5.0
7.5
10.0
12.5
Position (Mb)

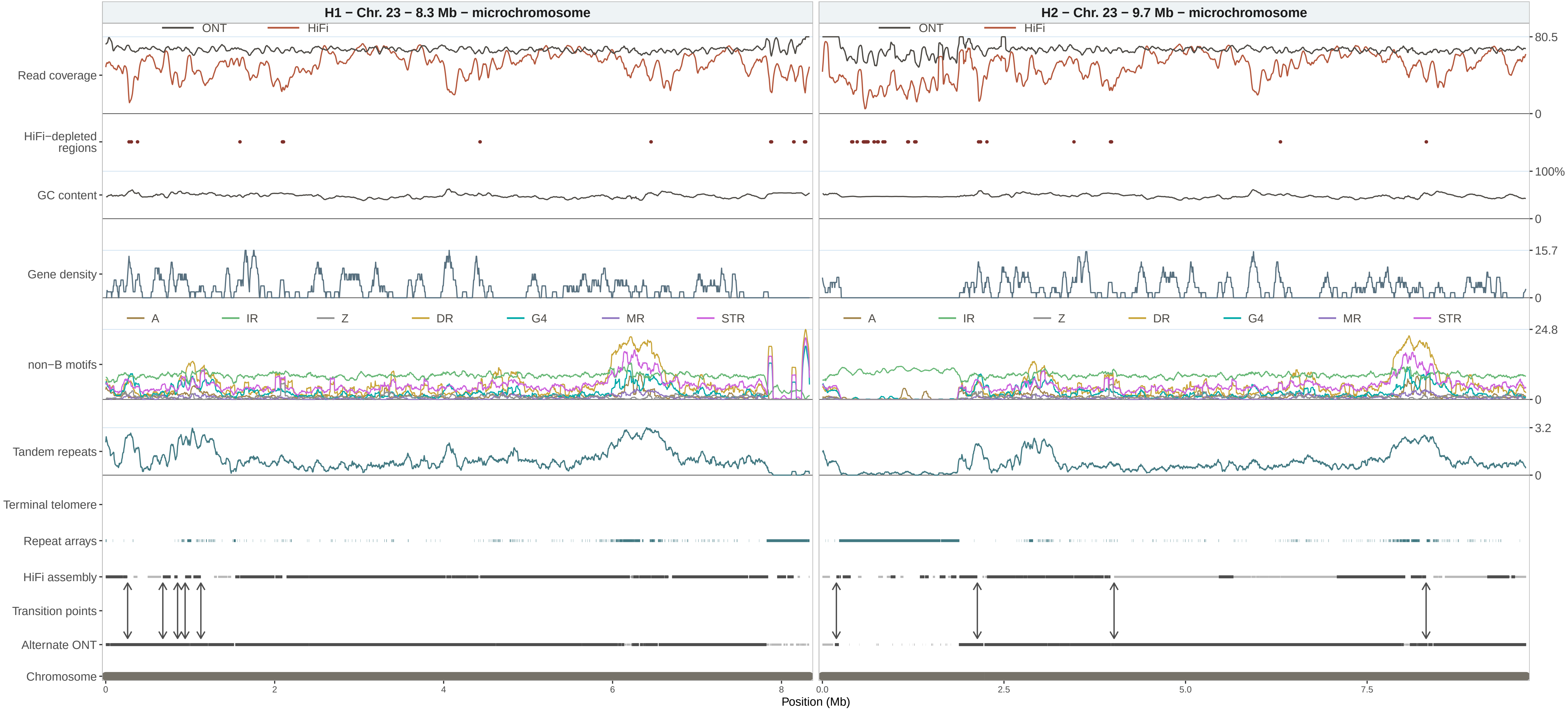

H1 − Chr. 23 − 8.3 Mb − microchromosome
H2 − Chr. 23 − 9.7 Mb − microchromosome
ONT
HiFi
Read coverage
HiFi−depleted regions
GC content
Gene density
non−B motifs
A
IR
Z
DR
G4
MR
STR
Tandem repeats
Terminal telomere
Repeat arrays
HiFi assembly
Transition points
Alternate ONT
Chromosome
80.5
0
100%
15.7
24.8
3.2
0
2
4
6
8
0.0
2.5
5.0
7.5
Position (Mb)

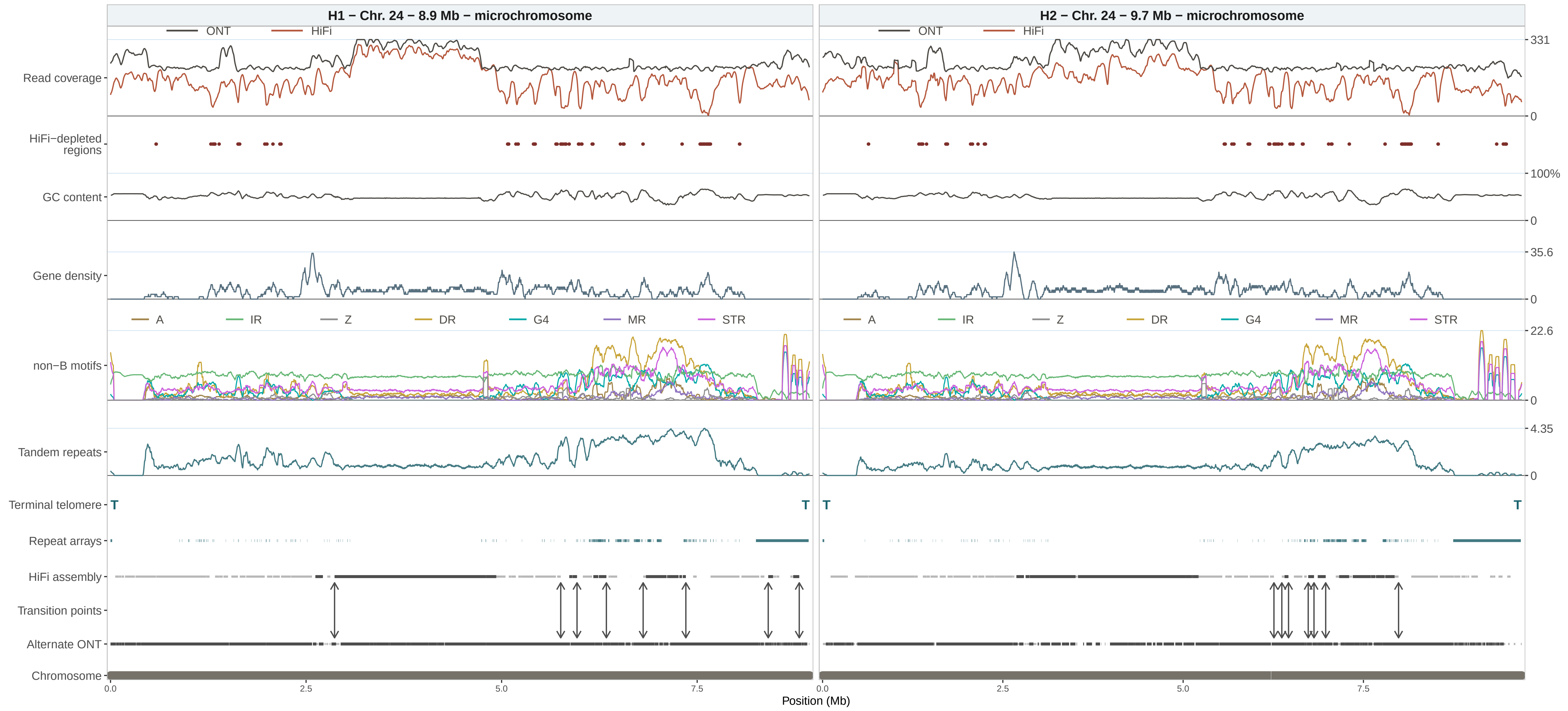
H1 − Chr. 24 − 8.9 Mb − microchromosome
H2 − Chr. 24 − 9.7 Mb − microchromosome
ONT
HiFi
Read coverage
HiFi−depleted regions
GC content
Gene density
non−B motifs
Tandem repeats
Terminal telomere
Repeat arrays
HiFi assembly
Transition points
Alternate ONT
Chromosome
A
IR
Z
DR
G4
MR
STR
T
331
0
100%
35.6
22.6
4.35
0.0
2.5
5.0
7.5
Position (Mb)

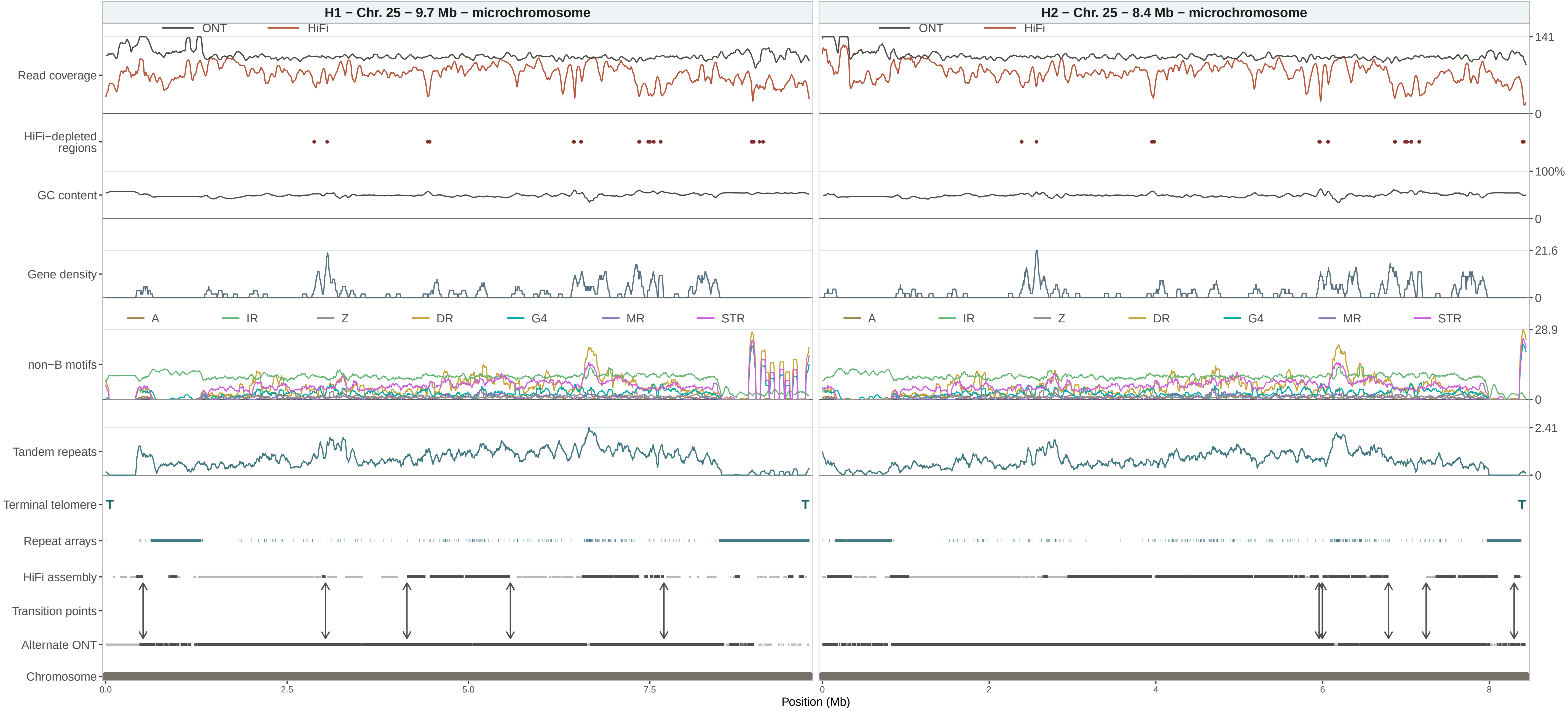

H1 − Chr. 25 − 9.7 Mb − microchromosome
H2 − Chr. 25 − 8.4 Mb − microchromosome
ONT
HiFi
Read coverage
HiFi−depleted regions
GC content
Gene density
non−B motifs
A
IR
Z
DR
G4
MR
STR
Tandem repeats
Terminal telomere
T
Repeat arrays
HiFi assembly
Transition points
Alternate ONT
Chromosome
141
0
100%
21.6
28.9
2.41
0.0
2.5
5.0
7.5
2
4
6
8
Position (Mb)

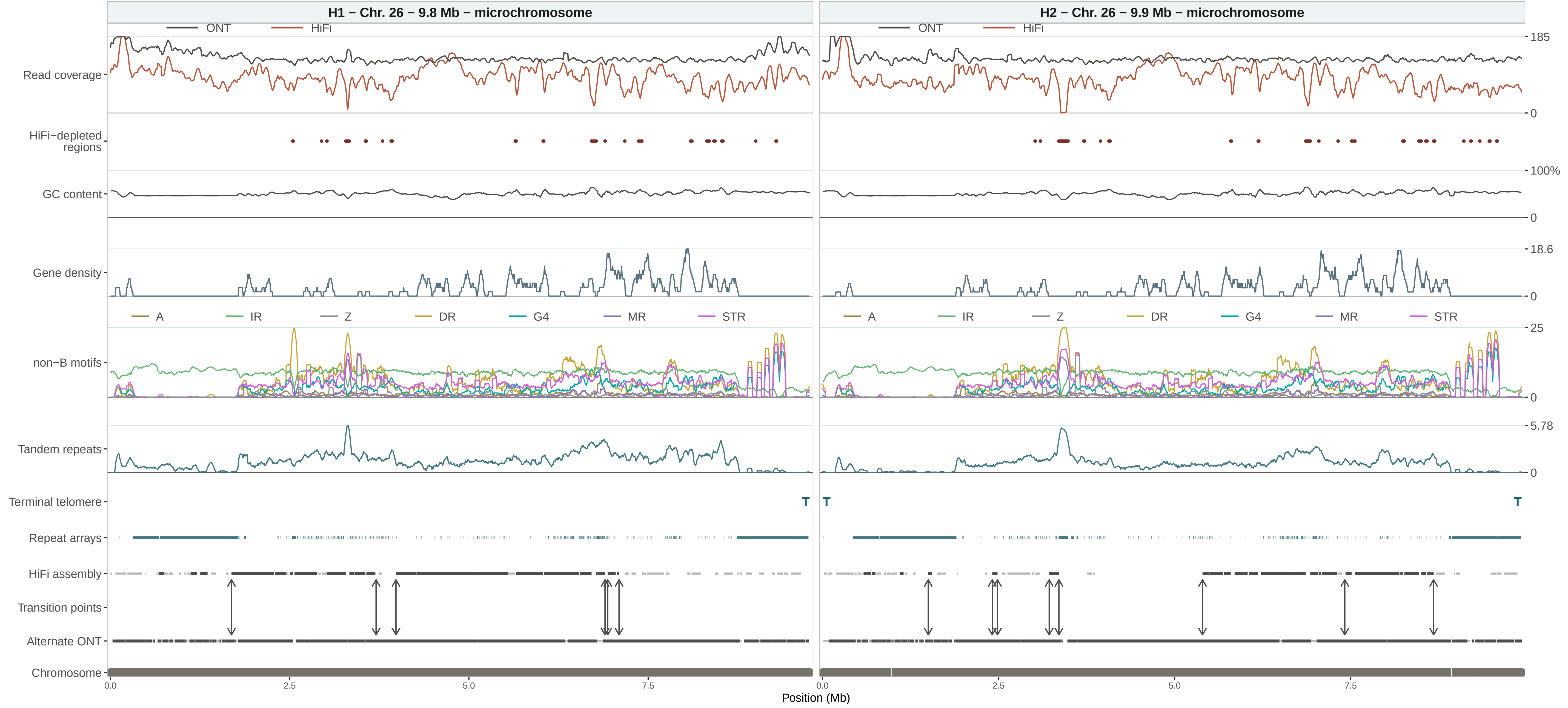
H1 − Chr. 26 − 9.8 Mb − microchromosome
H2 − Chr. 26 − 9.9 Mb − microchromosome
ONT
HiFi
Read coverage
HiFi−depleted regions
GC content
Gene density
non−B motifs
A
IR
Z
DR
G4
MR
STR
Tandem repeats
Terminal telomere
T
Repeat arrays
HiFi assembly
Transition points
Alternate ONT
Chromosome
185
0
100%
18.6
25
5.78
0.0
2.5
5.0
7.5
Position (Mb)

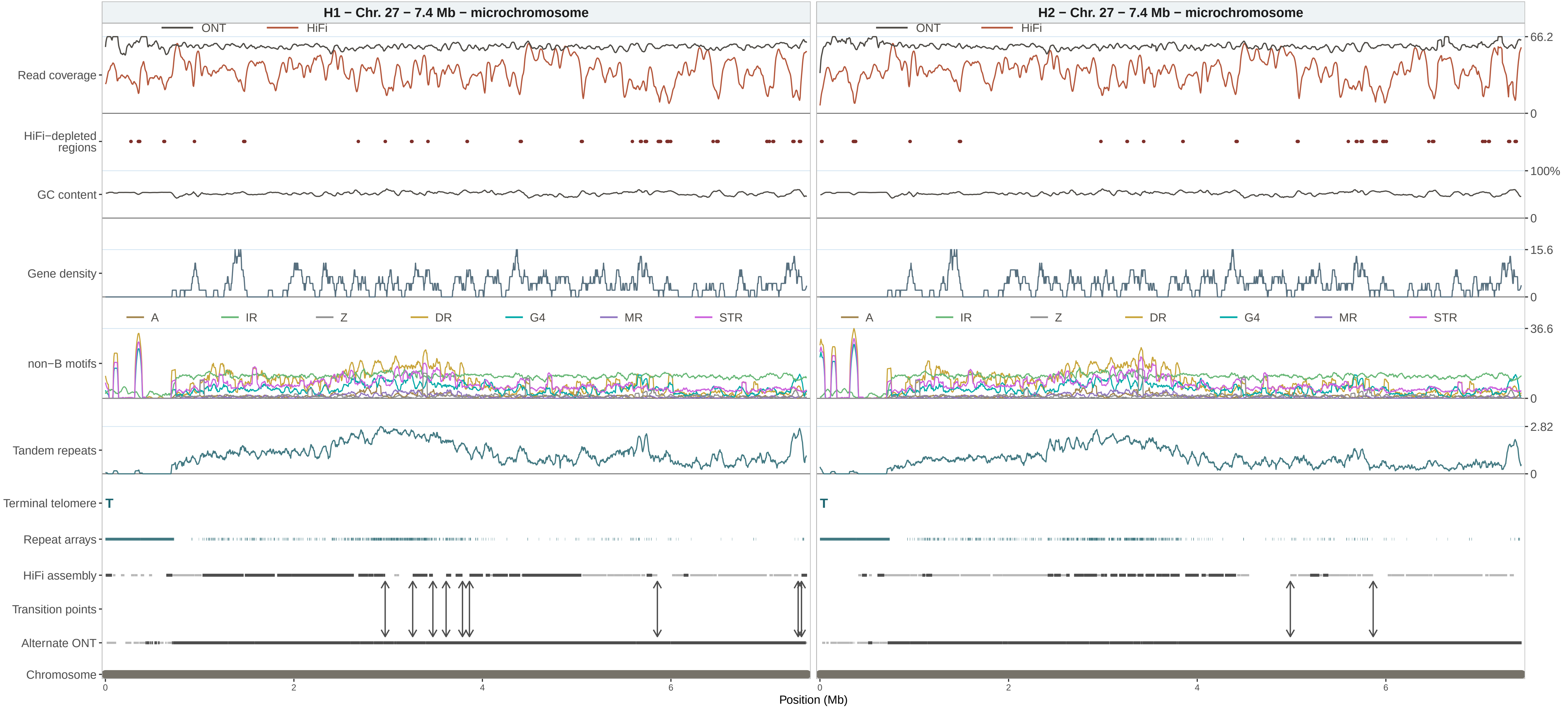

H1 − Chr. 27 − 7.4 Mb − microchromosome
H2 − Chr. 27 − 7.4 Mb − microchromosome
ONT
HiFi
Read coverage
HiFi−depleted regions
GC content
Gene density
non−B motifs
A
IR
Z
DR
G4
MR
STR
Tandem repeats
Terminal telomere
T
Repeat arrays
HiFi assembly
Transition points
Alternate ONT
Chromosome
66.2
0
100%
15.6
36.6
2.82
2
4
6
Position (Mb)

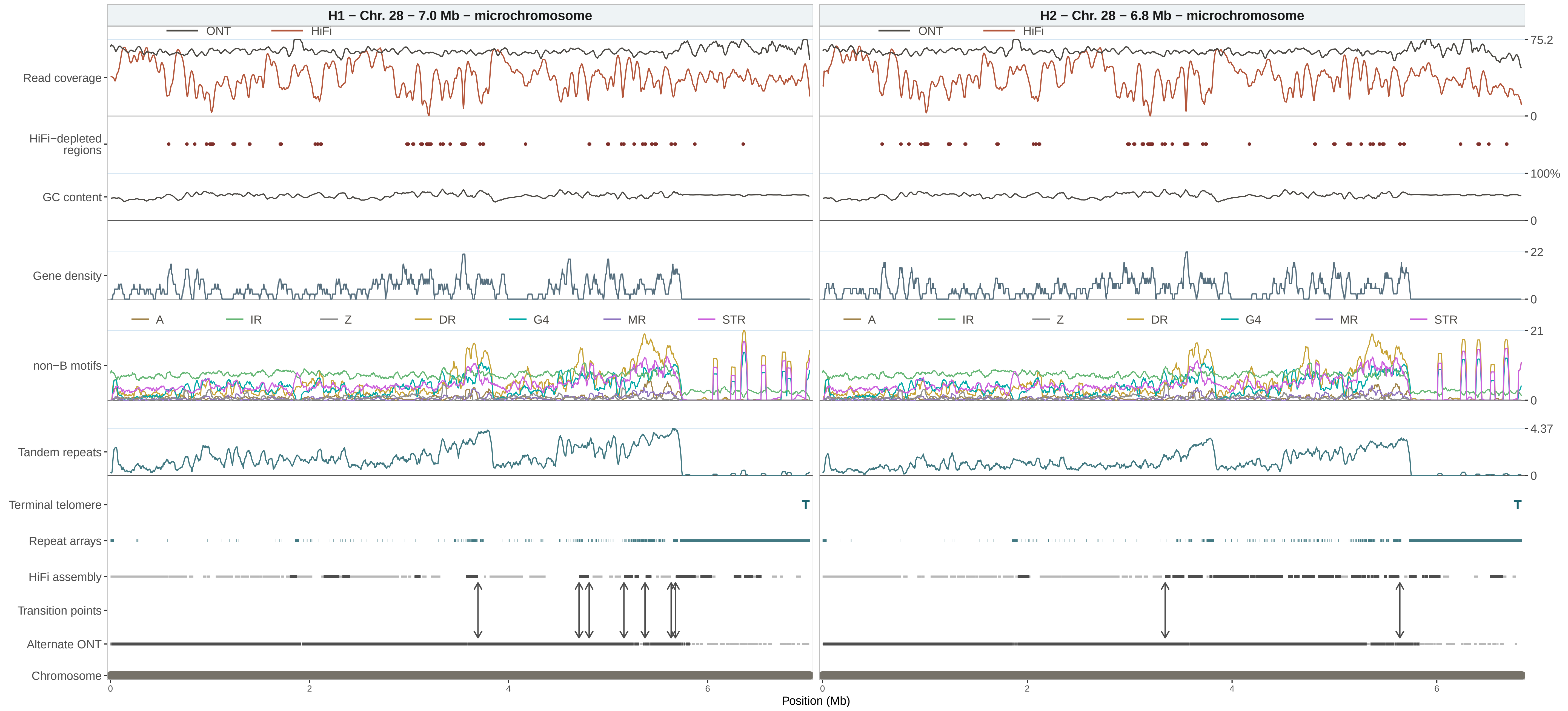
H1 − Chr. 28 − 7.0 Mb − microchromosome
H2 − Chr. 28 − 6.8 Mb − microchromosome
ONT
HiFi
Read coverage
HiFi−depleted regions
GC content
Gene density
non−B motifs
A
IR
Z
DR
G4
MR
STR
Tandem repeats
Terminal telomere
T
Repeat arrays
HiFi assembly
Transition points
Alternate ONT
Chromosome
75.2
0
100%
0
22
0
21
0
4.37
0
0
2
4
6
Position (Mb)

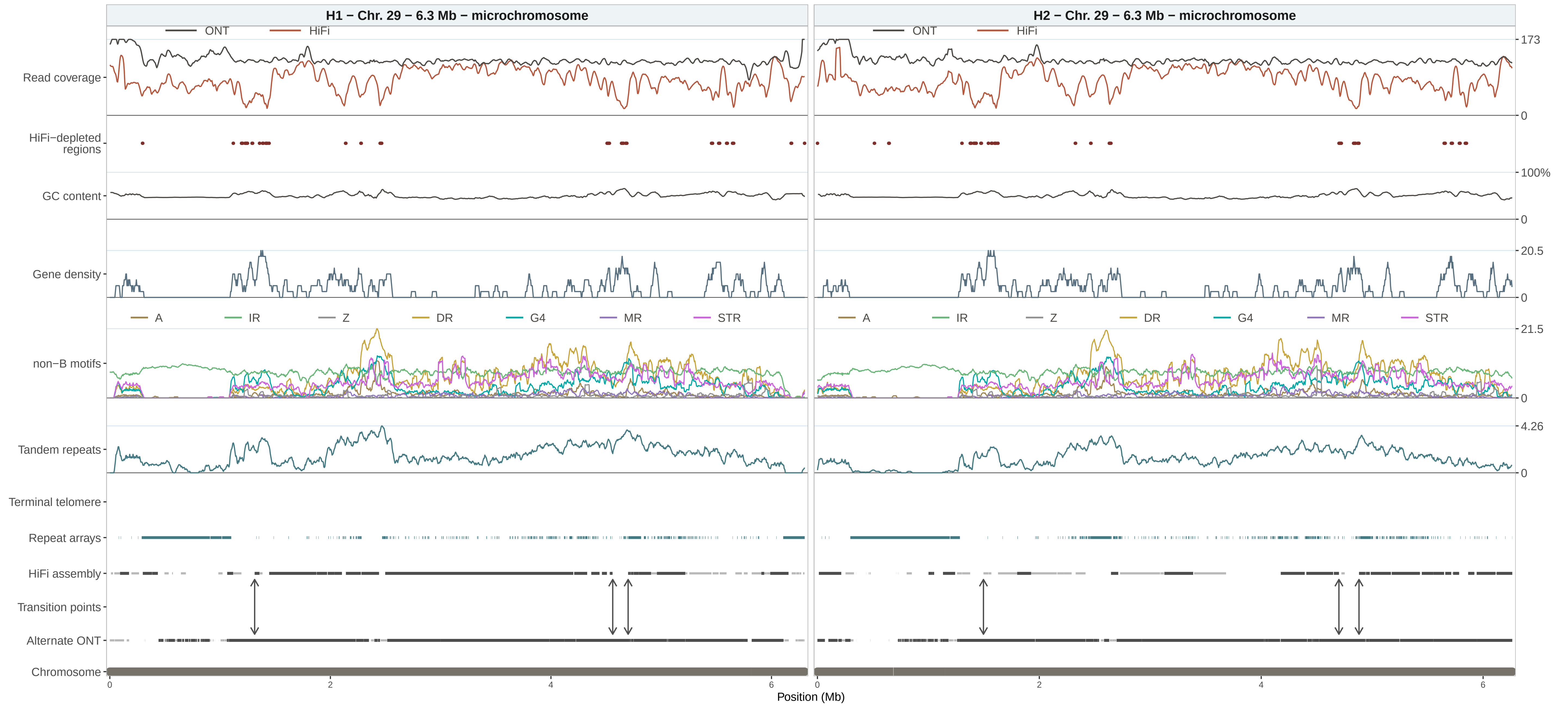

H1 − Chr. 29 − 6.3 Mb − microchromosome
H2 − Chr. 29 − 6.3 Mb − microchromosome
ONT
HiFi
Read coverage
HiFi−depleted regions
GC content
Gene density
non−B motifs
A
IR
Z
DR
G4
MR
STR
Tandem repeats
Terminal telomere
Repeat arrays
HiFi assembly
Transition points
Alternate ONT
Chromosome
173
0
100%
0
20.5
0
21.5
0
4.26
0
0
2
4
6
Position (Mb)

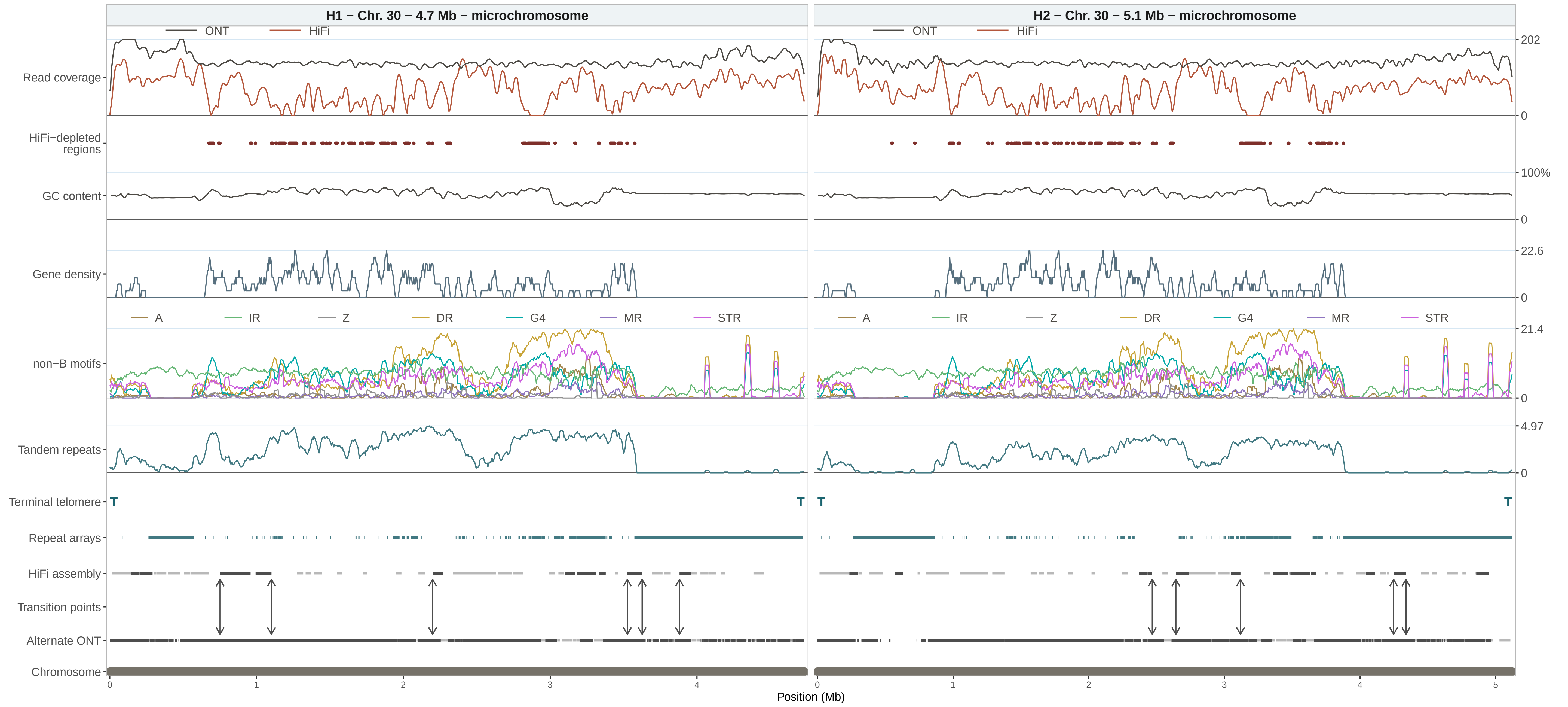

H1 − Chr. 30 − 4.7 Mb − microchromosome
H2 − Chr. 30 − 5.1 Mb − microchromosome
ONT
HiFi
Read coverage
HiFi−depleted regions
GC content
Gene density
non−B motifs
A
IR
Z
DR
G4
MR
STR
Tandem repeats
Terminal telomere
T
Repeat arrays
HiFi assembly
Transition points
Alternate ONT
Chromosome
202
0
100%
0
22.6
0
21.4
0
4.97
0
0
1
2
3
4
5
Position (Mb)

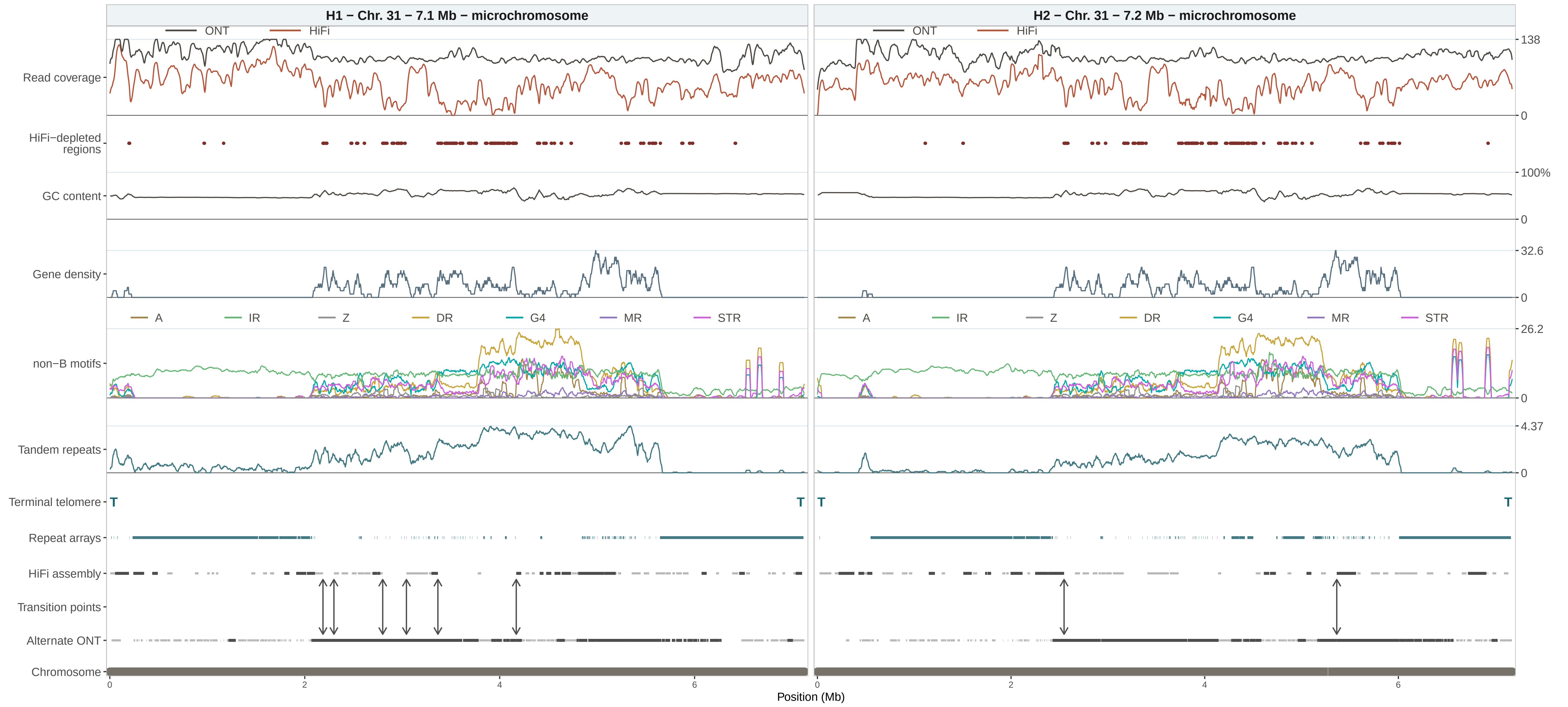

H1 − Chr. 31 − 7.1 Mb − microchromosome
H2 − Chr. 31 − 7.2 Mb − microchromosome
ONT
HiFi
Read coverage
HiFi−depleted regions
GC content
Gene density
non−B motifs
Tandem repeats
Terminal telomere
Repeat arrays
HiFi assembly
Transition points
Alternate ONT
Chromosome
A
IR
Z
DR
G4
MR
STR
T
138
0
100%
32.6
26.2
4.37
2
4
6
Position (Mb)

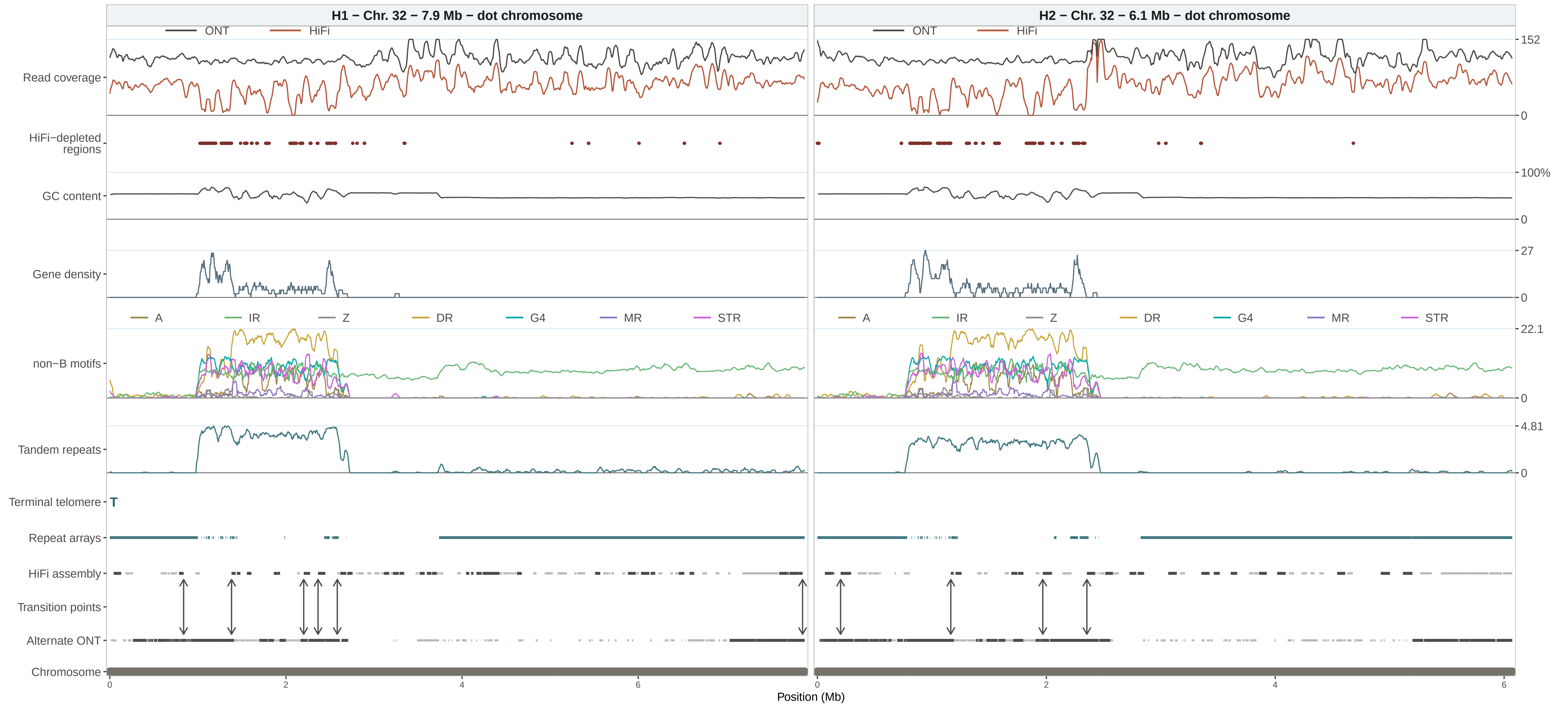

H1 − Chr. 32 − 7.9 Mb − dot chromosome
H2 − Chr. 32 − 6.1 Mb − dot chromosome
ONT
HiFi
Read coverage
HiFi−depleted regions
GC content
Gene density
non−B motifs
A
IR
Z
DR
G4
MR
STR
Tandem repeats
Terminal telomere
T
Repeat arrays
HiFi assembly
Transition points
Alternate ONT
Chromosome
152
0
100%
0
27
0
22.1
0
4.81
0
0
2
4
6
0
2
4
6
Position (Mb)

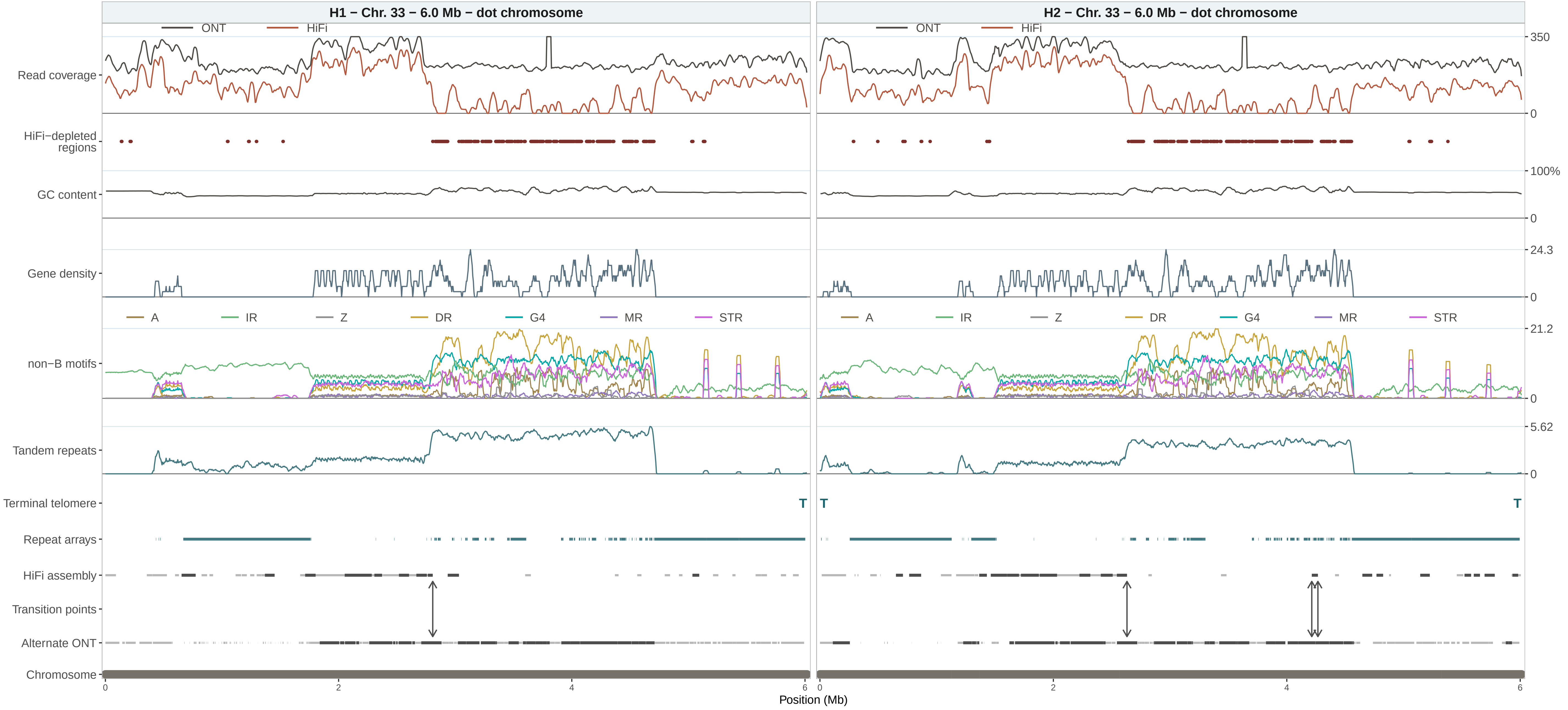

H1 − Chr. 33 − 6.0 Mb − dot chromosome
H2 − Chr. 33 − 6.0 Mb − dot chromosome
ONT
HiFi
Read coverage
HiFi−depleted regions
GC content
Gene density
non−B motifs
A
IR
Z
DR
G4
MR
STR
Tandem repeats
Terminal telomere
T
Repeat arrays
HiFi assembly
Transition points
Alternate ONT
Chromosome
350
0
100%
24.3
21.2
5.62
0
2
4
6
Position (Mb)

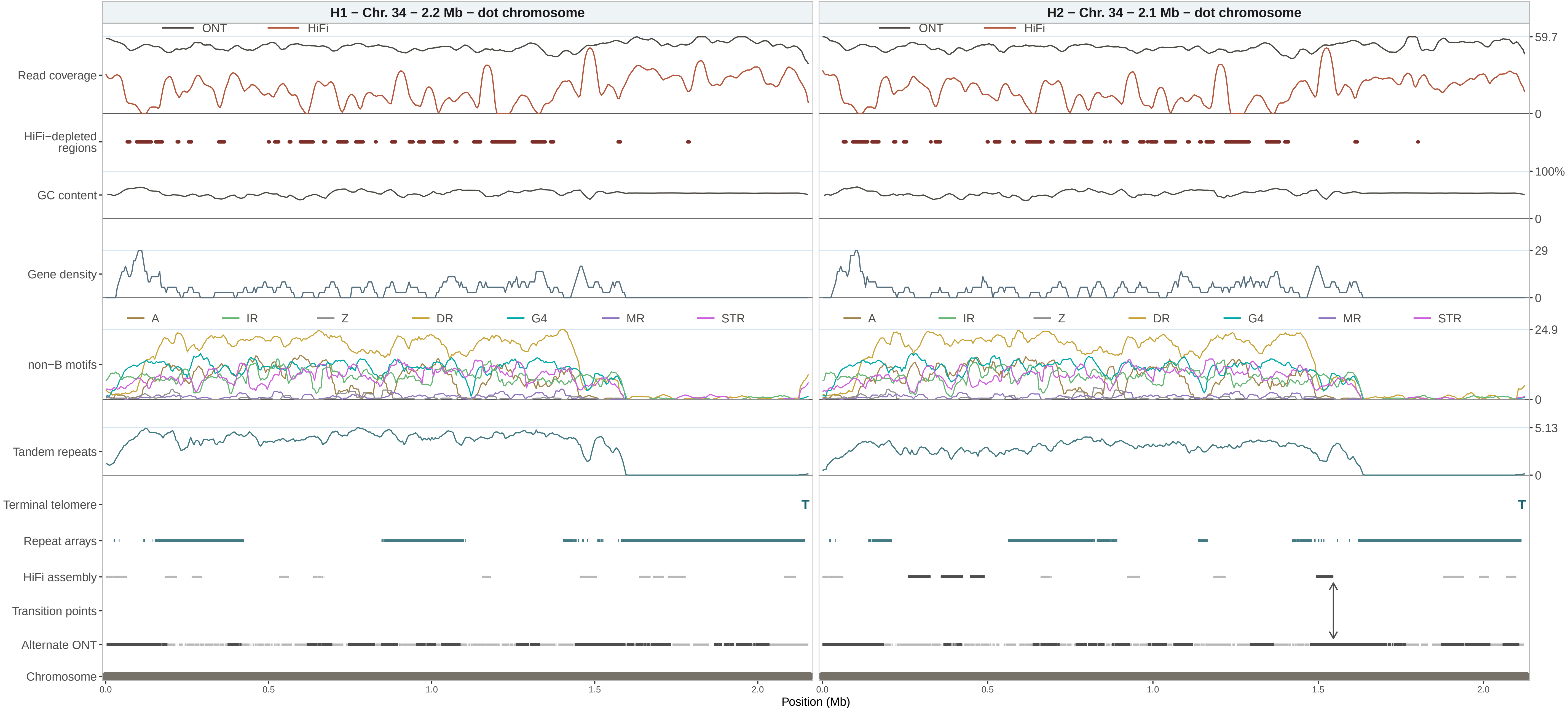
H1 − Chr. 34 − 2.2 Mb − dot chromosome
H2 − Chr. 34 − 2.1 Mb − dot chromosome
ONT
HiFi
59.7
0
Read coverage
HiFi−depleted regions
100%
GC content
29
Gene density
A
IR
Z
DR
G4
MR
STR
24.9
non−B motifs
5.13
Tandem repeats
Terminal telomere
T
Repeat arrays
HiFi assembly
Transition points
Alternate ONT
Chromosome
0.0
0.5
1.0
1.5
2.0
Position (Mb)

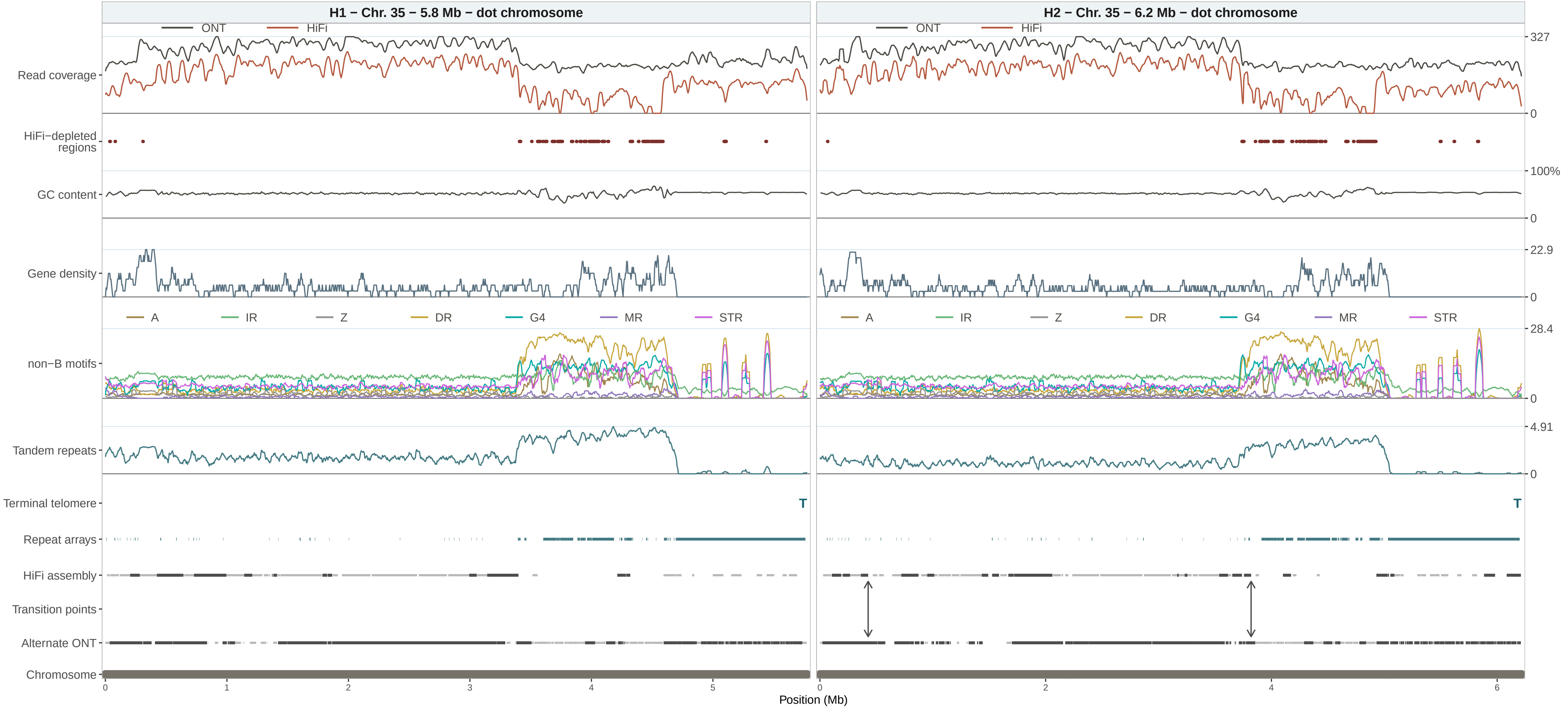

H1 − Chr. 35 − 5.8 Mb − dot chromosome
H2 − Chr. 35 − 6.2 Mb − dot chromosome
ONT
HiFi
Read coverage
HiFi−depleted regions
GC content
Gene density
non−B motifs
A
IR
Z
DR
G4
MR
STR
Tandem repeats
Terminal telomere
T
Repeat arrays
HiFi assembly
Transition points
Alternate ONT
Chromosome
327
0
100%
0
22.9
0
28.4
0
4.91
0
0
1
2
3
4
5
6
Position (Mb)

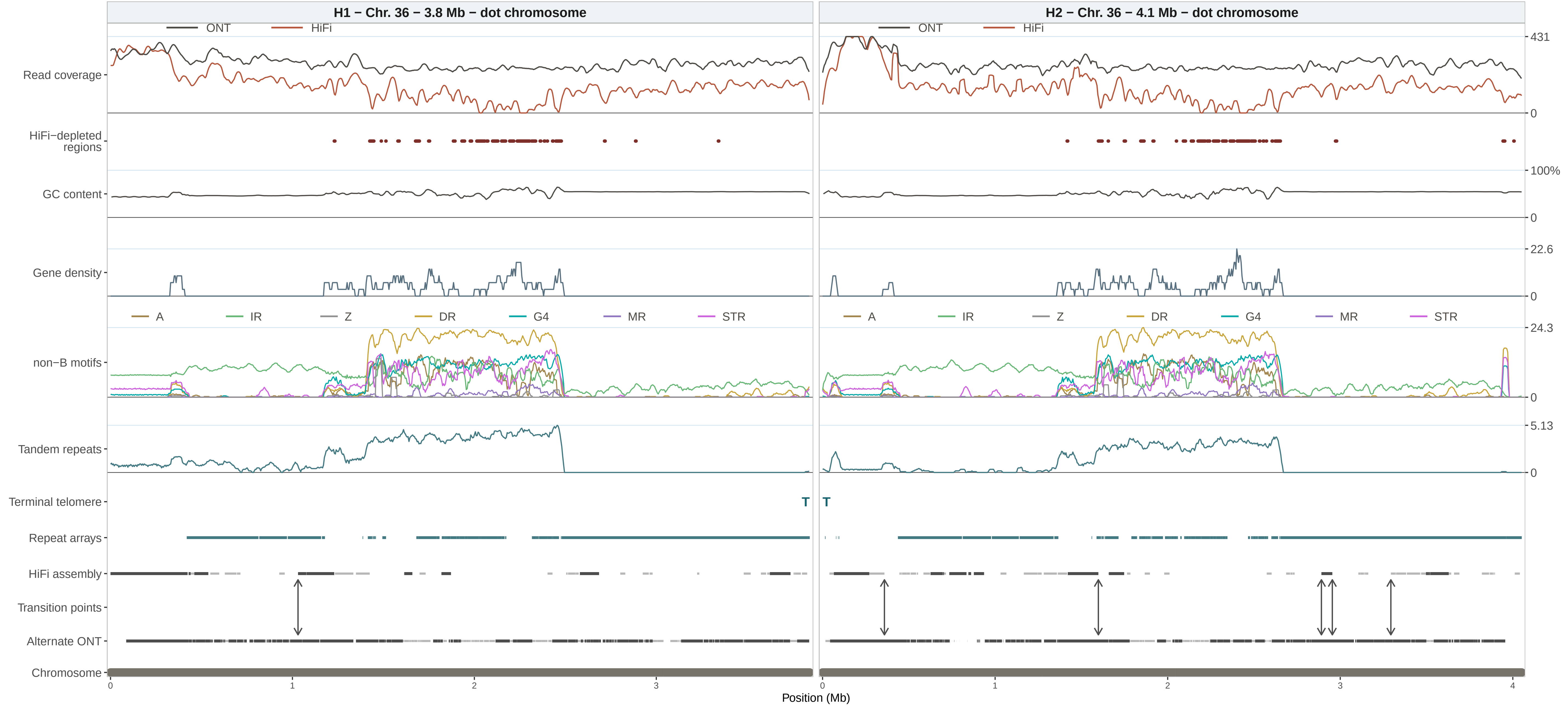

H1 − Chr. 36 − 3.8 Mb − dot chromosome
H2 − Chr. 36 − 4.1 Mb − dot chromosome
ONT
HiFi
Read coverage
HiFi−depleted regions
GC content
Gene density
non−B motifs
A
IR
Z
DR
G4
MR
STR
Tandem repeats
Terminal telomere
T
Repeat arrays
HiFi assembly
Transition points
Alternate ONT
Chromosome
431
0
100%
0
22.6
0
24.3
0
5.13
0
0
1
2
3
4
Position (Mb)

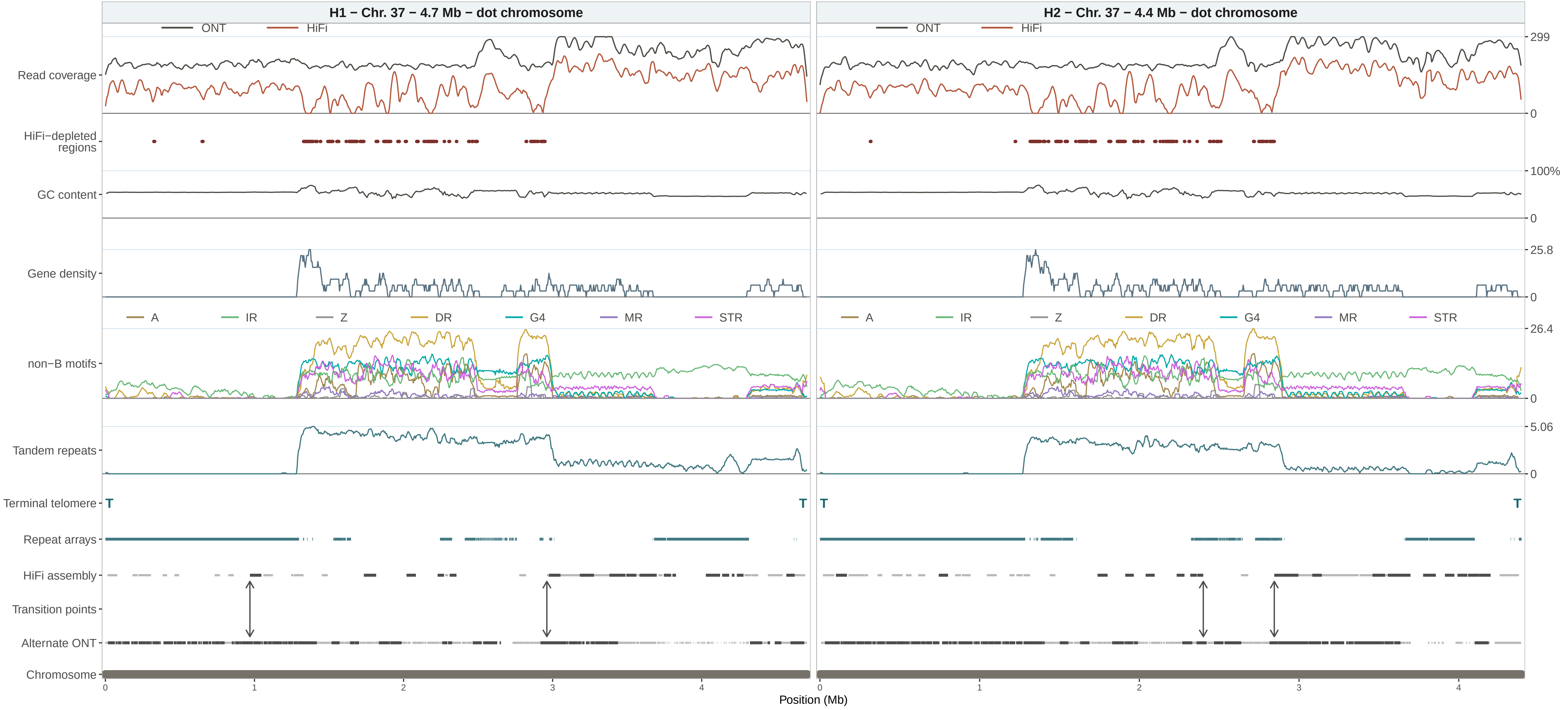
H1 − Chr. 37 − 4.7 Mb − dot chromosome
H2 − Chr. 37 − 4.4 Mb − dot chromosome
ONT
HiFi
Read coverage
HiFi−depleted regions
GC content
Gene density
non−B motifs
A
IR
Z
DR
G4
MR
STR
Tandem repeats
Terminal telomere
T
Repeat arrays
HiFi assembly
Transition points
Alternate ONT
Chromosome
299
0
100%
25.8
26.4
5.06
0
1
2
3
4
Position (Mb)

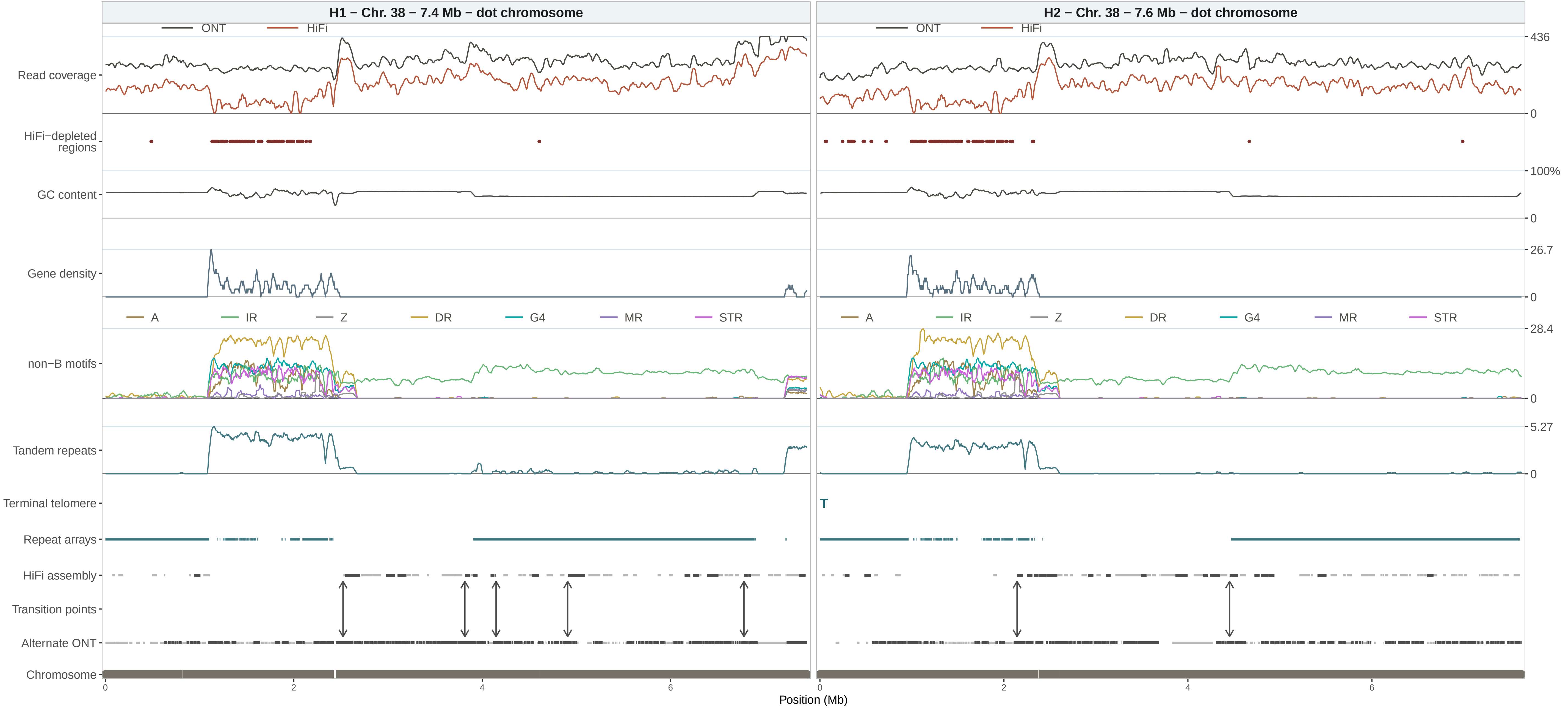

H1 − Chr. 38 − 7.4 Mb − dot chromosome
H2 − Chr. 38 − 7.6 Mb − dot chromosome
ONT
HiFi
Read coverage
HiFi−depleted regions
GC content
Gene density
non−B motifs
Tandem repeats
Terminal telomere
Repeat arrays
HiFi assembly
Transition points
Alternate ONT
Chromosome
A
IR
Z
DR
G4
MR
STR
T
436
0
100%
0
26.7
0
28.4
0
5.27
0
0
2
4
6
Position (Mb)

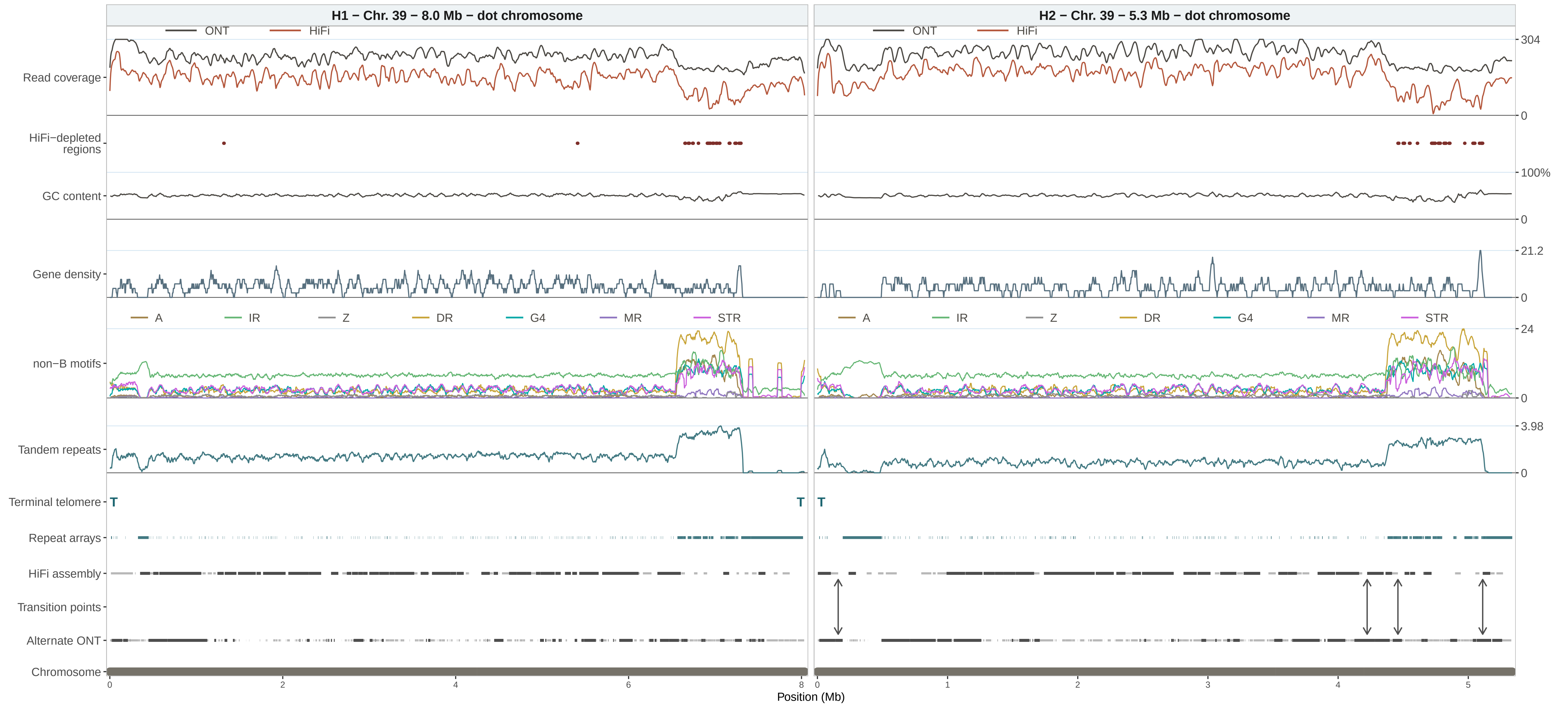
H1 − Chr. 39 − 8.0 Mb − dot chromosome
H2 − Chr. 39 − 5.3 Mb − dot chromosome
ONT
HiFi
Read coverage
HiFi−depleted regions
GC content
Gene density
A
IR
Z
DR
G4
MR
STR
non−B motifs
Tandem repeats
Terminal telomere
T
Repeat arrays
HiFi assembly
Transition points
Alternate ONT
Chromosome
304
0
100%
0
21.2
0
24
0
3.98
0
0
2
4
6
8
0
1
2
3
4
5
Position (Mb)

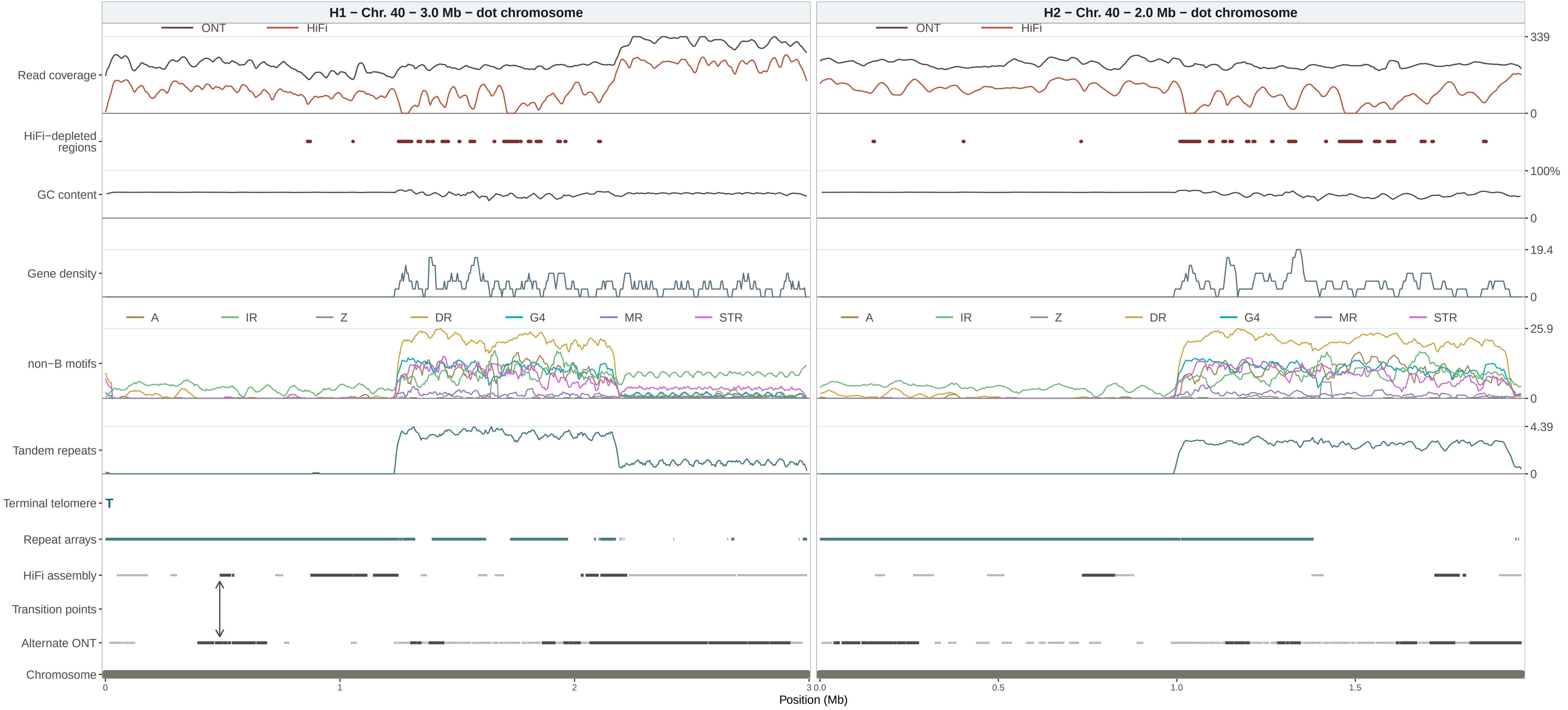

H1 − Chr. 40 − 3.0 Mb − dot chromosome
H2 − Chr. 40 − 2.0 Mb − dot chromosome
ONT
HiFi
Read coverage
HiFi−depleted regions
GC content
Gene density
non−B motifs
A
IR
Z
DR
G4
MR
STR
Tandem repeats
Terminal telomere
T
Repeat arrays
HiFi assembly
Transition points
Alternate ONT
Chromosome
339
0
100%
0
19.4
0
25.9
0
4.39
0
0
1
2
3
0.0
0.5
1.0
1.5
Position (Mb)

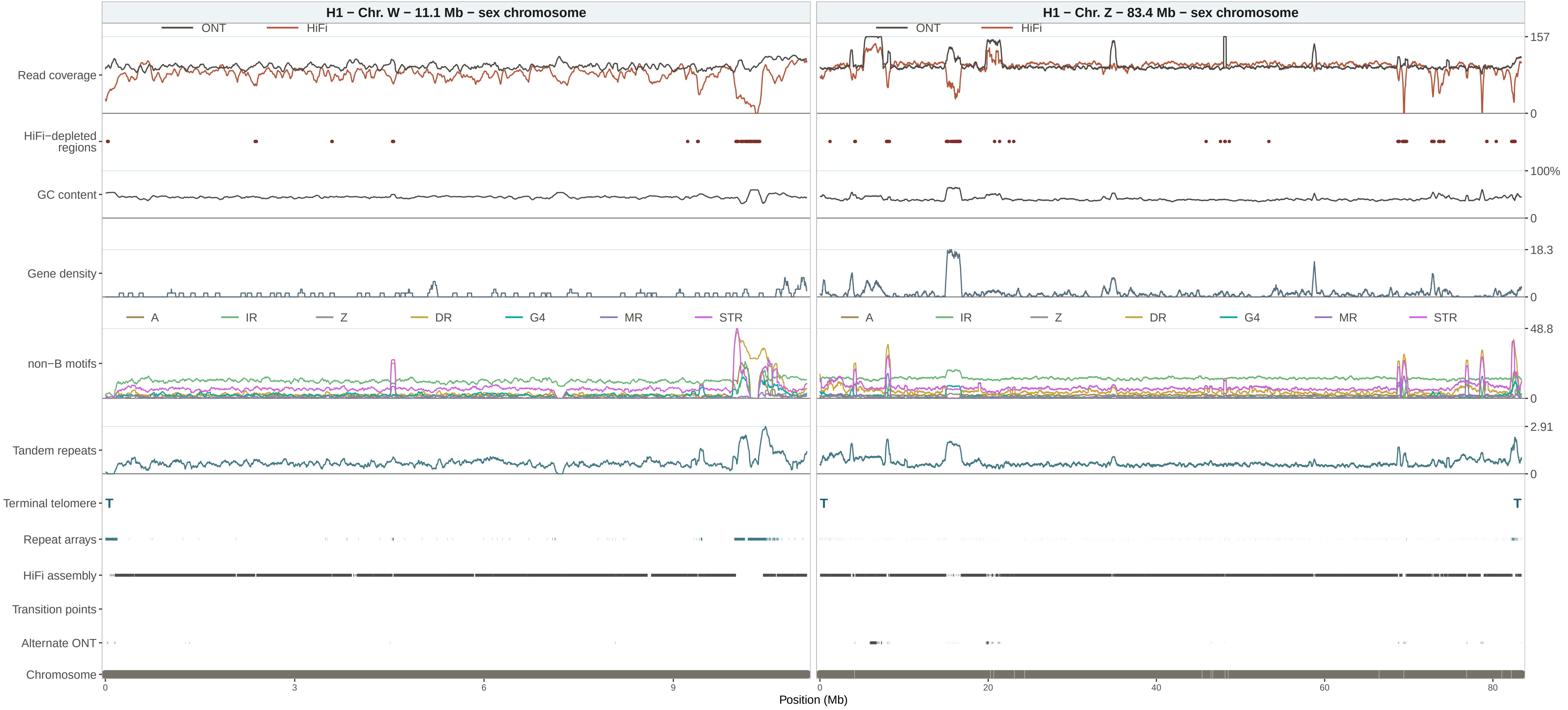
H1 – Chr. W – 11.1 Mb – sex chromosome
H1 – Chr. Z – 83.4 Mb – sex chromosome
ONT
HiFi
Read coverage
HiFi−depleted regions
GC content
Gene density
non−B motifs
A
IR
Z
DR
G4
MR
STR
Tandem repeats
Terminal telomere
T
Repeat arrays
HiFi assembly
Transition points
Alternate ONT
Chromosome
157
0
100%
18.3
48.8
2.91
0
3
6
9
20
40
60
80
Position (Mb)

**Supplementary Figure 4. Pairwise whole-chromosome comparison of HiFi read and assembly support against the ONT assemblies.** Hap1 and hap2 copies of each numbered chromosome are displayed side by side using their respective ONT assembly coordinates. Continuous tracks share a vertical scale within each chromosome pair, whereas genomic x-axes reflect the pseudo-haplotype-specific chromosome lengths. Tracks show ONT and HiFi read coverage, HiFi-depleted regions, GC content, gene density, predicted non-B-DNA motifs, ULTRA tandem repeats, terminal telomeres, TRASH2 repeat arrays, HiFi assembly support, alternate-ONT support, retained transition points, and chromosome span. HiFi-depleted regions are continuous intervals of exact HiFi zero coverage at least 1 kb long whose complete interval does not overlap ONT zero coverage. The "HiFi assembly" track pools support from both HiFi pseudo-haplotypes, whereas "Alternate ONT" shows support from the opposing ONT pseudo-haplotype. Pale grey marks show support from all alignments and dark grey marks high-confidence support from alignments at least 50 kb long with MAPQ at least 30. Double-headed arrows mark retained transition coordinates, and assembly N runs appear as white interruptions in the chromosome span when present. Right-hand values show the joint 99th-percentile coverage limit, 100% for GC, and observed maxima for the remaining quantitative tracks. Chromosomes 1–40 are presented in numerical order; W and Z, which occur only in H1, are displayed together on the final page. **Abbreviations** A: A-phased repeat, IR: inverted repeat, Z: Z-DNA motif, DR: direct repeat, G4: G-quadruplex motif, MR: mirror repeat, STR: short tandem repeat.
